# Alterations in Lysosomal, Glial and Neurodegenerative Biomarkers in Patients with Sporadic and Genetic Forms of Frontotemporal Dementia

**DOI:** 10.1101/2024.02.09.579529

**Authors:** Jennifer Hsiao-Nakamoto, Chi-Lu Chiu, Lawren VandeVrede, Ritesh Ravi, Brittany Vandenberg, Jack De Groot, Buyankhishig Tsogtbaatar, Meng Fang, Paul Auger, Neal S. Gould, Filippo Marchioni, Casey A. Powers, Sonnet S. Davis, Jung H. Suh, Jamal Alkabsh, Hilary W. Heuer, Argentina Lario Lago, Kimberly Scearce-Levie, William W. Seeley, Bradley F. Boeve, Howard J. Rosen, Amy Berger, Richard Tsai, Gilbert Di Paolo, Adam L. Boxer, Akhil Bhalla, Fen Huang, ALLFTD Consortium

## Abstract

**Background:** Frontotemporal dementia (FTD) is the most common cause of early-onset dementia with 10-20% of cases caused by mutations in one of three genes: *GRN*, *C9orf72*, or *MAPT*. To effectively develop therapeutics for FTD, the identification and characterization of biomarkers to understand disease pathogenesis and evaluate the impact of specific therapeutic strategies on the target biology as well as the underlying disease pathology are essential. Moreover, tracking the longitudinal changes of these biomarkers throughout disease progression is crucial to discern their correlation with clinical manifestations for potential prognostic usage.

**Methods:** We conducted a comprehensive investigation of biomarkers indicative of lysosomal biology, glial cell activation, synaptic and neuronal health in cerebrospinal fluid (CSF) and plasma from non-carrier controls, sporadic FTD (symptomatic non-carriers) and symptomatic carriers of mutations in *GRN, C9orf72,* or *MAPT*, as well as asymptomatic *GRN* mutation carriers. We also assessed the longitudinal changes of biomarkers in *GRN* mutation carriers. Furthermore, we examined biomarker levels in disease impacted brain regions including middle temporal gyrus (MTG) and superior frontal gyrus (SFG) and disease-unaffected inferior occipital gyrus (IOG) from sporadic FTD and symptomatic *GRN* carriers.

**Results:** We confirmed glucosylsphingosine (GlcSph), a lysosomal biomarker regulated by progranulin, was elevated in the plasma from *GRN* mutation carriers, both symptomatic and asymptomatic. GlcSph and other lysosomal biomarkers such as ganglioside GM2 and globoside GB3 were increased in the disease affected SFG and MTG regions from sporadic FTD and symptomatic *GRN* mutation carriers, but not in the IOG, compared to the same brain regions from controls. The glial biomarkers GFAP in plasma and YKL40 in CSF were elevated in asymptomatic *GRN* carriers, and all symptomatic groups, except the symptomatic *C9orf72* mutation group. YKL40 was also increased in SFG and MTG regions from sporadic FTD and symptomatic *GRN* mutation carriers. Neuronal injury and degeneration biomarkers NfL in CSF and plasma, and UCHL1 in CSF were elevated in patients with all forms of FTD. Synaptic biomarkers NPTXR, NPTX1/2, and VGF were reduced in CSF from patients with all forms of FTD, with the most pronounced reductions observed in symptomatic *MAPT* mutation carriers. Furthermore, we demonstrated plasma NfL was significantly positively correlated with disease severity as measured by CDR+NACC FTLD□SB in genetic forms of FTD and CSF NPTXR was significantly negatively correlated with CDR+NACC FTLD□SB in symptomatic *GRN* and *MAPT* mutation carriers.

**Conclusions:** In conclusion, our comprehensive investigation replicated alterations in biofluid biomarkers indicative of lysosomal function, glial activation, synaptic and neuronal health across sporadic and genetic forms of FTD and unveiled novel insights into the dysregulation of these biomarkers within brain tissues from patients with *GRN* mutations. The observed correlations between biomarkers and disease severity open promising avenues for prognostic applications and for indicators of drug efficacy in clinical trials. Our data also implicated a complicated relationship between biofluid and tissue biomarker changes and future investigations should delve into the mechanistic underpinnings of these biomarkers, which will serve as a foundation for the development of targeted therapeutics for FTD.

## Introduction

Frontotemporal dementia (FTD) is the leading cause of younger-onset dementia, frequently occurring before 65 years of age [^1^, ^2^]. FTD results in an impairment of executive function together with changes in personality and social behaviors, lapses in judgment, loss of insight, cognitive deficits, and sometimes language and/or motor dysfunction [^3^]. FTD is highly heritable, with 15–30% of cases caused by mutations in one of three genes: *GRN* (progranulin), *C9orf72* (chromosome 9 open reading frame 72), and *MAPT* (microtubule-associated protein tau) (Greaves, Rohrer, 2019^4^). The underlying pathology is termed frontotemporal lobar degeneration (FTLD) and consists of distinct underlying pathologies including aggregation of transactive response DNA-binding protein 43 (TDP-43), tau and fused-in-sarcoma (FUS) [^2^].

Heterozygous loss-of-function mutations in *GRN* typically result in progranulin (PGRN) levels approximately 50% of normal in FTD causing mutation carriers [^5^, ^6^, ^7^]. Hallmark pathological features in symptomatic *GRN* mutation carriers include central nervous system (CNS) lipofuscinosis, gliosis, and TDP-43 pathology [^8^, ^7^, ^9^, ^10^, ^11^]. In patients with FTD caused by hexanucleotide repeat expansions (HREs) in the *C9orf72* gene, hallmark pathology includes dipeptide repeat proteins, RNA foci, and TDP-43 pathology. Patients with *MAPT* mutations typically have tau protein aggregation [^12^; ^3^]. Approximately half of patients with sporadic behavioral variant FTD exhibit TDP-43 pathology (FTLD-TDP), with the other half mostly caused by tau pathology (FTLD-Tau) [^13^].

Lysosomal dysfunction is increasingly recognized as a critical player in the pathogenesis of various neurodegenerative diseases [^14^, ^15^, ^16^]. At a genetic level, mutations in multiple genes related to endolysosomal function have been linked to FTD, including *C9orf72, GRN, TARDBP, VCP, CHMP2B,* and *SQSTM1* [^17^]. Multiple lines of evidence suggest that different forms of tau or tau aggregates are degraded in part through the lysosomal-autophagy pathway and *MAPT* mutations could lead to lysosomal stress and functional dysregulation [^14^; ^18^]. Growing evidence positions PGRN as a key modulator of lysosomal function [^15^, ^19^]. Specifically, PGRN regulates the enzymatic activities of lysosomal hydrolases, such as cathepsin D, glucocerebrosidase (GCase), and lysosomal phospholipid bis(monoacylglycero)phosphate (BMP), which influences the metabolism of major GCase substrates, such as glucosylsphingosine (GlcSph), and other lysosomal hydrolases [^19^, ^20^, ^21^, ^15^]. Complete loss of PGRN caused by homozygous loss of function mutations results in the severe early onset lysosomal storage disorder, neuronal ceroid lipofuscinosis type 11 [^22^, ^15^].

Progressive neurodegeneration is a hallmark of FTD [^23^, ^24^]. A specific population of neurons in the frontal and temporal regions are particularly vulnerable to degeneration. This results in progressive changes in behavior, personality, and language in patients with FTD. Significant alterations in glial cells, which are critical for brain homeostasis and immune surveillance, have also been shown to be altered in patients with sporadic and genetic forms of FTD [^24^, ^25^].

To date, there are no approved treatments that can slow or reverse FTD progression. Given the heterogeneity of neuropathology in patients with sporadic forms of FTD, drug development efforts have mostly focused on FTD associated with *GRN, C9orf72,* or *MAPT* mutations. Clinical trials for patients with *GRN* mutation have been of particular interest based on the potential benefit by restoring normal PGRN levels. Therapeutic strategies aim to restore PGRN levels in the brain, including gene delivery using adeno-associated virus (AAV) vectors, modulation of extracellular PGRN levels using monoclonal antibodies directed against cell surface receptor sortilin, and replacement therapy using a recombinant PGRN that has been engineered to cross the blood-brain barrier [^2^]. To inform therapeutic development for FTD, it is essential to identify biomarkers that may be used to assess drug treatment effects and to select patients that may best respond to certain therapies.

## Materials and methods

### CSF and plasma samples

De-identified CSF & plasma samples from patients with sporadic FTD (symptomatic non-carriers, S-NC, n=47), symptomatic mutation carriers in *GRN* (S-*GRN*, n=31)*, C9orf72* (S-*C9orf72,* n=22) *or MAPT* (S-*MAPT*, n=25), asymptomatic *GRN* carriers (AS-*GRN*, n=34) and asymptomatic non-carrier controls (AS-NC, n=44) collected through the ARTFL LEFFTDS Longitudinal Frontotemporal Lobar Degeneration (ALLFTD) research network (NCT04363684), were obtained from National Centralized Repository for Alzheimer’s Disease and Related Dementias (NCRAD) and the University of California, San Francisco (UCSF). Clinical diagnoses were determined by an experienced behavioral neurologist using all available information at time of evaluation [^26^, ^27^, ^28^, ^29^]. The demographic and clinical phenotype characteristics are provided in Table 1.

**Table 1:**
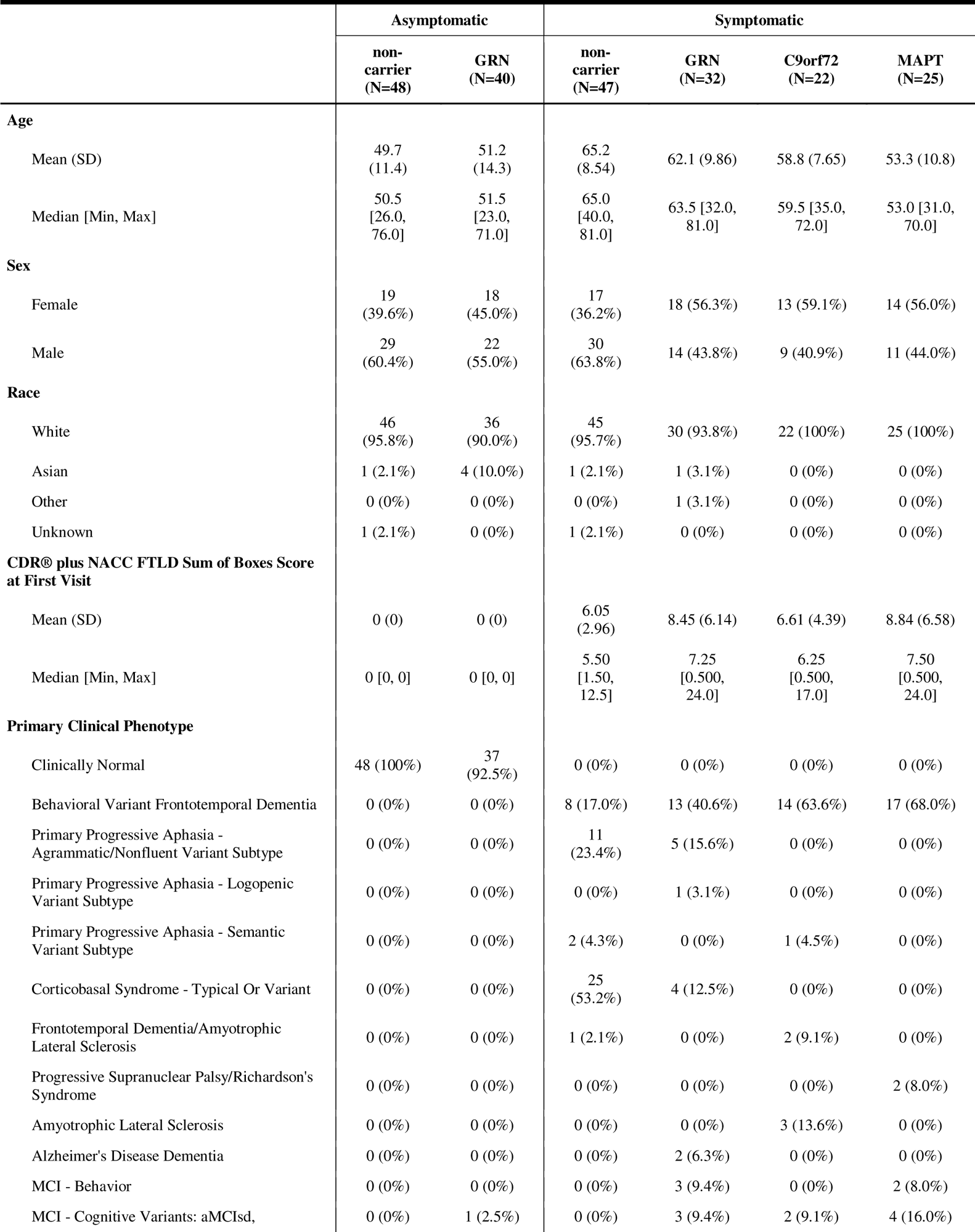
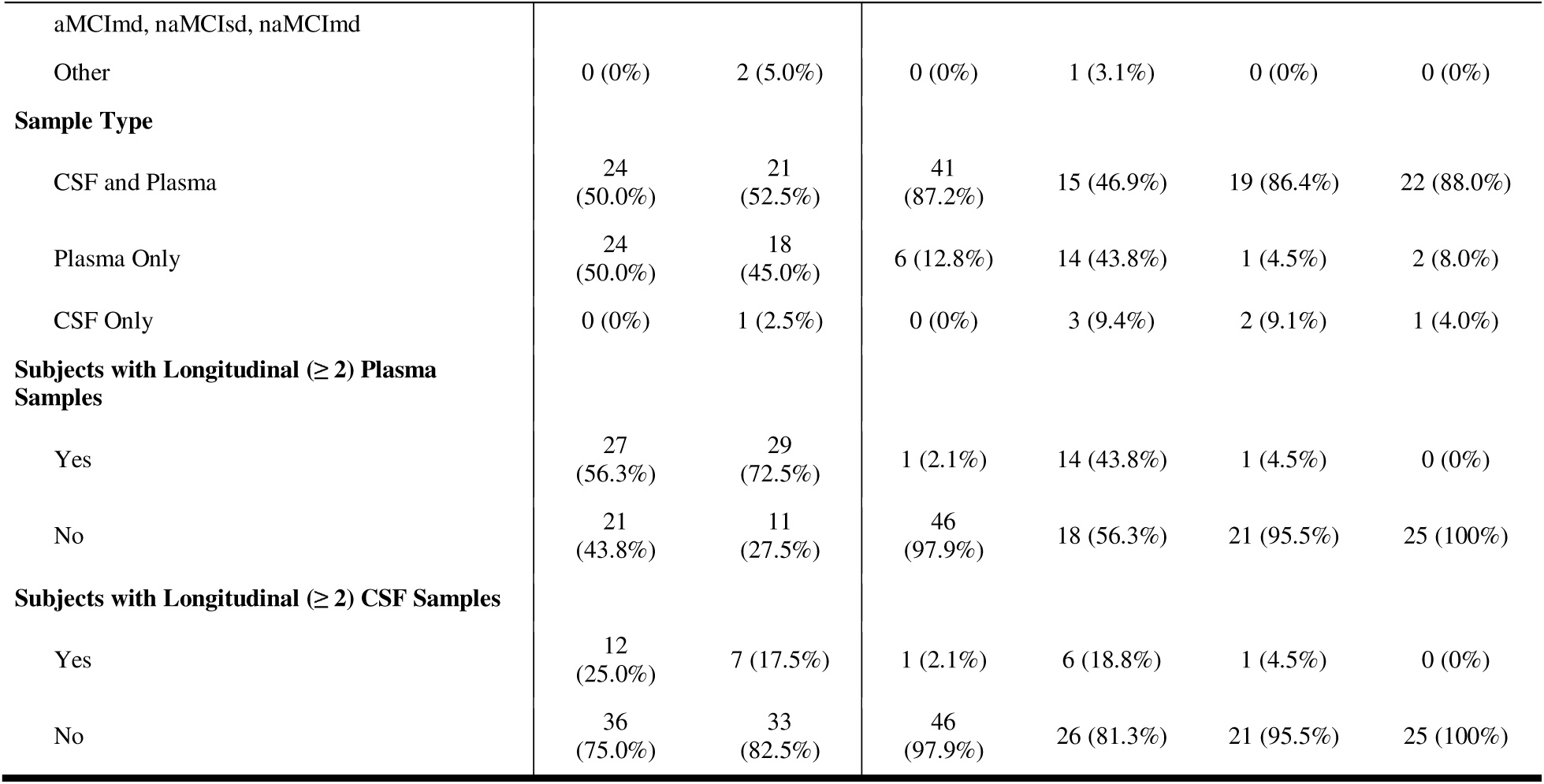
Demographics and clinical characteristics for biofluid samples.

### Human brain tissues

Brain samples from patients with sporadic FTD (S-NC, n=4) or symptomatic *GRN* mutation carriers (S-*GRN*, n=10) were obtained from Neurodegenerative Disease Brain Bank (NDBB) at UCSF. All patients had FTLD-TDP type A pathology. Authorization for autopsy was provided by the patients’ next of kin, and procedures were approved by the UCSF Committee on Human Research. Neuropathological diagnoses were made following consensus diagnostic criteria [^30^] and all selected cases have TDP-A pathology. Tissues from three brain regions were included in this study: generally affected degenerating regions in FTD, the middle temporal gyrus (MTG) and superior frontal gyrus (SFG) and region usually spared in FTD, the inferior occipital gyrus (IOG). Corresponding brain sections from age and sex matched non-carrier controls (n=15) were purchased from BioIVT. A demographic summary is provided in Table 2.

**Table 2:** Demographics and clinical diagnosis for brain tissues.

|  | Asymptomatic Control | Symptomatic |  |
| --- | --- | --- | --- |
|  | non-carrier (N=15) | non-carrier (N=4) | GRN (N=10) |
| **Age at Death** |  |  |  |
| Mean (SD) | 68.5 (5.85) | 69.3 (6.50) | 67.3 (6.65) |
| Median [Min, Max] | 70.0 [58.0, 76.0] | 68.0 [63.0, 78.0] | 67.0 [56.0, 78.0] |
| **Sex** |  |  |  |
| Female | 6 (40.0%) | 2 (50.0%) | 8 (80.0%) |
| Male | 9 (60.0%) | 2 (50.0%) | 2 (20.0%) |
| **Clinical Diagnosis** |  |  |  |
| Clinical Normal | 15 (100%) | 0 (0%) | 0 (0%) |
| Behavioral Variant Frontotemporal Dementia | 0 (0%) | 1 (25.0%) | 6 (60.0%) |
| Mixed Frontotemporal Dementia | 0 (0%) | 0 (0%) | 1 (10.0%) |
| Primary Progressive Aphasia, Unspecified | 0 (0%) | 0 (0%) | 1 (10.0%) |
| Corticobasal Syndrome | 0 (0%) | 2 (50.0%) | 1 (10.0%) |
| Progressive Non-Fluent Aphasia | 0 (0%) | 1 (25.0%) | 0 (0%) |
| Corticobasal syndrome/Progressive Non-Fluent Aphasia | 0 (0%) | 0 (0%) | 1 (10.0%) |

### Brain tissue homogenization for protein assays

Weighed brain tissue samples were processed for protein assays by addition of 10X volume NP-40 buffer with added cOmplete Protease Inhibitor (Roche #04693132001) and PhosStop (Roche 04906837001) phosphatase inhibitors. Samples were then homogenized using a 3mm tungsten carbide bead, shaken using the Qiagen TissueLyzer II (Cat No./ID: 85300) (2×3 minutes at 30 Hz). Samples were centrifuged at 14,000g for 20 min at 4°C, then supernatant was aliquoted for analysis. BCA was used to quantify total protein levels.

### Immunoassays

#### PGRN

CSF, plasma, and brain progranulin levels were measured using an R&D Systems DuoSet ELISA kit (# D42420). CSF was diluted 4x, plasma was diluted 100X, into sample diluent NS. Brain lysate was diluted to roughly 0.4mg/mL before being diluted 10x into sample diluent NS. Raw absorbance values were interpolated against a calibration curve provided with the assay kit. Brain PGRN data was normalized to protein concentration as measured by BCA.

### NfL

CSF & plasma neurofilament light chain (NfL) levels were analyzed using the Quanterix Simoa Neurofilament Light Advantage (NfL) kit (Quanterix Simoa; Lexington, Massachusetts, USA, # 103400), and analyzed on the SR-X instrument. CSF was diluted 100x, and plasma was diluted 4X, into sample diluent. Raw instrument values were interpolated against a calibration curve provided with the assay kit.

### Tau, GFAP, UCHL1

CSF & plasma Tau, GFAP & UCHL1 levels were analyzed using the Quanterix Simoa Neurology 4-Plex A kit (# 102153) and analyzed on the HD-X instrument. CSF was diluted 40x, and plasma was diluted 4X, into sample diluent. Raw instrument signals were interpolated against calibration curves for each analyte provided with the assay kit.

### YKL40

YKL40 levels in CSF, plasma, and brain were measured using MSD’s U-Plex kit (# K151VLK). CSF was diluted 400x, plasma was diluted 100X, and brain was diluted 2500x into sample diluent. Raw instrument values were interpolated against a calibration curve provided with the assay kit. Brain YKL40 data was normalized to protein concentration as measured by BCA.

## Targeted liquid chromatography with tandem mass spectrometry (LC-MS/MS) analysis for lysosomal biomarkers

### GlcSph in plasma

To prepare samples for LC-MS/MS analysis, aliquots of 20 µL of calibration standards (STDs), quality control samples (QCs), and unknown plasma samples are transferred to a clean 96-well plate. 20 µL of 1% Bovine Serum Albumin (BSA) (Sigma Aldrich cat # A2153-100G) in water solution is added to STDs and QCs while 20 µL of methanol is added to plasma samples and plasma matrix QCs. 10 µL of internal standard working solution (5 ng/mL of GlcSph-d5 (Avanti Polar Lipids cat # 860636P-1mg) in acetonitrile/isopropyl alcohol/water 92.5/5/2.5 with 0.5% Formic Acid and 5 mM ammonium formate) is added to each sample followed by 150 µL of 1% ammonium hydroxide in water to adjust pH. After mixing, the samples are loaded onto an ISOLUTE^®^ SLE+ 200 µL Supported Liquid Extraction Plate (Biotage cat # 820-0200-P01). After the samples are completely absorbed in the plate sorbent, ethyl acetate is used to elute the analyte into a clean collection plate. The eluent is dried down completely under purified nitrogen gas flow at 40 °C and reconstituted in 150 µL of acetonitrile/isopropyl alcohol/water 92.5/5/2.5 with 0.5% Formic Acid and 5 mM ammonium formate before the samples are injected to LC-MS/MS for analysis.

LC-MS/MS analyses are performed on an ExionLC AD UHPLC system coupled with a Sciex API 6500 Triple Quad mass spectrometer (AB Sciex, Redwood City, CA). The MRM transitions for GlcSph (Avanti Polar Lipids cat # 860535P-5mg) and GlcSph-d5 are 462.3 to 282.1 and 467.6 to 287.4, respectively. The Declustering Potential is 60 v and Collisional Energy is 31 v. HPLC chromatography is established on an Acquity BEH HILIC column (1.7 µm, 100 × 2.1 mm) (Waters Co. cat # 186003461) and the column is kept at 55 °C during the run.

For the LC separation of GlcSph from the matrix interference, the two mobile phases used are 0.1% Formic Acid and 10 mM ammonium formate in water (mobile phase A) and 0.1% Formic Acid in acetonitrile (mobile phase B). The flow rate is 0.4 mL/min. Mobile phase B concentration is initially set at 93% and kept for 6.5 min, and then decreased to 50% at 6.51 min and kept for 0.99 min before increased to 93% at 7.51 min. The gradient is ended at 10 min after holding at 93% B for 2.49 min.

### BMP (18:1/18:1) and (22:6/22:6) in plasma and CSF

To prepare plasma samples for LC-MS/MS analysis, aliquots of 50 µL of calibration standards (STDs), quality control samples (QCs), and unknown plasma samples are transferred to a clean 96-well plate. 50 µL of 1% Bovine Serum Albumin (BSA) in water solution is added to STDs and QCs while 50 µL of methanol/isopropyl alcohol 50/50 is added to plasma samples and plasma matrix QCs. 10 µL of internal standards working solution (500 ng/mL of BMP-d5 (22:6/22:6) (Avanti Polar Lipids cat # 792469) and 50 ng/mL of BMP-d5 (18:1/18:1) (Avanti Polar Lipids cat # 792706) in methanol) is added to each sample followed by 100 µL of water to adjust pH. After mixing, the samples are loaded onto an ISOLUTE^®^ SLE+ 200 µL Supported Liquid Extraction Plate. After the samples are completely absorbed in the plate sorbent, ethyl acetate is used to elute the analyte into a clean collection plate. The eluent is dried down completely under purified nitrogen gas flow at 40 °C and reconstituted in 100 µL of methanol/water 70/30 before the samples are injected to LC-MS/MS for analysis.

To prepare CSF samples for LC-MS/MS analysis, aliquots of 50 µL of calibration standards (STDs), quality control samples (QCs), and unknown CSF samples are transferred to a clean 96-well plate. 50 µL of aCSF (Tocris Bioscience cat # A115-50) is added to STDs and QCs while 50 µL of methanol/isopropyl alcohol 50/50 is added to CSF samples and CSF matrix QCs. 10 µL of internal standards working solution (100 ng/mL of BMP-d5 (22:6/22:6) and 10 ng/mL of BMP-d5 (18:1/18:1) in methanol) is added to each sample followed by 300 µL of water to adjust pH. After mixing, the samples are loaded onto an ISOLUTE^®^ SLE+ 400 µL Supported Liquid Extraction Plate (Biotage cat # 820-0400-P01). After the samples are completely absorbed in the plate sorbent, ethyl acetate is used to elute the analyte into a clean collection plate. The eluent is dried down completely under purified nitrogen gas flow at 40 °C and reconstituted in 100 µL of methanol/water 70/30 before the samples are injected to LC-MS/MS for analysis.

LC-MS/MS analyses are performed on an ExionLC AD UHPLC system coupled with a Sciex API 6500+ Triple Quad mass spectrometer. The MRM transitions for BMP (22:6/22:6) (Avanti Polar Lipids cat # N/A, custom order), BMP (18:1/18:1) (Avanti Polar Lipids cat # 857133C-5mg), BMP-d5 (22:6/22:6) and BMP-d5 (18:1/18:1) are 865.6 to 327.0, 773.6 to 281.0, 870.6 to 327.0 and 778.6 to 281.0, respectively. The Declustering Potential is -60 v and Collisional Energy is -45 v and -51 v for BMP (22:6/22:6) and BMP (18:1/18:1), respectively. HPLC chromatography is established on an Acquity BEH C8 column (1.7 µm, 100 × 2.1 mm) (Waters Co. cat # 91813-701) and the column is kept at 55 °C during the run.

For the LC separation of BMP species from the matrix interference, the two mobile phases used are 0.01% ammonium hydroxide and 1 mM ammonium acetate in water (mobile phase A) and 0.01% ammonium hydroxide in acetonitrile (mobile phase B). The flow rate is 0.5 mL/min. Mobile phase B concentration is initially set at 75% and kept for 0.2 min before increasing to 90% for 3.3 min, and then to 95% at 3.51 min. Mobile phase B concentration was kept at 95% until 4.50 min before decreasing to 75% at 4.51 min. The gradient is ended at 10 min after holding at 75% B for 5.49 min.

### GM2 (d36:1) in CSF

To prepare CSF samples for LC-MS/MS analysis, aliquots of 25 µL of calibration standards (STDs), quality control samples (QCs), and unknown CSF samples are transferred to a clean 96-well plate. 25 µL of aCSF is added to STDs and QCs while 25 µL of methanol/isopropyl alcohol 50/50 is added to CSF samples and CSF matrix QCs. 100 µL of internal standard working solution (0.5 ng/mL of GM2-d7 (Matreya LLC Lipids and Biochemicals cat # 9003931) is added to each sample. The samples are mixed and then centrifuged (10 min at 4 °C, 3000 rpm) before injected to LC-MS/MS for analysis.

LC-MS/MS analyses are performed on an ExionLC AD UHPLC system coupled with a Sciex API 7500 Triple Quad mass spectrometer (AB Sciex, Redwood City, CA). The MRM transitions for GM2 (Matreya LLC Lipids and Biochemicals cat # 1542) and GM2-d7 are 1382.7 to 290.1 and 1389.7 to 290.1, respectively. The Entrance Potential is -10 v and Collisional Energy is -76 v. HPLC chromatography is established on a Thermo Scientific Hypersil GOLD C4 (1.9 µm, 50 × 2.1 mm) (Thermo Fisher Scientific cat # 25502-052130) and the column is kept at 55 °C during the run.

For the LC separation of GM2 species from the matrix interference, the two mobile phases used are 0.01% acidic acid and 5 mM ammonium acetate in water (mobile phase A) and 0.01% acidic acid in acetonitrile (mobile phase B). The flow rate is 0.6 mL/min. Mobile phase B concentration is initially set at 60% and increased to 68% B for 2 min, and then to 95% at 2.01 min. The gradient was kept at 95% B for 0.99 min before decreasing to 5% B at 3.01 min and kept at 5% B for 0.99 min. The gradient is increased to 60% B at 4.01 min and is ended at 5 min after holding at 60% B for 0.99 min.

### GlcSph in CSF

CSF GlcSph was quantified at a CRO (KCAS Bioanalytical & Biomarker Services, Olathe, KS). To prepare CSF samples for LC-MS/MS analysis, aliquots of 100 µL of calibration standards (STDs), quality control samples (QCs), and unknown CSF samples are transferred to a clean 96-well plate. 300 µL of MPAB (Mixture of mobile phase A and mobile phase B: 97.5:2.5 (v/v))) with ISTD (2 pg/mL of Glucosyl(ß) Sphingosine-d5 (Avanti Polar Lipids, Catalog# 860636)) is added to STDs, QCs, CSF samples and CSF matrix QCs. The samples are vortexed and then centrifuged (5 min at 4°C, 3500 rpm) before transferring 350uL to a fresh 96-well plate. The plate is dried at 37°C until completely dry before reconstituted in 100uL of MPAB and injected to LC-MS/MS for analysis.

LC-MS/MS analyses are performed on a Shimadzu LC AD HPLC system coupled with a Triple Qtrap 7500+ LC-MS/MS with Turbo-Ion Spray Interface mass spectrometer (Sciex). The MRM transitions for GlcSph (Vendor & Cat #) and GlcSph-d5 are 462.4 to 282.3 and 467.3 to 287.3, respectively. The Entrance Potential is -10 v and Collisional Energy is -30 v. HPLC chromatography is established on a Halo Hilic (2.7 µm, 4.6 × 150 mm) (Advanced Materials Technology Cat #92814-701) and the column is kept at 50°C during the run.

For the LC separation of GlcSph species from the matrix interference, the two mobile phases used are 0.5% Formic Acid in ACN (mobile phase A) and 50 mM Ammonium Formate in 60/40/0.1 MeOH/DiH2O/Formic Acid (mobile phase B). The flow rate is 1.00 mL/min. Mobile phase B concentration is initially set at 25% and increased to 45% B over 3.10 min, and then to 80% at 3.20 min. The gradient was kept at 80% B for 0.30 min before decreasing to 25% B at 3.60 min and kept at 25% B for 1.90 min.

## Targeted liquid chromatography-mass spectrometry (LCMS) analysis for metabolites and lipids

### CSF and plasma extraction

Plasma samples were diluted 1:20, and CSF diluted 1:5, into methanol + internal standards. Samples were then shaken (5 min at 4°C, 700rpm), incubated for 1 hr at -20°C, then centrifuged (20 min at 4°C, 4,000 x g), followed by transfer of supernatant to a 96 well plate with glass inserts and sealed in preparation for LCMS analysis.

### CSF

CSF Metabolite and lipid analyses were performed by liquid chromatography (UHPLC Nexera X2) coupled to electrospray mass spectrometry (QTRAP 6500+).

Lipids were analyzed in both positive and negative mode, using an UPLC BEH C18 column (150 × 2.1 mm, 1.7 μm particle size, Waters Corp.), maintained at 55 °C, with gradient elution, and flow rate of 0.25 mL/min over 10 min gradient as described in (Logan et al., 2021^20^).

Metabolites were analyzed in positive mode, using a UPLC BEH amide column (150 × 2.1 mm, 1.7 μm particle size, Waters Corp.), maintained at 40 °C, with gradient elution, and flow rate of 0.40 mL/min over 10 min gradient as described in (Logan et al., 2021^20^).

Metabolites were analyzed in negative mode, using an InfinityLab Poroshell 120 HILIC-Z P (50 × 2.1 mm, 2.7 μm particle size, Agilent Technologies Inc.), maintained at 20°C, with gradient elution, and flow rate of 0.50 mL/min over 10 min gradient in negative ionization mode. For negative ionization mode, Mobile phase A consisted of water with 10 mM ammonium acetate, pH 9. Mobile phase B consisted of acetonitrile:water 9:1 with 10 mM ammonium acetate, pH 9. The gradient was programmed as follows: 0.0–1.0 min at 95% B; 1.0–6.0 min to 50% B; 6.0–6.5 min to 95% B; and 6.5–10.0 min at 95% B. Electrospray ionization was performed in negative ion mode. For the QTRAP 6500+ we applied the following settings: curtain gas at 40 V; collision gas was set at medium ion spray voltage at -4500 V; temperature at 600°C; ion source Gas 1 at 50 psi; ion source Gas 2 at 60 psi; entrance potential at -10 V; and collision cell exit potential at -15.0 V.

See Supplemental Tables 3 & 4 for Metabolomic and Lipidomic run parameters.

### Plasma

Plasma metabolite and lipid analyses were performed as described for CSF with minor modifications to LCMS system used for lipidomic analysis in the positive mode. Lipidomic analysis in positive mode was performed on liquid chromatography (Agilent Infinity II1290) coupled to electrospray mass spectrometry (TQ 6495C). For the Agilent TQ 6495C we applied the following settings: gas temp at 180°C; gas flow 17 l/min; nebulizer 35 psi; sheath gas temp 350°C; sheath gas flow 10l/min; capillary 3500V; nozzle voltage 500 V.

### Proteomic analysis of CSF and brain tissues

A targeted proteomic assay for a selected panel of protein biomarkers including neuronal pentraxin receptor (NPTXR), neuronal pentraxin 1 (NPTX1), neuronal pentraxin 2 (NPTX2), and 14–3-3 proteins (eta, epsilon, zeta/delta) was developed. Supplemental Table 1 shows the proteins and their respective proteotypic peptides targeted in the multiple reaction monitoring mass spectrometry analysis (MRM MS). To monitor the performance of the assay over time, quality control (QC) sample replicates were injected at regular intervals during runs.

### CSF in solution digest

In solution digest of CSF was performed on an AssayMap Bravo liquid handling instrument (Agilent) using a custom designed protocol. 100uL CSF, to which 5 µL Pierce digest indicator (Thermo 84841) is added to a final concentration of 0.15 μg. DTT is added to a concentration of 6.4 mM and heated at 60°C for 30 min. The samples are cooled for an additional 30 min followed by the addition of 12 mM iodoacetamide at room temperature for 30 min. 2 µg Mass Spec grade Trypsin/LysC (Promega) is added to each sample and incubated at 37°C overnight. The digestion is halted by acidifying the samples to 12.5% formic acid.

### Brain tissue digest

Lysate was precipitated in acetone at -20°C for 4 hr, centrifuged at 15,000 xg for 20 min and the pellet resuspended in 8 M urea in 100 mM Tris, pH of 8. The sample is reduced with 5 mM DTT at 37°C for 1 hr then alkylated with 10 mM iodoacetamide for 30 min in the dark. Next the sample is diluted 5:1 with 100 mM triethylammonium bicarbonate buffer and digested with Trypsin/LysC at 37°C overnight. The digestion is quenched with 1% formic acid and sample desalted using the RPS cartridges (Agilent)

### Multiple reaction monitoring mass spectrometry (MRM MS)

Following the digestion, 6 µL of stable isotopically labelled (SIL) synthetic peptides (Cell Signaling Technologies) master mix was added as an internal standard for every target peptide at a concentration of 300 fmol. The full list of target peptides and transitions can be found in Supplemental Table 5. The samples were then desalted using 25 µL RPS AssayMap cartridges (G5496-60023). Briefly, the cartridges were primed with 250 µL 50% ACN/0.1% TFA, conditioned with 200 µL 0.1% TFA, loaded with 200 µL of sample, washed with 250 µL 0.1% TFA, then finally eluted in 75 µL 50% ACN/0.1% formic acid. The eluted peptide was dried in a Speedvac (Genevac) then reconstituted in 40 µL 0.1% formic acid. The analysis was performed using an Agilent 1290 Infinity II UPLC system (1290 high speed binary pump, a thermostat autosampler and a thermostat multicolumn compartment) coupled with the Agilent 6495c triple-quadruple mass spectrometer. Approximately 25 ug of purified tryptic digest were injected on an Agilent Eclipse Plus C18 trap column (2.1×5mm, 8µm) connected to an Agilent Eclipse Plus C18 column (2.1×150mm, 8µm). Peptides were loaded for 1 minute with an initial mobile phase composition of 97% Aqueous Formic Acid (Mobile Phase A) and 3% Acetonitrile, 0.1% Formic Acid (Mobile Phase B). The peptides were separated using a linear gradient from 3-38%B in 51 minutes at a flow rate of 0.2 ml/min and with constant column temperature of 50°C. The Agilent 6495c triple-quadruple mass spectrometer was operated in positive ion mode with MRM scan type. The Agilent AJS ESI source was set up as follow: Gas temperature 220°C, Gas Flow 19 L/min, Nebulizer 25psi, Sheath Gas Temperature 275°C, Sheath Gas Flow 11 L/min, Capillary Voltage 3250V. The additional Electron Multiplier Voltage (Delta EMV) was set at +400V and the two ion funnels RF were set at 150V and 60V for the high and low pressure. Data was analyzed in Skyline, peptide quantity is expressed as the ratio of the endogenous (Light) peptide to the standard (SIL) peptide.

### Statistical analyses

All analyses were performed using R statistical software. Measured values were transformed using a log base 2 transformation. In the biomarker analysis of plasma and CSF samples, differences between groups were estimated using a linear mixed-effects model, with random subject-level intercepts to address the repeated measurements. In the linear mixed-effects models, p-values for fixed effects were evaluated using Wald tests, which compare the ratio of the estimated coefficient to its standard error. To account for the lack of balance in age and sex across groups, both covariates were adjusted across all analytes. In situations where age or sex information was missing for certain subjects in the selected population, no adjustments for demographics information were made in the model. For the analysis of brain samples, differences between groups and p-values were estimated using a robust linear model to minimize the impact of outliers on regression analysis, with age at death and sex included as covariates in the model [^31^]. Robust linear model, a form of weighted least squares regression, uses robust standard errors and t-statistics to obtain p-values.

Spearman’s rank correlation, a non-parametric statistical method, was used to estimate the strength and direction of association without assuming a linear relationship between biomarkers and CDR+NACC FTLD-SB scores. This analysis includes only the initial available sample in the dataset that corresponds to a CDR+NACC FTLD-SB score greater than zero for each subject.

To examine the association between baseline NfL level and annualized change in CDR+NACC FTLD-SB score in S-*GRN* subjects, spearman’s rank correlation was used. The annualized change in CDR+NACC FTLD-SB score was incorporated through a linear regression model for each subject. In this analysis, baseline refers to the initial visit in the dataset where the CDR+NACC FTLD-SB is greater than zero. Only *GRN* carriers, possessing at least one visit post-baseline in the dataset with both NfL and an available CDR+NACC FTLD-SB score, were included.

For longitudinal analysis, we only focused on groups with more than 5 subjects with longitudinal samples, and thus only examined *GRN* carriers. The longitudinal trends in plasma and CSF samples were assessed using the mean annual change along with its corresponding 95% confidence interval. Subjects were classified into 3 groups: asymptomatic *GRN* carriers (AS-*GRN*), symptomatic (S-*GRN*) and converters. AS-*GRN* were defined as *GRN* carriers with CDR+NACC FTLD-SB scores consistently equal to 0. S-*GRN* were defined as *GRN* carriers with CDR+NACC FTLD-SB scores consistently exceeding 0. Converters were *GRN* subjects with CDR+NACC FTLD-SB scores that changed during their longitudinal visits (either from 0 to > 0, or > 0 back to 0).

## Ethical approval

This study involves human participants and was approved by Johns Hopkins Medicine IRB serves as the Single IRB for the ALLFTD Consortium (CR00042454 / IRB00227492). All participating sites additionally obtain local IRB approvals. Participants gave informed consent to participate in the study before taking part.

## Data availability

All data needed to evaluate the conclusions in the paper are present in the paper and/or the Supplemental Materials. ALLFTD data are available upon request from qualified investigators (www.allftd.org/data). Additional data request can be made by contacting the corresponding authors.

## Results

For CSF and plasma samples, the demographic and clinical features of study groups are listed in Table 1. The median ages in the asymptomatic groups were younger than the symptomatic groups, except the *MAPT* group. A majority of the subjects were white. Mixed clinical phenotypes were present in symptomatic FTD groups. For the sporadic FTD group, 53% presented with corticobasal syndrome, with only 17% behavioral variant FTD (bvFTD) and 28% primary progressive aphasia (PPA). The symptomatic mutation carriers had a significant percentage of bvFTD (41% in S-*GRN*, 64% in S-*C9orf72*, 68% in S-*MAPT*), with a lower percentage of PPA (19% in S-*GRN*, 5% in S-*C9orf72,* 0% in S-*MAPT*).

For the biomarker study with postmortem brain tissues, three brain regions were obtained: two disease-impacted degenerating regions: middle temporal gyrus (MTG) and superior frontal gyrus (SFG), and a generally spared region inferior occipital gyrus (IOG). FTLD-TDP type A were detected in sporadic FTD (S-NC) and symptomatic *GRN* mutation carriers (S-*GRN*). The ages were similar across the groups and there was a higher percentage of males in the S-*GRN* group as compared to the non-carrier control group (Table 2).

### PGRN levels reduced in *GRN* mutation carriers

To characterize the samples, we first determined the PGRN levels in CSF, plasma, and brain tissues. As expected, PGRN was reduced in the biofluid of symptomatic *GRN* carriers (S-*GRN*: CSF 49.7%, p<0.0001; plasma 30.0%, P<0.0001) and asymptomatic *GRN* carriers (AS-*GRN*: CSF 48.6%, p<0.0001; plasma 28.6%, p<0.0001) compared to non-carrier controls (Supplemental Fig. 1A and 1B). In brain tissues, PGRN is most reduced in IOG from symptomatic *GRN* carriers compared to the same region from controls (IOG, 55.9%, P<0.001; MTG 90.4%, p = 0.315; SFG 78.9%, P=0.047) (Supplemental Fig. 1C).

**Figure 1:**
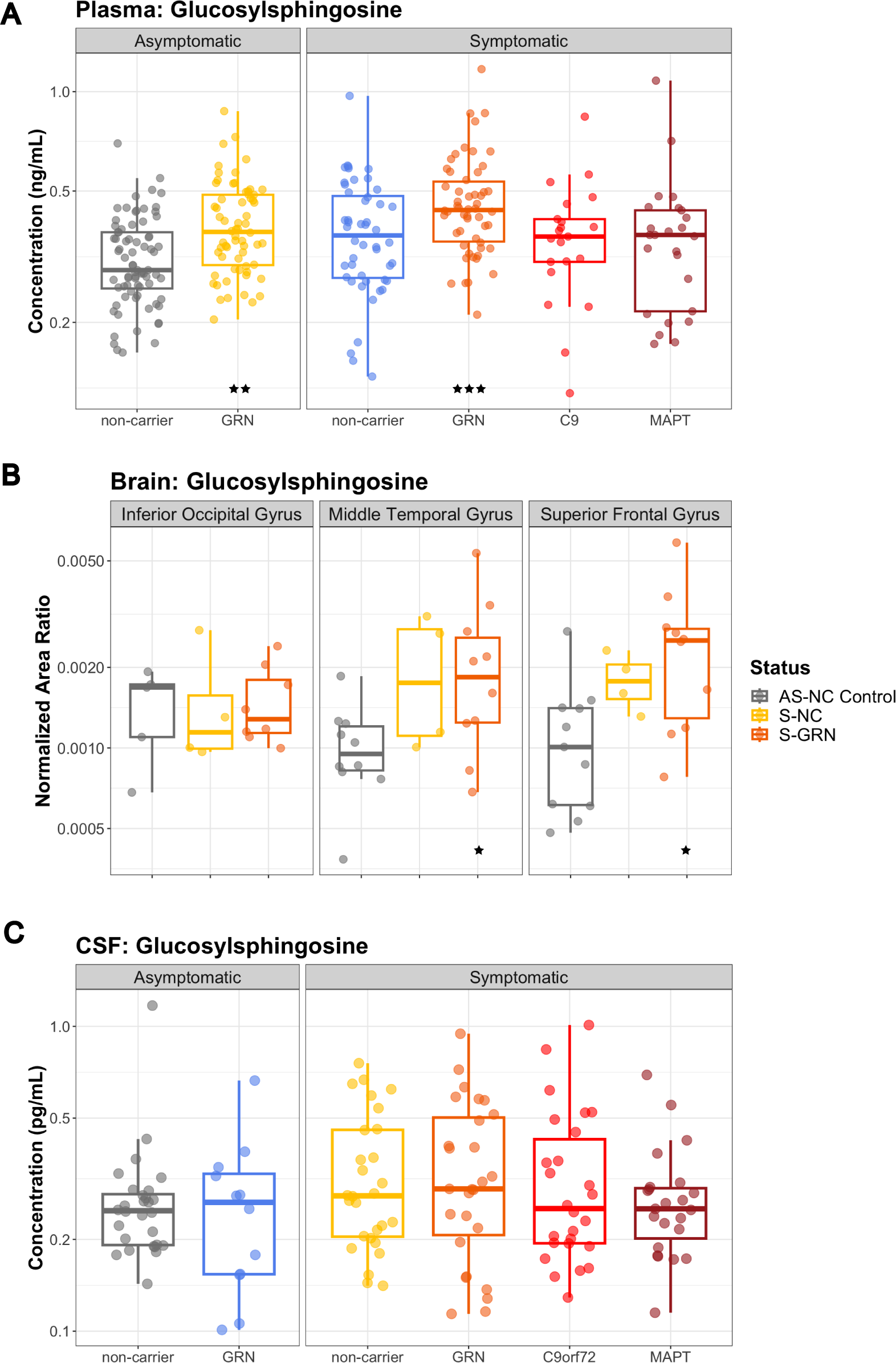
Lysosomal biomarker GlcSph in plasma, CSF and brain tissues. (A) Plasma GlcSph levels in AS-NC controls, AS-*GRN*, S-NC, S-*GRN*, S-*C9orf72* and S-*MAPT*, (B) GlcSph in IOG, MTG and SFG from AS-NC Control, S-NC and S-*GRN*, (C) CSF GlcSph in AS-NC controls, AS-*GRN*, S-NC, S-*GRN*, S-*C9orf72* and S-*MAPT.* For CSF/plasma, each dot represents an individual subject visit. For brain tissues, each dot represents an individual subject. Box plots are median ± interquartile range (IQR). Statistical analysis was performed in comparison to AS-NC controls, *p<0.05, **p<0.01, ***p<0.001. AS-NC: asymptomatic non-carrier controls; AS-*GRN*: asymptomatic *GRN* mutation carriers; S-NC: symptomatic non-carriers or sporadic FTD; S-*GRN*: symptomatic *GRN* mutation carriers; S-*C9orf72*: symptomatic *C9orf72* mutation carriers; S-*MAPT*: symptomatic *MAPT* mutation carriers; IOG: inferior occipital gyrus; MTG: medial temporal gyrus; SFG: superior frontal gyrus.

### Lysosomal biomarkers altered in patients with FTD

Consistent with the previously published data [^20^], GlcSph was significantly elevated in plasma from GRN mutation carriers (S-*GRN*, 136.6%, P<0.001; AS-*GRN*,127.6%, P=0.002) compared to non-carrier controls (Fig. 1A). There were no significant changes of plasma GlcSph in other symptomatic groups (Fig. 1A). GlcSph concentrations in CSF were much lower than plasma, and a trend of elevation in CSF for S-*GRN* was observed (120.9%, P=0.182) (Fig. 1C). In brain tissues, GlcSph was significantly increased in temporal and frontal gyrus but not in occipital gyrus from S-*GRN* compared to controls (IOG: 101.9% P=0.942; MTG: 176.4% P=0.040; SFG: 180.6%, P=0.042) (Fig. 1B). Trends of increased GlcSph in disease-affected brain regions were also observed in sporadic FTD (S-NC, IOG: 84.1% P=0.574; MTG: 179.2% P=0.076; SFG: 173.8%, P=0.097) (Fig. 1B).

In addition, lysosomal ganglioside GM2(d36:1) was significantly increased in the disease-affected brain regions of S-*GRN* (IOG: 138.5% P=0.064; MTG: 192.7% P<0.001; SFG: 168.9%, P=0.001) and in all three brain regions of sporadic FTD (S-NC, IOG: 180.5%, P=0.008; MTG: 146.0%, P=0.003; SFG: 161.2%, P=0.012) (Fig. 2A). No significant changes were detected for GM2(d36:1) in the CSF of patients with FTD, with or without genetic mutations (Fig. 2B). The lysosomal globoside GB3(d18:1/18:0), also known as globotriaosylceramide, was highly increased in the disease-affected brain regions from S-*GRN* (IOG: 208.2% P=0.169; MTG: 1295.0% P<0.001; SFG: 1355.4%, P<0.001) and in sporadic FTD compared to non-carrier controls (IOG: 113.7% P=0.843; MTG: 255.9% P=0.070; SFG: 355.3%, P=0.033) (Fig. 2C). GB3 in CSF was below the detection limit of the current assay.

**Figure 2:**
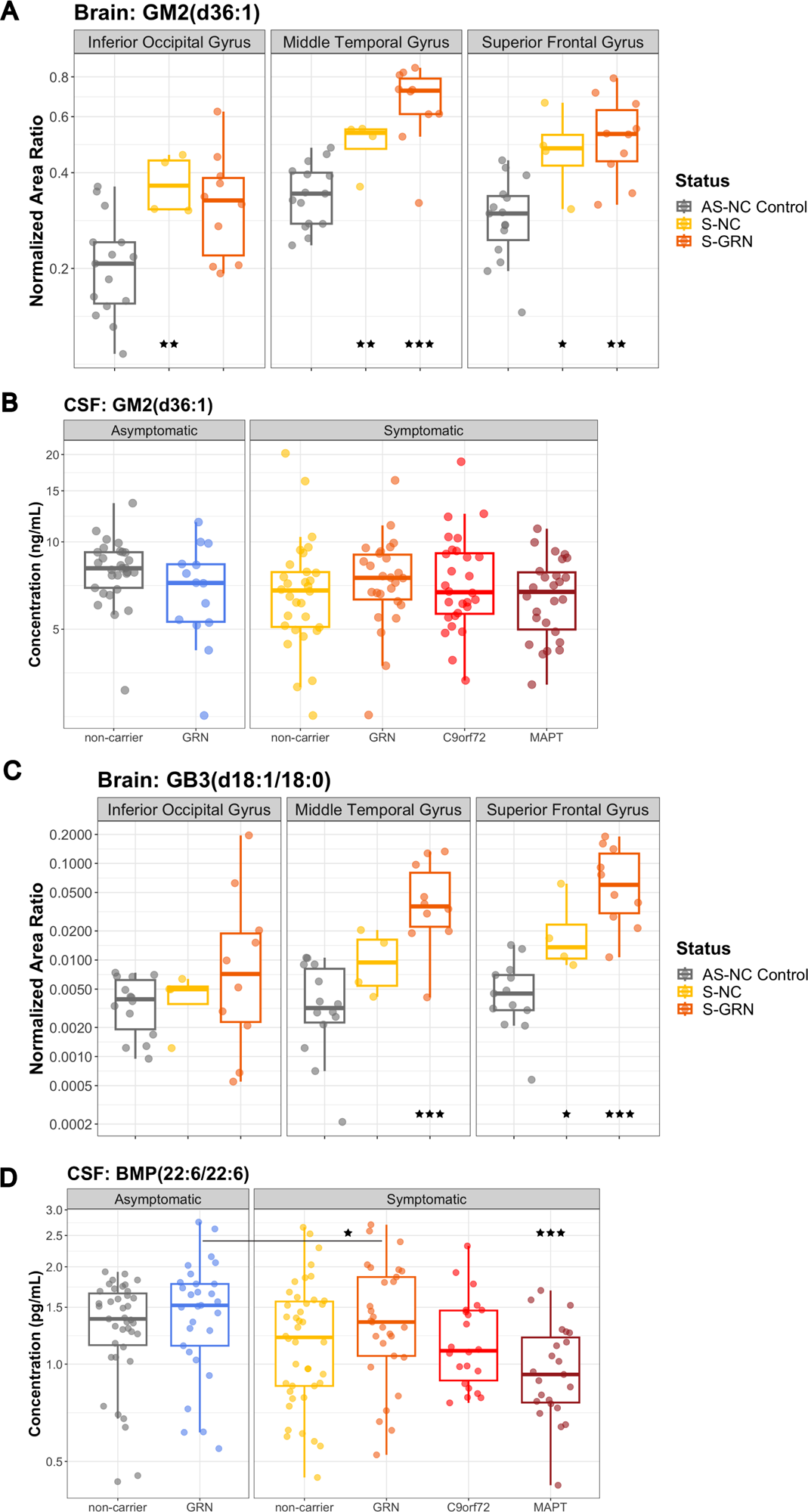
Lysosomal biomarkers GM2 and GB3 in biofluids and brain tissues. (A) GM2(d36:1) and (C) GB3(d18:1/18:0) in IOG, MTG and SFG from AS-NC Control, S-NC and S-*GRN*. (B) GM2(d36:1) and (D) BMP(22:6/22:6) in CSF from AS-NC controls, AS-*GRN*, S-NC, S-*GRN*, S-*C9orf72* and S-*MAPT*. For CSF, each dot represents an individual subject visit. For brain tissues, each dot represents an individual subject. Box plots are median ± interquartile range (IQR). Statistical analysis was performed in comparison to AS-NC controls, and an additional comparison made between S-*GRN* and AS-*GRN* in (D). *p<0.05, **p<0.01, ***p<0.001. AS-NC: asymptomatic non-carrier controls; AS-*GRN*: asymptomatic *GRN* mutation carriers; S-NC: symptomatic non-carriers or sporadic FTD; S-*GRN*: symptomatic *GRN* mutation carriers; S-*C9orf72*: symptomatic *C9orf72* mutation carriers; S-*MAPT*: symptomatic *MAPT* mutation carriers; IOG: inferior occipital gyrus; MTG: medial temporal gyrus; SFG: superior frontal gyrus.

For another key lysosomal biomarker bis(monoacylglycero)phosphate (BMP), a significant reduction of CSF BMP(22:6/22:6) was observed in symptomatic *MAPT* group compared to non-carrier controls (S-NC: 80.9%, P=0.064; S-*GRN*: 82.0%, P=0.106; S-*C9orf72*: 83.0%, P=0.131; S-*MAPT*: 71%, P=0.005) (Fig. 2D). Notably, a significant reduction of CSF BMP(22:6/22:6) was also observed for symptomatic *GRN* carriers as compared to asymptomatic *GRN* carriers (83.3%, P=0.031) (Fig. 2D). Intriguingly, BMP(22:6/22:6) was significantly increased in brain tissues of IOG and MTG in S-*GRN* (IOG: 203.7% P=0.002; MTG: 173.5% P=0.005; SFG: 154.6%, P=0.053) and sporadic FTD (S-NC, IOG: 238.5% P=0.002; MTG: 171.4% P=0.028; SFG: 120.4%, P=0.486) compared to non-carrier controls (Supplemental Fig. 2A). Similar trend of changes were also observed for BMP(18:1/18:1) in CSF and brain tissues as for BMP(22:6/22:6) (data not shown). BMP(22:6/22:6) in plasma was generally unchanged, except for a significant elevation in sporadic FTD (S-NC: 164.5%, P=0.009; S-*GRN*: 109.5%, P=0.621; S-*C9orf72*: 118.6%, P=0.452; S-*MAPT*: 80.6%, P=0.300; Supplemental Fig. 2B).

### Glial biomarkers altered in patients with FTD

Consistent with the occurrence of astrogliosis, levels of the reactive astrocyte marker, glial fibrillary acidic protein (GFAP), were increased in plasma from symptomatic groups, except *C9orf72,* relative to non-carrier controls (S-NC: 131.9%, P=0.021; S-*GRN*: 146.7%, P=0.001; S-*C9orf72*: 97.3%, P=0.845; S-*MAPT*: 129.5%, P=0.050) (Fig. 3A). GFAP in plasma was also increased in the AS-*GRN* (127.6%, P=0.030) (Fig. 3A). No significant increases of GFAP were detected in the CSF samples (Fig. 3B).

**Figure 3:**
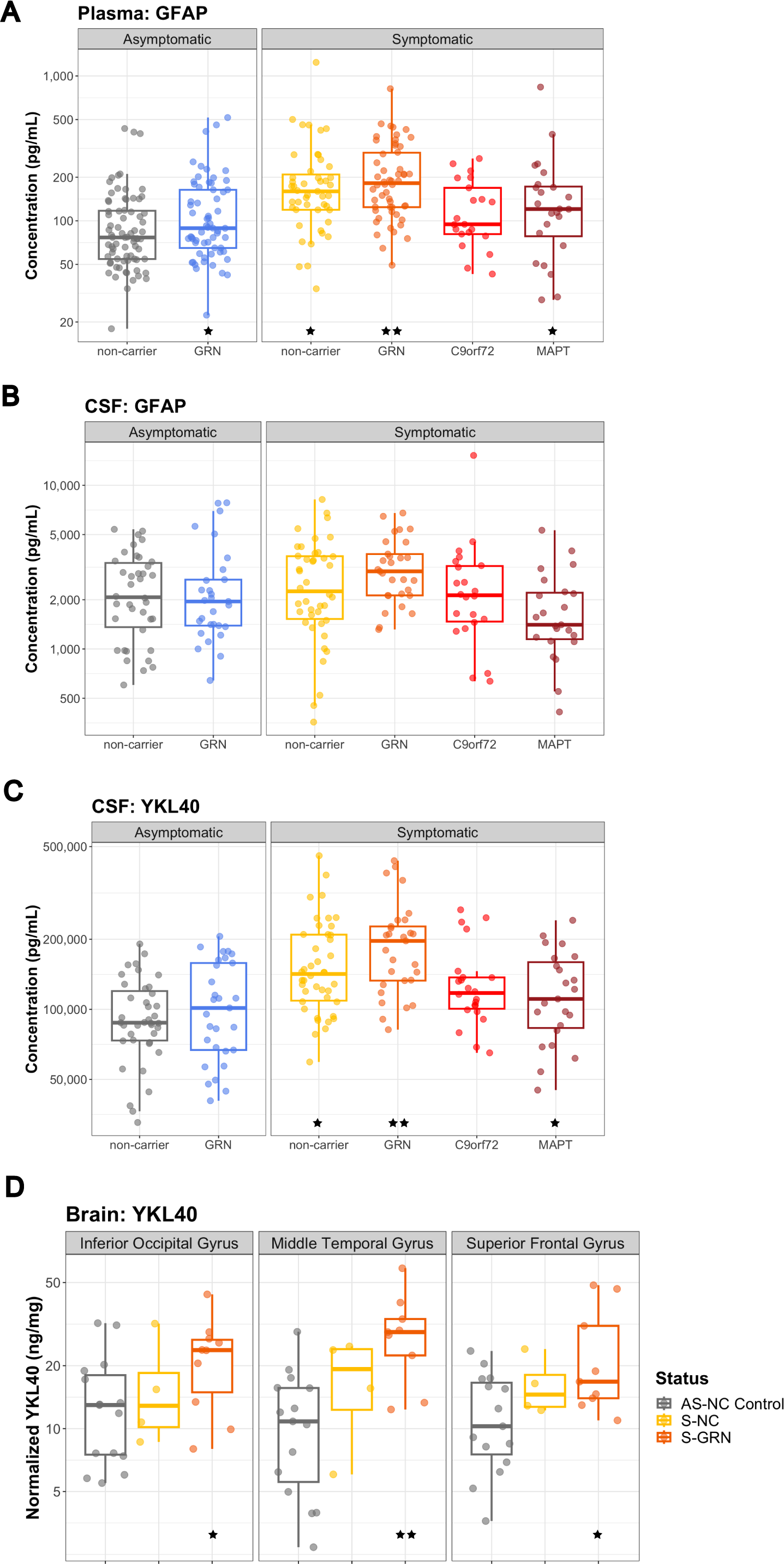
Glial biomarkers GFAP and YKL40 in biofluids and brain tissues. (A) CSF GFAP, (B) plasma GFAP, and (C) CSF YKL40 from AS-NC controls, AS-*GRN*, S-NC, S-*GRN*, S-*C9orf72* and S-*MAPT*. (D) YKL40 in IOG, MTG and SFG from AS-NC Control, S-NC and S-*GRN*. For CSF/plasma, each dot represents an individual subject visit. For brain tissues, each dot represents an individual subject. Box plots are median ± interquartile range (IQR). Statistical analysis was performed in comparison to AS-NC controls, *p<0.05, **p<0.01. AS-NC: asymptomatic non-carrier controls; AS-*GRN*: asymptomatic *GRN* mutation carriers; S-NC: symptomatic non-carriers or sporadic FTD; S-*GRN*: symptomatic *GRN* mutation carriers; S-*C9orf72*: symptomatic *C9orf72* mutation carriers; S-*MAPT*: symptomatic *MAPT* mutation carriers; IOG: inferior occipital gyrus; MTG: medial temporal gyrus; SFG: superior frontal gyrus.

Another glial biomarker YKL40 was significantly increased in CSF of symptomatic groups, except *C9orf72,* relative to non-carrier controls (S-NC: 128.4%, P=0.025; S-*GRN*: 139.7%, P=0.006; S-*C9orf72*: 121.3%, P=0.108; S-*MAPT*: 127.0%, P=0.038; Fig. 3C). Increase of YKL40 was also observed in AS-*GRN* (127.6%, P=0.030) (Fig. 3C). In the brain tissues, YKL40 was significantly increased in all three brain regions of symptomatic *GRN* mutation carriers compared to controls (S-*GRN*, IOG: 183.0%, P=0.050; MTG: 293.2%, P=0.002; SFG: 180.2%, P=0.020) (Fig. 3D). Plasma YKL40 levels were not altered in any group (Supplemental Fig. 3A).

### Neurodegeneration biomarkers altered in patients with FTD

Consistent with previous publications [^23^, ^32^, ^33^, ^34^], NfL, a biomarker for axonal injury and neuronal degeneration, was elevated in CSF in all symptomatic groups, as well as asymptomatic *GRN* carriers, as compared to non-carrier controls (AS-*GRN*: 193.2%, P=0.007; S-NC: 202.5%, P=0.003; S-*GRN*: 266.1%, P<0.001; S-*C9orf72*: 275.4%, P<0.001; S-*MAPT*: 234.8%, P<0.001) (Fig. 4A). Plasma NfL was also elevated in all symptomatic groups, as well as AS-*GRN* (AS-*GRN*: 141.0%, P=0.008; S-NC: 197.4%, P<0.001; S-*GRN*: 305.9%, P<0.001; S-*C9orf72*: 236.1%, P<0.001; S-*MAPT*: 215.4%, P<0.001) (Fig. 4B). Brain NfL levels measured by MRM MS proteomics were decreased in all three brain regions of symptomatic groups (peptide sequence VLEAELLVLR, S-NC, IOG: 41.1%, P=0.018; MTG: 35.2%, P=0.008; SFG: 24.4%, P=0.002; S-*GRN*, IOG: 40.3%, P=0.005; MTG: 25.7%, P<0.001; SFG: 11.8%, P<0.001) as compared to controls (Supplemental Fig. 4B).

**Figure 4:**
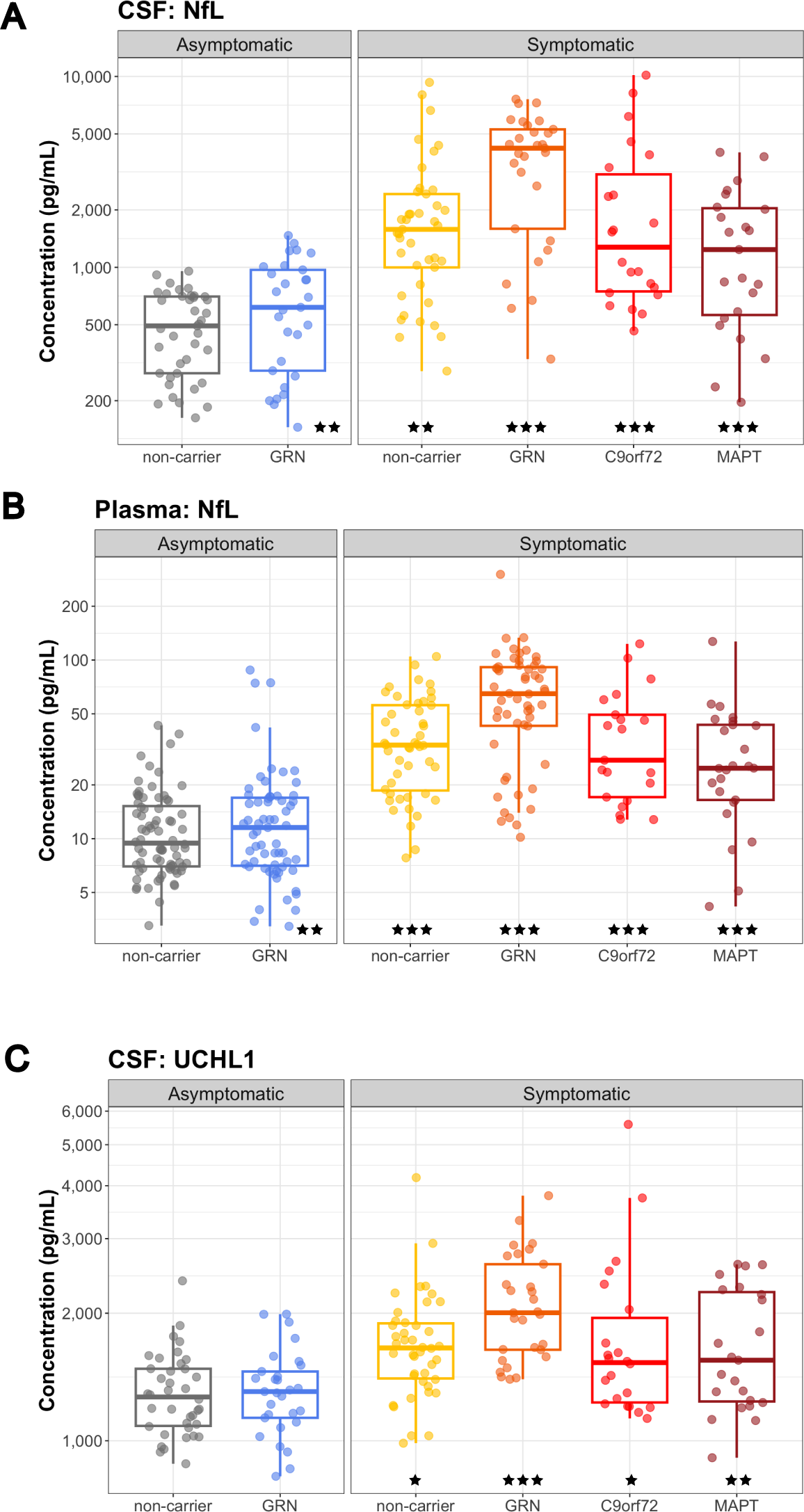
Neurodegeneration biomarkers NfL and UCHL1 in biofluids. (A) CSF NfL, (B) plasma NfL, and (C) CSF UCHL1 from AS-NC controls, AS-*GRN*, S-NC, S-*GRN*, S-*C9orf72* and S-*MAPT*. Each dot represents an individual subject visit. Box plots are median ± interquartile range (IQR). Statistical analysis was performed in comparison to AS-NC controls, *p<0.05, **p<0.01, ***p<0.001. AS-NC: asymptomatic non-carrier controls; AS-*GRN*: asymptomatic *GRN* mutation carriers; S-NC: symptomatic non-carriers or sporadic FTD; S-*GRN*: symptomatic *GRN* mutation carriers; S-*C9orf72*: symptomatic *C9orf72* mutation carriers; S-*MAPT*: symptomatic *MAPT* mutation carriers.

Ubiquitin C-Terminal Hydrolase L1 (UCHL1) was elevated in the CSF of all symptomatic groups as compared to non-carrier controls (S-NC: 117.4%, P=0.044; S-*GRN*: 142.6%, P<0.001; S-*C9orf72*: 123.5%, P=0.015; S-*MAPT*: 126.7%, P=0.005; Fig. 4C). In plasma, a trend towards elevation was observed in symptomatic *GRN* group as compared to non-carrier controls (S-*GRN*: 128.7%, P=0.068) (Supplemental Fig. 4C). UCHL1 measured by MRM MS proteomics was decreased in disease-affected brain regions in symptomatic GRN mutation carriers compared to non-carrier controls (peptide sequence LGFEDGSVLK, S-*GRN*, IOG: 79.5%, P=0.465; MTG: 40.6%, P=0.086; SFG: 40.0%, P=0.004) (Supplemental Fig. 4B).

### Synaptic biomarkers dysregulated in patients with FTD

Synaptic biomarker changes were determined by a targeted proteomic panel of 27 proteins (Supplemental Table 1). Since we were focused on brain relevant biomarkers, we only evaluated this panel in the CSF and brain samples, and not plasma. Neuronal pentraxin 1 (NPTX1), neuronal pentraxin 2 (NPTX2) and neuronal pentraxin receptor (NPTXR) were significantly reduced in CSF in all symptomatic groups with the highest changes observed in S-*C9orf72* & S-*MAPT*. For NPTXR, the decrease in CSF was significant for all symptomatic groups as well as asymptomatic *GRN* carriers compared to non-carrier controls (peptide sequence LVEAFGGATK, AS-*GRN*: 67.7%, P=0.027; S-NC: 62.5%, P=0.005; S-*GRN*: 59.3%, P=0.005; S-*C9orf72*: 52.7%, P<0.001; S-*MAPT*: 41.7%, P<0.001) (Fig. 5A). In brain tissues, NPTXR was significantly decreased in frontal gyrus for S-*GRN* compared to non-carrier controls (peptide sequence LVEAFGGATK, IOG: 91.2%, P=0.712; MTG: 150.5%, P=0.113; SFG: 59.8%, P=0.043) (Supplemental Fig. 5A).

**Figure 5:**
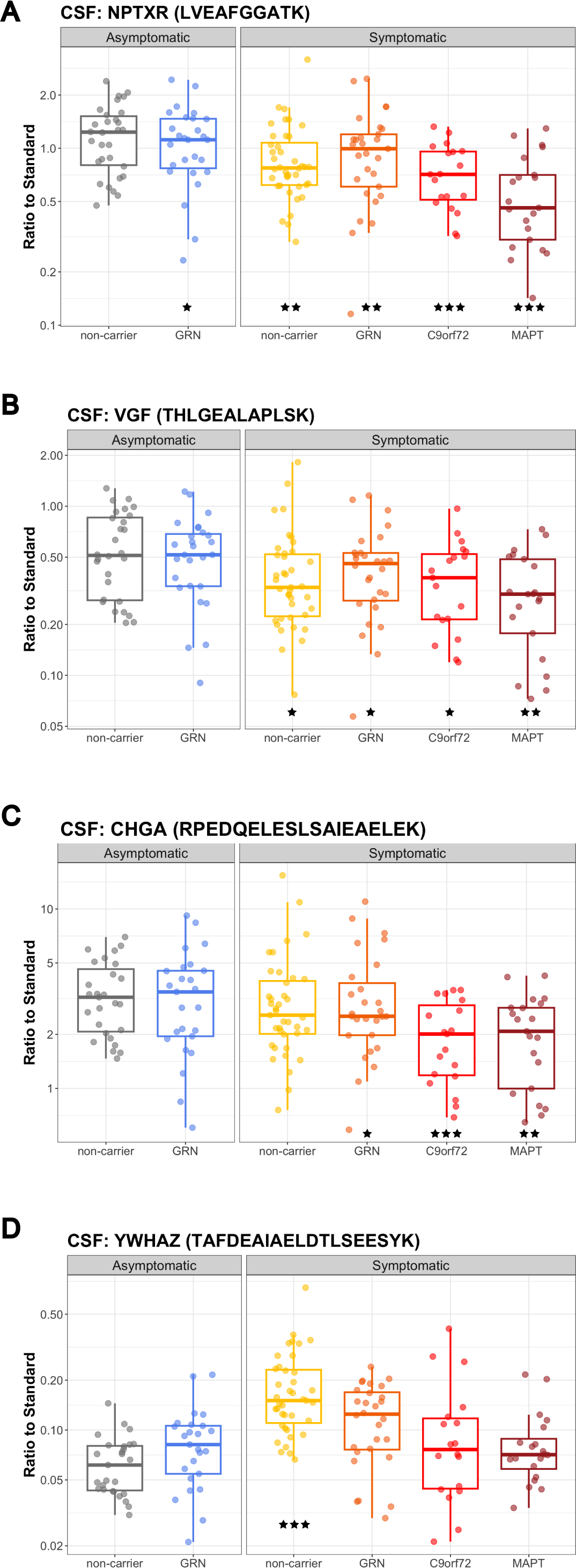
Synaptic biomarkers in CSF. (A) NPTXR (peptide sequence: LVEAFGGATK) (B) VGF (THLGEALAPLSK) (C) CHGA (RPEDQELESLSAIEAELEK) and (D) YWHAZ (TAFDEAIAELDTLSEESYK) in CSF samples from AS-NC controls, AS-GRN, S-NC, S-GRN, S-C9orf72 and S-MAPT. Each dot represents an individual subject visit. Box plots are median ± interquartile range (IQR). Statistical analysis was performed in comparison to AS-NC controls, *p<0.05, **p<0.01, ***p<0.001 (unadjusted for multiple comparison in the proteomic panel). AS-NC: asymptomatic non-carrier controls; AS-*GRN*: asymptomatic *GRN* mutation carriers; S-NC: symptomatic non-carriers or sporadic FTD; S-*GRN*: symptomatic *GRN* mutation carriers; S-*C9orf72*: symptomatic *C9orf72* mutation carriers; S-*MAPT*: symptomatic *MAPT* mutation carriers.

VGF nerve growth factor inducible (VGF) was significantly decreased in CSF from all symptomatic groups (peptide sequence THLGEALAPLSK, AS-*GRN*: 67.0%, P=0.0633; S-NC: 62.5%, P=0.025; S-*GRN*: 63.6%, P=0.041; S-*C9orf72*: 58.5%, P=0.018; S-*MAPT*: 51.3%, P=0.002) (Fig. 5B). VGF was highly decreased in the disease-affected brain regions in sporadic FTD (S-NC, IOG: 92.8%, P=0.790; MTG: 43.8%, P=0.043; SFG: 44.0%, P=0.002) and in S-*GRN* (IOG: 72.3%, P=0.170; MTG: 53.2%, P=0.084; SFG: 29.0%, P<0.001) (Supplemental Fig. 5B).

Chromogranin A (CHGA) in CSF was reduced for all symptomatic groups except sporadic FTD (peptide sequence RPEDQELESLSAIEAELEK, AS-*GRN*: 67.9%, P=0.055; S-NC: 72.3%, P=0.095; S-*GRN*: 64.4%, P=0.033; S-*C9orf72*: 48.4%, P<0.001; S-*MAPT*: 54.7%, P=0.002) (Fig. 5C). No significant changes for CHGA were detected in the patient brain samples (Supplemental Fig. 5C).

In contrast to the reduction of the synaptic biomarkers described above, YWHAZ (also named 14-3-3ζ) in CSF was significantly increased in sporadic FTD (peptide sequence TAFDEAIAELDTLSEESYK, AS-*GRN*:109.1%, P=0.650; S-*GRN*: 131.2%, P=0.166; S-NC: 201.6%, P<0.001; S-*C9orf72*: 104.0%, P=0.848; S-*MAPT*: 105.2%, P=0.792) (Fig. 5D). YWHAZ was decreased in the SFG from S-*GRN* relative to controls (IOG: 88.8%, P=0.756; MTG: 67.5%, P=0.334; SFG: 46.7%, P=0.019) (Supplemental Fig. 5D).

### Correlation of biomarkers with disease severity

We evaluated the correlation of key CSF and plasma biomarkers with disease severity (CDR+NACC FTLD□SB) for each form of FTD (CDR+NACC FTLD□Global > 0 from each subject included in the analysis) (Supplemental Table 2). A significant correlation of CSF NfL and CDR+NACC FTLD□SB was observed in symptomatic *C9orf72* and *MAPT*, but not in symptomatic *GRN* and non-carriers (Fig. 6A). Although the correlation did not reach statistical significance for symptomatic *GRN*, a concentration cutoff of ∼2000 pg/mL of NfL in CSF seems to separate patients with relatively mild disease (CDR+NACC FTLD□SB < 5) from patients with more advanced disease (CDR+NACC FTLD□SB > 5) (Fig. 6A). Plasma NfL and CDR+NACC FTLD□SB were positively correlated for all symptomatic genetic groups, but not for sporadic FTD (Fig. 6B).

**Figure 6:**
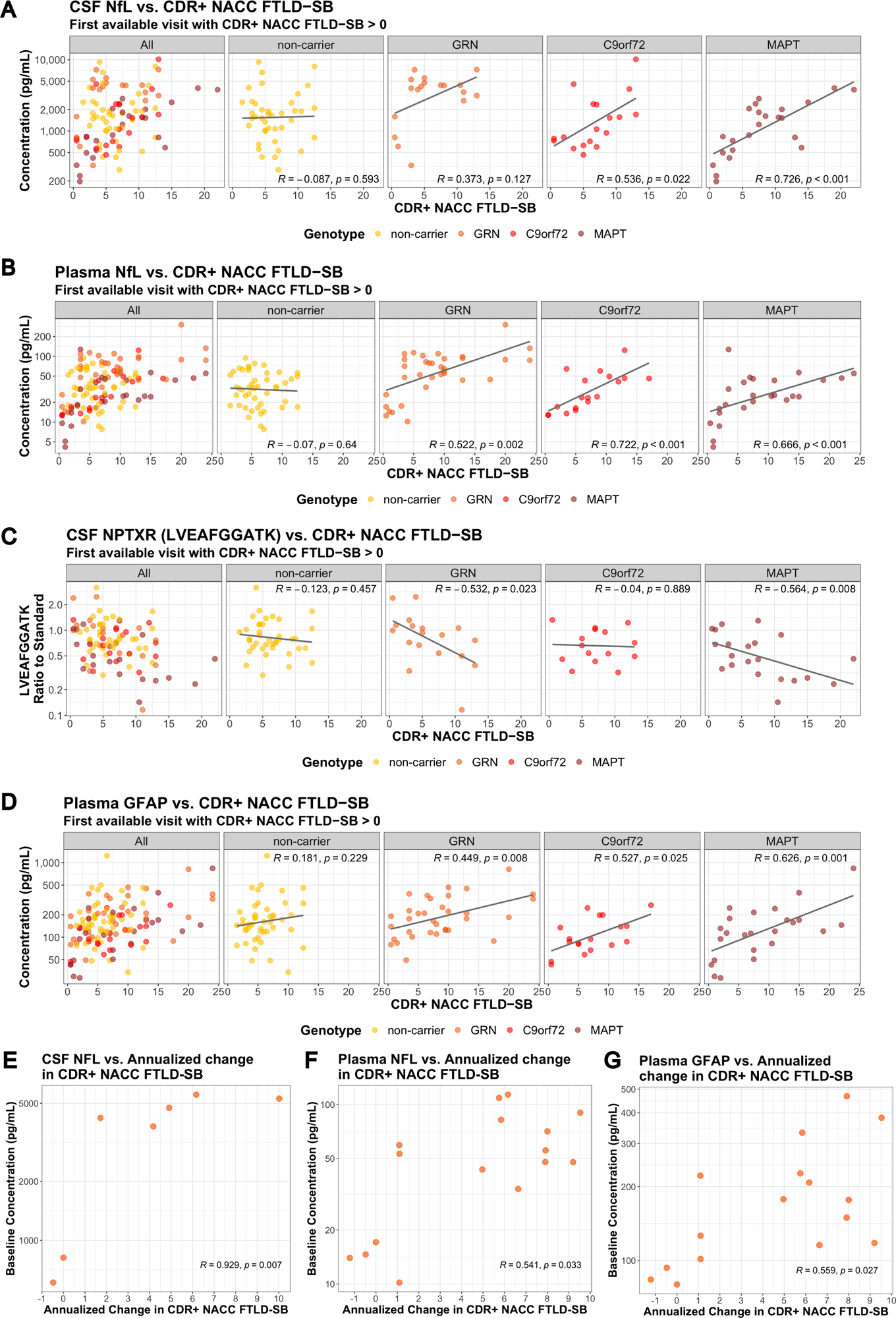
Correlation of NfL, NPTXR and GFAP with clinical measures, and correlation of baseline NfL with the rate of disease progression. Correlation of biomarkers and CDR+NACC FTLD□SB (A) CSF NfL. (B) Plasma NfL. (C) CSF NPTXR (peptide sequence: LVEAFGGATK). (D) Plasma GFAP. In *GRN* mutation carriers, correlation of key biomarkers at the first available visit and annualized change in CDR+NACC FTLD□SB (E) baseline CSF NfL, (F) baseline plasma NfL, and (G) baseline plasma GFAP. Spearman’s correlation coefficients and associated p-values are shown for each patient group. Only patients with CDR+NACC FTLD□SB > 0 at the first available visit are included. For (A)-(D), a fitted line was incorporated into the scatter plot, created using a linear regression model, to provide a visual representation of the overall data trend.

Synaptic biomarkers NPTXR, NPTX1, NPTX2 and VGF in CSF were significantly negatively correlated with CDR+NACC FTLD□SB in symptomatic *GRN* and *MAPT*, but not in symptomatic *C9orf72* and non-carriers (Fig. 6C; Supplemental Table 2). Astroglial biomarker GFAP in plasma was significantly positively correlated with CDR+NACC FTLD□SB for all symptomatic genetic groups, but not for non-carriers (Fig. 6D; Supplemental Table 2).

Lysosomal biomarker BMP(22:6/22:6) in CSF was significantly negatively correlated with CDR+NACC FTLD□SB in symptomatic *GRN* mutation carriers (Supplemental Table 2).

### Longitudinal variations of CSF and plasma biomarkers in *GRN* mutation carriers

Within this dataset, 12 participants and 32 participants with *GRN* mutations had longitudinal CSF and plasma data, respectively, and the corresponding CDR+NACC FTLD□SB scores for those timepoints. Among them, 6 subjects were defined as Converters, who had CDR+NACC FTLD-SB scores changed from 0 to >0 during their longitudinal sample collection.

NfL in CSF and plasma, and GFAP in plasma showed small increases over time as measured by their mean annual change (95% confidence interval) and the highest mean annual changes were observed in Converters (CSF NfL: AS-*GRN* 13.8% (6.0%, 22.2%), S-*GRN* 11.1% (1.6%, 21.6%); plasma NfL: AS-*GRN* 4.6% (-0.3%, 9.7%), S-*GRN* 6.6% (-6.0%, 20.7%), Converters 21.4% (-5.0%, 40.3%); plasma GFAP: AS-*GRN* 4.5% (-3.5%, 13.2%), S-*GRN* 7.1% (-0.2%, 14.9%), Converters 9.0% (-6.7%, 27.4%)) (Supplemental Fig. 6A-C). CSF YKL40 and plasma GlcSph remain relatively stable over time (CSF YKL40: AS-*GRN* 0.8% (-4.7%, 6.6%), S-*GRN* 1.8% (-3.0%, 6.9%); plasma GlcSph: AS-*GRN* -5.7% (-10.5%, -0.8%), S-*GRN* -0.9% (-10.3%, 9.5%), Converters 6.8% (-5.9%, 21.2%)) (Supplemental Fig. 6D-E).

In addition, we found baseline NfL in CSF and plasma, and GFAP in plasma correlated positively with the rate of disease progression as measured by the annualized change in CDR+NACC FTLD□SB (Spearman’s correlation coefficients r; baseline CSF NfL: R=0.929, P=0.007; baseline plasma NfL: R=0.541, P=0.033; baseline plasma GFAP: R=0.559, P=0.027) (Fig. 6E-F).

## Discussion

In this study, we replicated data for key biomarkers in biofluid from patients with FTD that have been reported in the literature with an independent cohort of samples. We also investigated the directionality of changes for key biomarkers in blood, CSF and brain tissues from sporadic FTD and symptomatic *GRN* mutation carriers. Our data highlights the complexity of translating molecular changes in specific tissues to reliable and accessible biomarkers in biofluids, underscoring the importance of understanding the underlying biological implications and factors influencing biomarker transport, release, and detection in biofluids.

### Lysosomal biomarkers

Previous investigations with *Grn* knockout (KO) mice and *in vitro* cellular models demonstrated the impact of PGRN deficiency on lysosomal function and associated biomarkers [^20^]. Our knowledge is now further strengthened by the notable elevation of lysosomal metabolite biomarkers such as GlcSph, GM2, and GB3 in disease-affected brain regions in symptomatic *GRN* mutation carriers. GlcSph is a well-established lysosomal biomarker in Gaucher’s disease, where it accumulates due to defective mutations in GCase [^35^, ^36^]. In the *Grn* KO mice, elevated GlcSph levels in brain, liver and plasma were corrected by a brain penetrant PGRN replacement therapy enabled by a protein transport vehicle (PTV-PGRN) [^20^]. Plasma GlcSph was increased only in *GRN* mutation carriers, not in other symptomatic genetic mutation groups or non-carriers, which is consistent with other publications [^20^, ^37^]. Significant increases of GlcSph were also detected in the diseases affected brain regions from symptomatic GRN mutation carriers, while no significant differences in GlcSph in CSF were observed for symptomatic *GRN* mutation carriers. The absence of significant changes of GlcSph in CSF could be attributed to various factors. This may include inherently low levels of GlcSph in CSF [^38^], confined brain regions with elevated levels leading to dilution of contribution to CSF by other brain regions. Technical considerations may contribute like GlcSph may not be stable over the ∼3-7 years of sample storage time.

GM2 and GB3 are key indicators of lysosomal activity and abnormal accumulation of GM2 and GB3 are often found in lysosomal storage disorders [^39^, ^40^]. The prominent increases of GM2 and GB3 in disease-affected brain tissues from symptomatic *GRN* mutation carriers further support the hypothesis of PGRN mediated lysosomal dysregulation in the pathogenesis of FTD. PGRN deficiency in lysosomes leads to gangliosidosis, which may contribute to neuroinflammation, and neurodegeneration susceptibility observed in FTLD [^21^]. Interestingly, elevated GM2 and GB3 in the brain tissues from sporadic FTD with FTLD-TDP type A pathology suggests that the role of lysosomal dysfunction in neurodegeneration is not limited to *GRN* mutations [^16^]. Albeit the prominent increases of GM2 and GB3 in brain tissues, no significant changes of GM2 were detected in CSF from patients with sporadic FTD or *GRN* mutation carriers and GB3 levels in CSF is below the detection limit of current assay. This suggests a potential compartmentalization or limited translocation of these molecules between brain tissues and the CSF, posing challenges to identify biomarkers in biofluid that closely related to changes in the diseases affected tissues.

While previous research in homozygous *Grn* KO mice revealed significant decrease of BMP levels in brain, signifying lysosomal dysfunction due to the loss of PGRN [^20^, ^21^], our current study observed only a trend of reduction of BMP in CSF and a trend of increases in disease-affected brain regions from symptomatic *GRN* mutation carriers. This disconnect could be attributed to the complete loss of PGRN protein in *Grn* KO mice, potentially yielding a greater impact on BMP levels, in contrast to the heterozygous mutation in FTD patients with *GRN* mutation. Other potential factors for the disconnect could be that the TDP-43 pathology or other physiological features in the terminal stage of the FTLD in human patients could were not recapitulated in mouse models. It is worth noting that a significant reduction of BMP in CSF was observed for symptomatic as compared to the asymptomatic *GRN* mutation carriers, which could indicate an underlying lysosomal defect that aggravate as disease progression and indeed, we observed a significant negative correlation of CSF BMP(22:6/22:6) levels with CDR+NACC FTLD□SB in GRN carriers (Supplemental table 2).

Intriguingly, we observed more pronounced reduction of CSF BMP (22:6/22:6) in symptomatic *MAPT* mutation carriers, which implies lysosomal dysfunction is also present in primary tauopathies as suggested previously [^14^]. Indeed, accumulation of autophagic and lysosomal markers in human brain tissue from patients with primary tauopathies have been reported suggesting a defect of the autophagosome-lysosome pathway that may contribute to the development of tau pathology [^41^]. Tau seeds isolated from the brain of Alzheimer’s Disease induced lysosomal stress, and inhibition of lysosomal enzyme GCase contributed to the accumulation of tau in human primary fibroblasts [^42^]. Further investigation of the mechanistic link of tau pathology and lysosomal dysregulation, and exploration of biomarker opportunities in various tauopathies such as progressive supranuclear palsy (PSP) will be warranted.

### Glial biomarkers

GFAP and YKL40 have emerged as pivotal glial biomarkers in neurodegenerative diseases [^43^, ^44^]. GFAP in plasma and YKL40 in CSF were highly increased in symptomatic *GRN*, *MAPT* and non-carrier groups, but not in *C9orf72,* indicating specific genetic mutations might impact glial cell activation and their responses to neuropathology. PGRN has been shown to regulate innate immune response and inflammation, and loss of PGRN leads to astrogliosis and microgliosis in mouse models [^45^, ^46^, ^20^, ^10^]. Plasma GFAP levels have been shown to be correlated with multiple indicators of FTD disease severity, including cortical thickness and global disease severity as measured by the Frontotemporal Dementia Rating Scale (FRS) and CDR+NACC FTLD□SB [^44^, ^47^]. High plasma GFAP levels predicted future temporal lobe atrophy, declines in cognitive function, and decreased survival in longitudinal analyses [^48^, ^49^, ^47^]. Consistent with these studies, our study showed that plasma GFAP levels are positively correlated with CDR+NACC FTLD□SB for symptomatic *GRN*, *C9orf72* and *MAPT* mutation carriers. We further demonstrated that plasma GFAP positively correlates with the rate of disease progression in symptomatic GRN mutation carriers. To our knowledge, the prognostic correlation of baseline plasma GFAP levels to the rate of disease progression is a novel finding for S-*GRN*.

In addition to YKL40 increases in CSF, our data showed YKL40 is highly increased in brain tissues of symptomatic *GRN* mutation carriers, aligning with previous findings [^43^, ^50^, ^51^]. Notably, one study reported no significant increases in YKL40 in brain samples from patients with FTD, despite observing elevated levels in paired premortem CSF [^52^], but these subjects also had different underlying genotypes or had a different clinical diagnosis (FTLD-Tau; FTLD with amyotrophic lateral sclerosis, FTLD-ALS; and dementia with Lewy body, DLB). The precise role of YKL40 in FTD pathogenesis and its correlation with clinical metrics remain insufficiently characterized.

### Neurodegenerative and synaptic biomarkers

NfL is a well-known clinical biomarker for many neurodegenerative diseases such as amyotrophic lateral sclerosis (ALS), FTD and spinal muscular atrophy (SMA). Consistent with previous publications, our data showed NfL in CSF and plasma are increased in patients with all forms of FTD with the highest elevation observed in symptomatic *GRN* mutation carriers [^53^, ^23^, ^32^, ^33^], and baseline levels correlate positively with the rate of disease progression [^23^]. In patients with dementia, the highest NfL levels are observed in patients with FTD as compared to other types of dementia such as AD, and NfL levels correlate with disease severity and survival, regardless of clinical diagnosis [^54^, ^23^]. NfL levels rise early in the disease course and predict the onset of symptoms in presymptomatic genetic carriers [^55^, ^56^, ^57^, ^32^, ^34^]. In symptomatic patients, high NfL levels predict multiple indicators of disease progression, including brain atrophy, declines in cognitive function as measured by neuropsychological evaluation, worsening disease severity as measured by the CDR+NACC FTLD□SB, and shorter survival time [^58^, ^56^, ^32^]. In preclinical studies with *Grn* KO mice, a protein replacement strategy by engineering protein transport vehicle (PTV):PGRN decreased NfL levels in CSF [^20^]. Taken together, NfL will continue to be an important biomarker in clinical trials for patient stratification and drug treatment responses, especially given the precedence that was recently set by the FDA’s accelerated approval of Qalsody (tofersen) to treat patients with amyotrophic lateral sclerosis (ALS) associated with a mutation in the superoxide dismutase 1 (SOD1) gene (SOD1-ALS) mainly based on significant decreases in blood NfL levels.

UCHL1 increases in CSF and plasma have been demonstrated in acute neuronal injury and neurodegenerative diseases such as AD and ALS [^59^, ^60^]. UCHL1 in blood has been approved by the FDA as a prognostic biomarker for mild traumatic brain injury (TBI) [^61^, ^59^]. Notably, our study is the first to demonstrate the increases of UCHL1 in the CSF and decreases in SFG from symptomatic *GRN* mutation carriers. UCHL1 functions as a deubiquitinating enzyme and a ubiquitin ligase and plays an important role in the ubiquitin-proteasome pathway (UPP). UCHL1 loss of function mutations in humans have been associated with early onset neurodegeneration [^62^]. The reduction of UCHL1 in brain tissues from symptomatic *GRN* might reflect an overall loss in neurons, but it might also play a pathogenic role.

Our data confirmed that synaptic biomarker neuropentraxins and receptors (NPTX1, NPTX2, and receptor NPTXR) were decreased in all forms of FTD, particularly in symptomatic *C9orf72* and *MAPT* mutation carriers [^63^]. They have also been shown to be decreased in patients with AD and are potential biomarkers for AD progression [^64^]. Our study showed that they were correlated with disease severity, supporting its prognostic value for disease progression in FTD. In addition, other synaptic biomarkers VGF, CHGA and YWHAZ were also altered in CSF from symptomatic genetic mutations and non-carriers, consistent with previous findings [^65^, ^32^, ^50^]. More studies will be needed to further understand the biological and clinical implication of these findings. In addition, these synaptic biomarkers were altered in the brain tissues from symptomatic *GRN* mutation carriers and non-carriers, particularly in disease affected brain regions, indicating that synaptic dysregulation could be a key pathophysiological mechanism of FTD.

### The relationship of biomarker changes in biofluid and postmortem brain tissues

Studying the relationship between biomarkers in biofluid and disease-affected tissues is important for accurate biomarker data interpretation and gaining a deeper understanding of the biological mechanisms underlying these markers. This has been best showcased in Alzheimer’s disease where accumulation of amyloid plaques and Tau tangles in brain tissues correlate with the reduction of Abeta peptides and increase of Tau (pTau) in biofluids. However, such studies are relatively scarce in FTD. To the best of our knowledge, our study represents a pioneering effort in measuring key biomarker changes across different matrices (CSF, blood, and brain tissues), despite the caveat that antemortem biofluid samples were not derived from the same subjects as postmortem brain tissues. Our data reveals a nuanced and biomarker-specific relationship between changes in biofluid and tissue biomarkers, shedding light on the complexity of these interactions. Notably, robust biomarker changes observed in brain tissues, such as GB3 and GM2, do not consistently translate into discernible changes in biofluid levels. For some biomarkers such as NfL and UCHL-1, the directional changes in brain tissues are not the same as the alterations observed in biofluids. Biofluid biomarker levels appear to reflect a dynamic equilibrium shaped by the interplay of biomarker diffusion from tissues and degradation/turnover in circulation. In contrast, biomarker levels in tissues provide a snapshot of the accumulative status at the terminal stage of the disease. Additionally, biomarker assays may exhibit varying detection sensitivity and specificity for analytes in different matrices, adding an additional layer of complexity to interpretation.

### Biomarker changes in asymptomatic *GRN* mutation carriers

We found significant elevation of GlcSph & GFAP in plasma, and a significant decrease in CSF NPTXR and CHGA in asymptomatic *GRN* mutation carriers as compared to non-carrier controls. We also found NfL increases in CSF and plasma in asymptomatic *GRN* mutation carriers, consistent with previous published data that showed elevated plasma or serum NfL levels in asymptomatic (or pre-symptomatic) *GRN, C9orf72 or MAPT* [^55^, ^55^, ^57^, ^34^, ^33^, ^23^]. For potential prevention studies, it would be critical to identify pre-symptomatic carriers that may convert to symptomatic phases in the timeframe of clinical studies. Biomarkers, including NfL and others that were confirmed or newly identified in our studies, might be used together to model disease age and to identify potential subjects close to their disease onset [^34^].

### Limitations

The present study is subject to several limitations. While the multi-center design of the ALLFTD study enabled the collection of CSF and plasma samples from a sizable cohort of genetic and sporadic FTD cases in the United States, the inherent challenge lies in the relatively modest sample size within each group. This constraint may limit the robustness of statistical analyses, particularly in correlation assessments with clinical measures. The sporadic FTD group contained a mixture of diagnoses and underlying pathologies, including likely some cases with AD since 50% were CBS. This limits the ability to draw conclusions from this group. Furthermore, the number of subjects with longitudinal sample collection is small, limiting our ability to comprehensively assess biomarker changes throughout the progression of the disease. The retrospective collection of samples, stored in biobanks for varying durations, could potentially influence biomarker assessments due to uncertainties of their long-term storage stability. Notably, the brain tissue samples exhibit limitations in sample size, and the healthy control samples were obtained from a different source.

## Conclusions

Taken together, the biomarker data presented here offers valuable insights into the pathogenesis of FTD, particularly FTD associated with *GRN* mutations, and could serve as key indicators for assessing drug treatment effects in clinical trials. Future work could focus on assessing more longitudinal biofluid samples, especially in the *C9orf72* and *MAPT* mutation groups. Of particular interest will be collecting more data for asymptomatic genetic carriers, especially in the few years before and after symptom onset. Lastly, it would be of interest to conduct analysis in matched ante-mortem biofluid and post-mortem brain samples from the same subjects to investigate the relationships between brain and biofluid of specific analytes.

## Abbreviations

FTD: Frontotemporal Dementia
ALS: Amyotrophic Lateral Sclerosis
MTG: middle temporal gyrus
SFG: superior frontal gyrus
IOG: inferior occipital gyrus
GlcSph: glucosylsphingosine
TDP-43: transactive response DNA-binding protein 43
FTLD: frontotemporal lobar degeneration
CNS: central nervous system
BMP: lysosomal phospholipid bis(monoacylglycero)phosphate
Gcase: glucocerebrosidase
bvFTD: behavioral variant FTD
CBS: corticobasal syndrome
PPA: primary progressive aphasia
NfL: neurofilament light chain
CSF: cerebrospinal fluid
KO: knockout
SMA: spinal muscular atrophy

## Disclosure statement

J.H., C-L.C., R.R., B.T., M. F., N.S.G., F. M., S.S.D., J.H.S., J.A., A.B., R.T., G.D.P., A.B., F.H. are current employees and shareholders of Denali Therapeutics; B.V. is a former intern of Denali Therapeutics. C.A.P is a former contractor of Denali Therapeutics. J.D., P.A., K.S.L., are former employees and shareholders of Denali Therapeutics.

A.L.B served as a paid consultant to AGTC, Alector, Alzprotect, Amylyx, Arkuda, Arrowhead, Arvinas, Aviado, Eli Lilly, GSK, Humana, Merck, Modalis, Muna, Oligomerix, Oscotec, Pfizer, Roche, Switch, Transposon and UnlearnAI.

L.V. is the site PI for Biogen-sponsored clinical trials and has consulted for Retrotope.

B.F.B. is an investigator for clinical trials sponsored by Biogen, Alector, EIP Pharma, Cognition Therapeutics, and Transposon; a Scientific Advisory Board member for the Tau Consortium, AFTD, LBDA, and GE HealthCare; and serves as a member of the Data Safety Monitoring Board for a clinical trial on mesenchymal stem cells in MSA.

No other disclosures were reported.

## Funding resources

This work was funded by Denali Therapeutics.

ALLFTD is funded by NIH grant number U19AG063911.

A.L.B receives funds from NIH grants (R01AG038791, P01AG019724) and Bluefield Project to Cure FTD and Rainwater Charitable Foundation.

L.V. receives funds from NIH K23AG073514.

## Author contributions

F.H, G.D.P, L.V., A.B. A.L.B conceived and designed the study. J.H., R. R., B.V, J.D., B.T., M.F., P. A., N.S.G., F.M., C.A.P., S.S.D, J.A., analyzed the samples and generated the data. J.H., C-L. C., F. H. A.B. analyzed the data. H. W. H, L.V, A. L. L, W.W.S. provided support for human samples. J.H. C-L. C. and F.H. authored the manuscript with input from all the authors. F.H, A.B, A.L.B, G.D.P, L.V., J.H., C-L.C., R.T., A. B., K.S.L., W.S., J.H.S. P.A., N.S.G. provided data interpretation and manuscript editing. ALLFTD investigators including A.L.B, B.F.B, H.J.R collected the clinical data and biomarker samples. All authors read and approved the final manuscript.

## Acknowledgements

- CSF and plasma samples from the National Centralized Repository for Alzheimer’s Disease and Related Dementias (NCRAD), which receives government support under a cooperative agreement grant (U24 AG021886) awarded by the National Institute on Aging (NIA), were used in this study.
- Data collection and dissemination of the data presented in this manuscript was supported by the ALLFTD Consortium (U19: AG063911, funded by the National Institute on Aging and the National Institute of Neurological Diseases and Stroke) and the former ARTFL & LEFFTDS Consortia (ARTFL: U54 NS092089, funded by the National Institute of Neurological Diseases and Stroke and National Center for Advancing Translational Sciences; LEFFTDS: U01 AG045390, funded by the National Institute on Aging and the National Institute of Neurological Diseases and Stroke). The authors acknowledge the invaluable contributions of the study participants and families as well as the assistance of the support staffs at each of the participating sites.
- Human brain tissue samples were provided by the Neurodegenerative Disease Brain Bank at the University of California, San Francisco, which receives support from NIH grants P30AG062422, P01AG019724, U01AG057195, and U19AG063911, as well as the Rainwater Charitable Foundation and the Bluefield Project to Cure FTD.
- We thank KCAS Bioanalytical Services (Olathe, KS) for measuring GlcSph in CSF.
- We thank Drs. Dolores Diaz, René Meisner, Todd Logan, Yuda Zhu, Fabian Model, Joseph Lewcock for their valuable scientific inputs on the data interpretation and manuscript and Dr. Paul Benton, Catherine Bedard and Eric Liang for their technical support.

