## Supplemental materials for "Alterations in Lysosomal, Glial and Neurodegenerative Biomarkers in Patients with Sporadic and Genetic Forms of Frontotemporal Dementia"

#### Figure Legends

**Supplemental Figure 1**

(A) Plasma progranulin (B) CSF progranulin levels in AS-NC, AS-*GRN*, S-NC, S-*GRN*, S-*C9orf72* and S-*MAPT*. (C) Brain progranulin levels in IOG, MTG and SFG from AS-NC Control, S-NC and S-*GRN*. For CSF/plasma, each dot represents an individual subject visit. For brain, each dot represents an individual subject. Box plots are median ± interquartile range (IQR). *p<0.05, **p<0.01, ***p<0.001. AS-NC: asymptomatic non-carrier controls; AS-*GRN*: asymptomatic *GRN* mutation carriers; S-NC: symptomatic non-carriers or sporadic FTD; S-*GRN*: symptomatic *GRN* mutation carriers; S-*C9orf72*: symptomatic *C9orf72* mutation carriers; S-*MAPT*: symptomatic *MAPT* mutation carriers; IOG: inferior occipital gyrus; MTG: medial temporal gyrus; SFG: superior frontal gyrus.

**Supplemental Figure 2**

(A) BMP (22:6/22:6) in IOG, MTG and SFG from AS-NC, S-NC and S-*GRN*. (B) Plasma BMP (22:6/22:6) in AS-NC controls, AS-*GRN*, S-NC, S-*GRN*, S-*C9orf72* and S-*MAPT*. For CSF/plasma, each dot represents an individual subject visit. For brain, each dot represents an individual subject. For brain, each dot represents an individual subject visit. Box plots are median ± interquartile range (IQR). Statistical analysis was performed in comparison to non-carrier healthy controls, *p<0.05, **p<0.01. AS-NC: asymptomatic non-carrier controls; AS-*GRN*: asymptomatic *GRN* mutation carriers; S-NC: symptomatic non-carriers or sporadic FTD; S-*GRN*: symptomatic *GRN* mutation carriers; S-*C9orf72*: symptomatic *C9orf72* mutation carriers; S-*MAPT*: symptomatic *MAPT* mutation carriers; IOG: inferior occipital gyrus; MTG: medial temporal gyrus; SFG: superior frontal gyrus.

**Supplemental Figure 3**

(A) Plasma YKL40 levels in AS-NC, AS-*GRN*, S-NC, S-*GRN*, S-*C9orf72* and S-*MAPT*. Each dot represents an individual subject visit. Box plots are median ± interquartile range (IQR). AS-NC: asymptomatic non-carrier controls; AS-*GRN*: asymptomatic *GRN* mutation carriers; S-NC: symptomatic non-carriers or sporadic FTD; S-*GRN*: symptomatic *GRN* mutation carriers; S-*C9orf72*: symptomatic *C9orf72* mutation carriers; S-*MAPT*: symptomatic *MAPT* mutation carriers.

**Supplemental Figure 4**

(A) NEFL (peptide sequence: VLEAELLVLR) and (B) UCHL1 (peptide sequence: LGFEDGSVLK) in brain regions of IOG, MTG and SFG from AS-NC Control, S-NC and S-*GRN*. (C) Plasma UCHL1 from AS-NC controls, AS-*GRN*, S-NC, S-*GRN*, S-*C9orf72* and S-*MAPT*. For plasma, each dot represents an individual subject visit. For brain tissues, each dot represents an individual subject. Box plots are median ± interquartile range (IQR). Statistical analysis was performed in comparison to non-carrier healthy controls, *$p<0.05$, **$p<0.01$, ***$p<0.001$ (unadjusted for multiple comparison in the proteomic panel). AS-NC: asymptomatic non-carrier controls; AS-*GRN*: asymptomatic *GRN* mutation carriers; S-NC: symptomatic non-carriers or sporadic FTD; S-*GRN*: symptomatic *GRN* mutation carriers; S-*C9orf72*: symptomatic *C9orf72* mutation carriers; S-*MAPT*: symptomatic *MAPT* mutation carriers; IOG: inferior occipital gyrus; MTG: medial temporal gyrus; SFG: superior frontal gyrus.

**Supplemental Figure 5**

(A) NPTXR (peptide sequence: LVEAFGGATK), (B) VGF (THLGEALAPLSK), (C) CHGA (RPEDQELESLSAIEAELEK) and (D)YWHAZ (TAFDEAIAELDTLSEESYK) in brain regions of IOG, MTG and SFG from AS-NC Control, S-NC and S-*GRN*. Each dot represents an individual subject. Box plots are median ± interquartile range (IQR). Statistical analysis was performed in comparison to non-carrier healthy controls, *p<0.05, **p<0.01, ***p<0.001 (unadjusted for multiple comparison in the proteomic panel). S-NC: symptomatic non-carriers or sporadic FTD; S-*GRN*: symptomatic *GRN* mutation carriers; IOG: inferior occipital gyrus; MTG: medial temporal gyrus; SFG: superior frontal gyrus.

**Supplemental Figure 6**

Longitudinal biomarker levels over time for *GRN* mutation carriers. Displaying mean annual change with corresponding 95% confidence interval. Closed dots are clinically diagnosed with symptomatic FTD and open dots are asymptomatic. (A) CSF NfL (B) Plasma NfL (C) Plasma GFAP (D) CSF YKL40 (E) Plasma GlcSph.

### Supplemental Figure 1

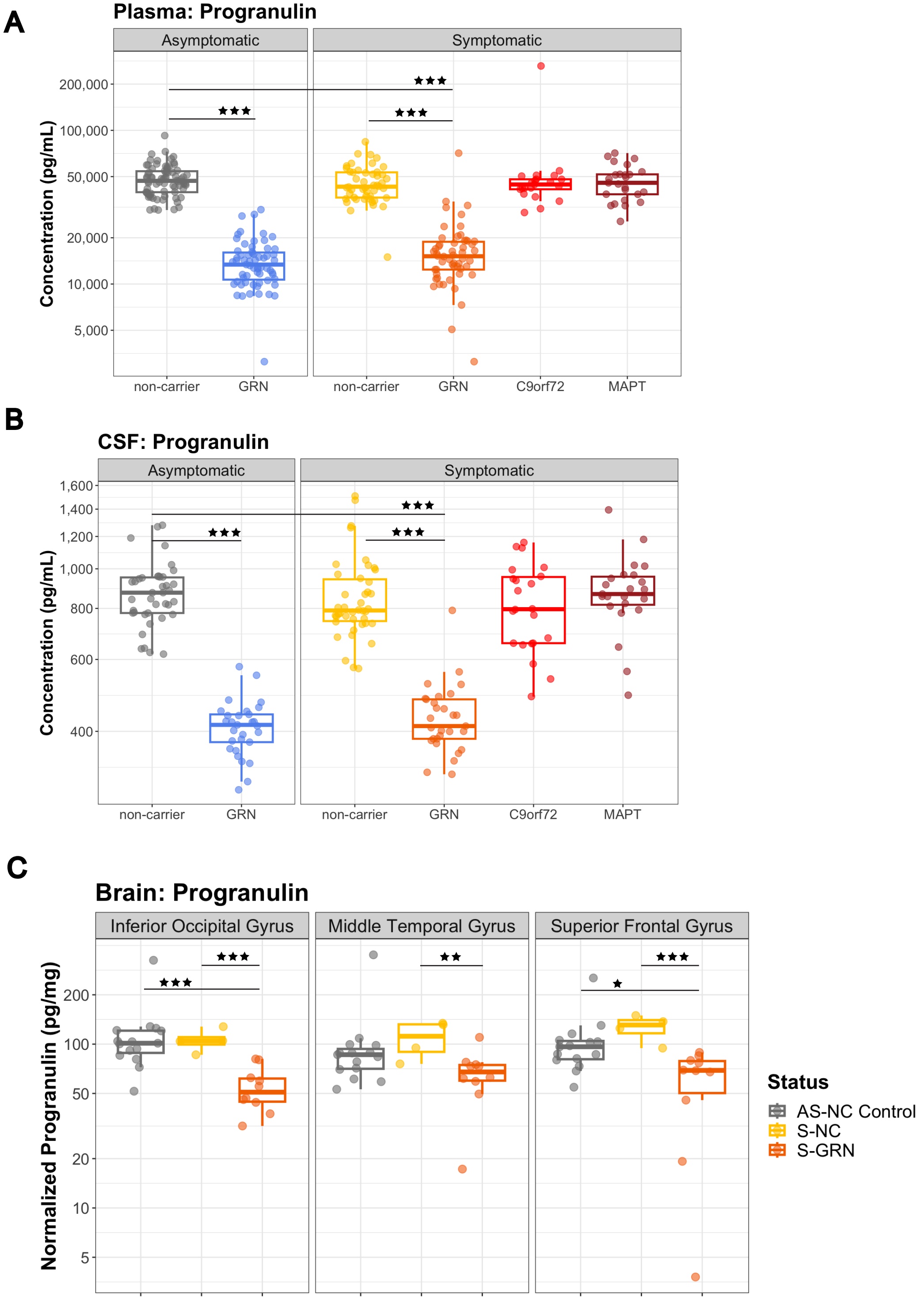

#### Supplemental Figure 2

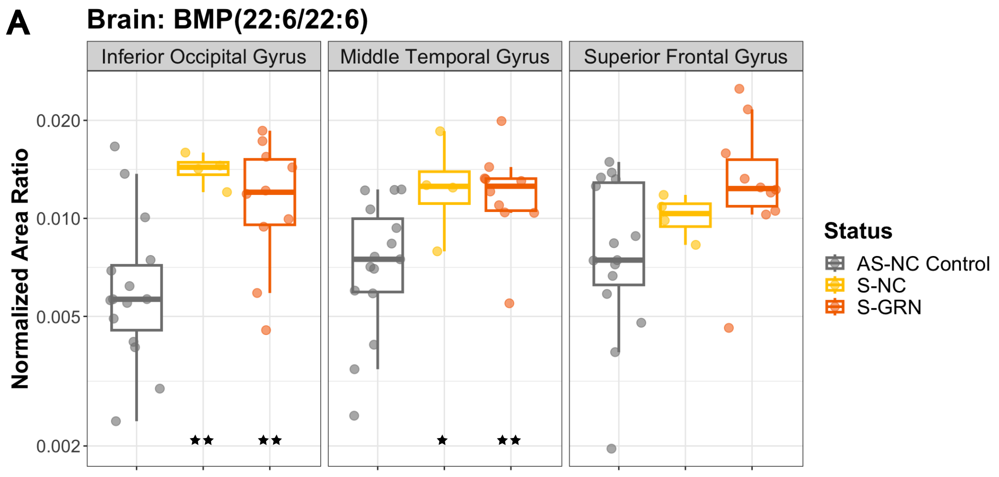


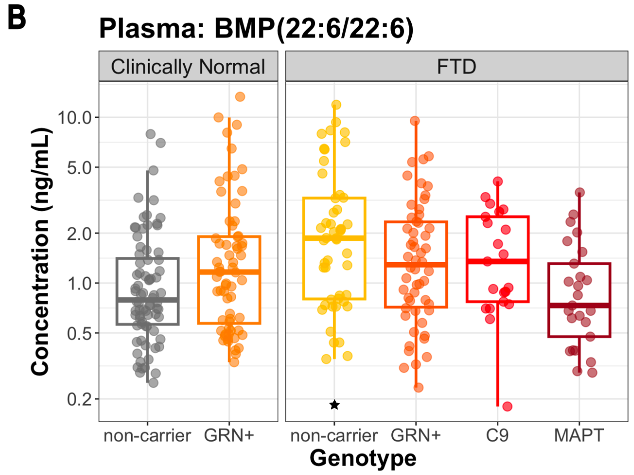

### Supplemental Figure 3

**A**

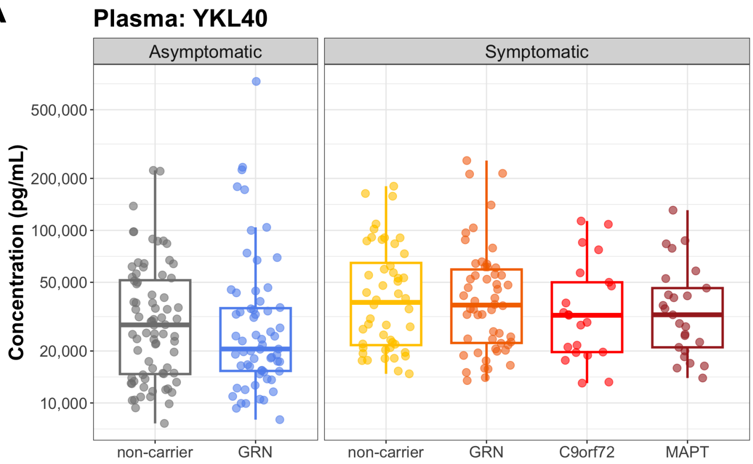

#### Supplemental Figure 4

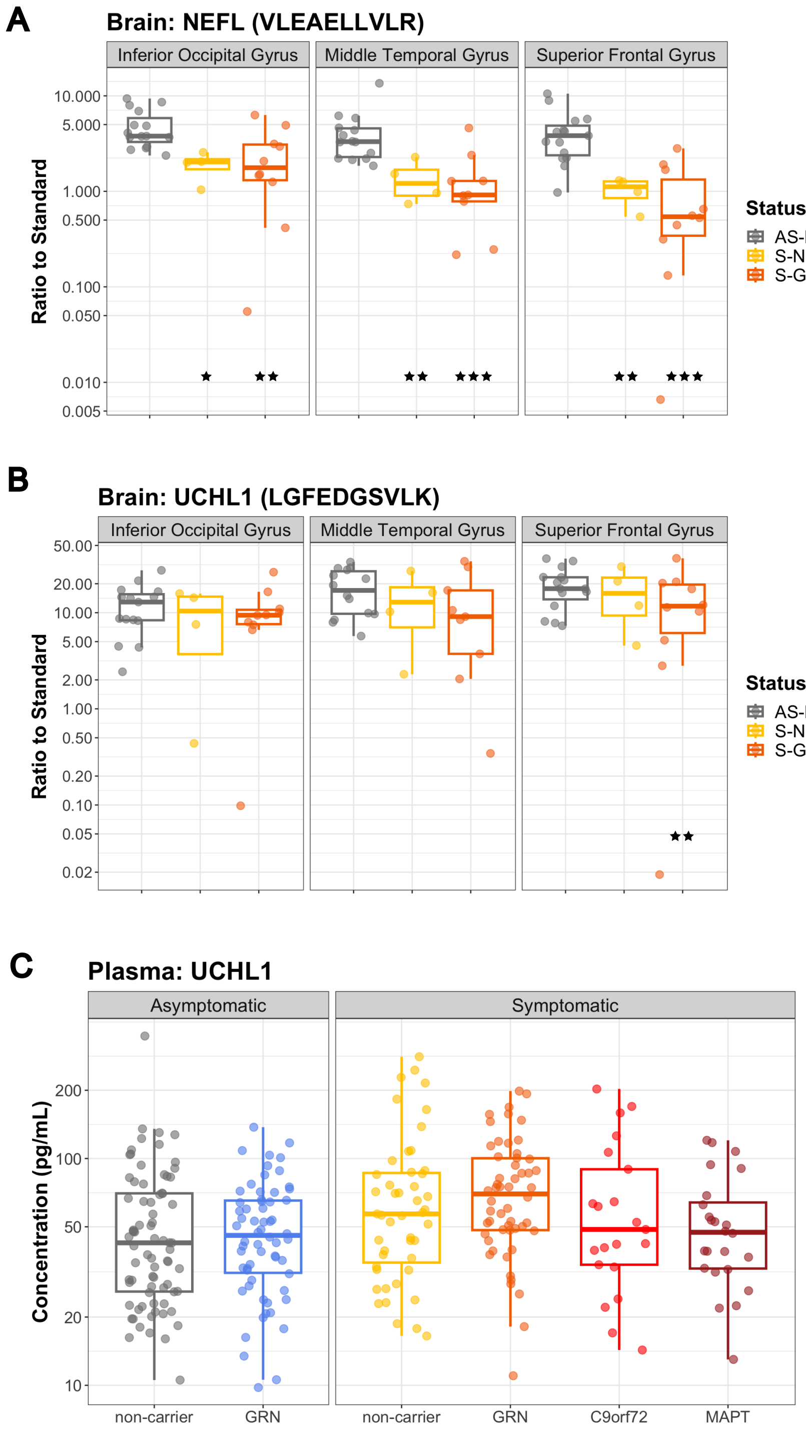

### Supplemental Figure 5

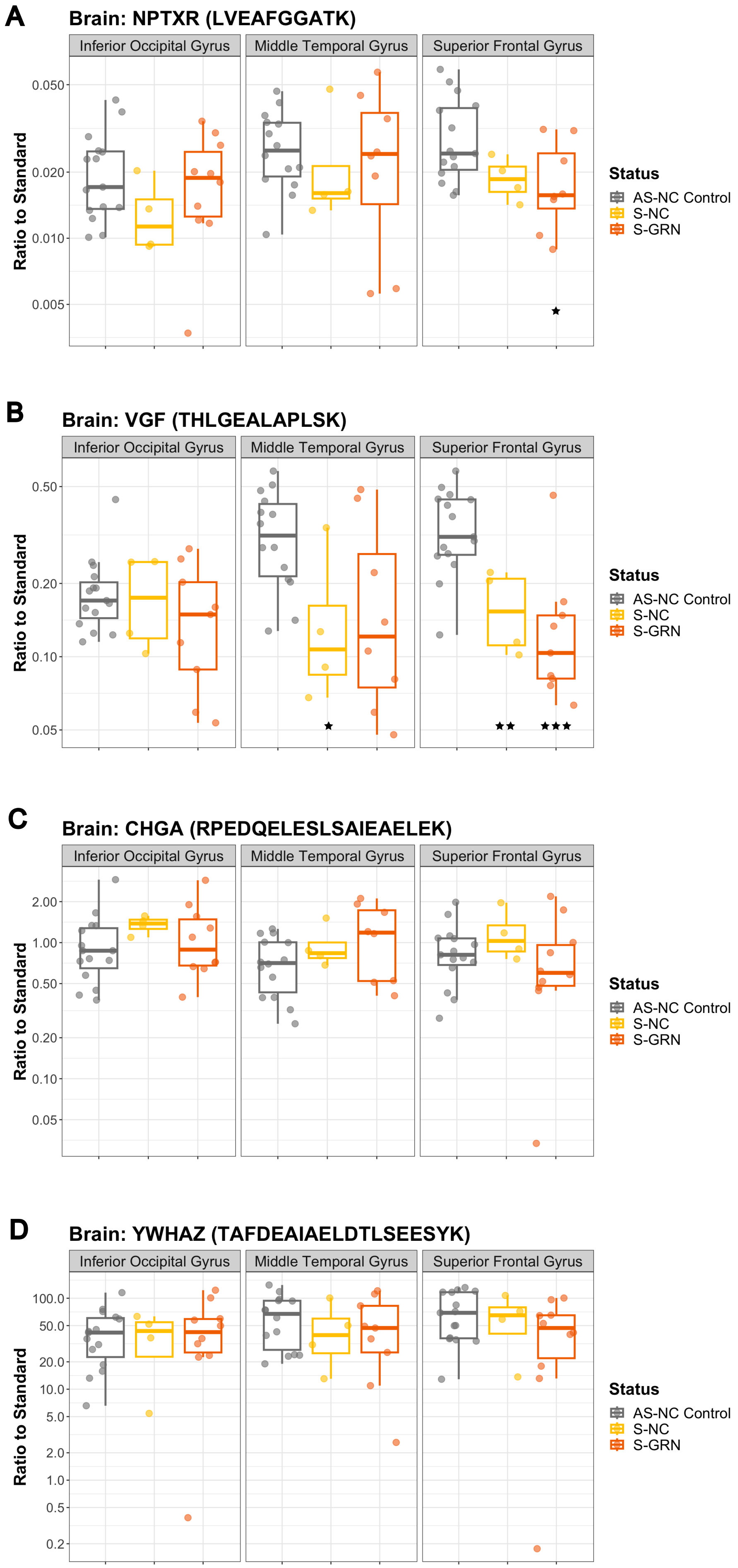

### Supplemental Figure 6

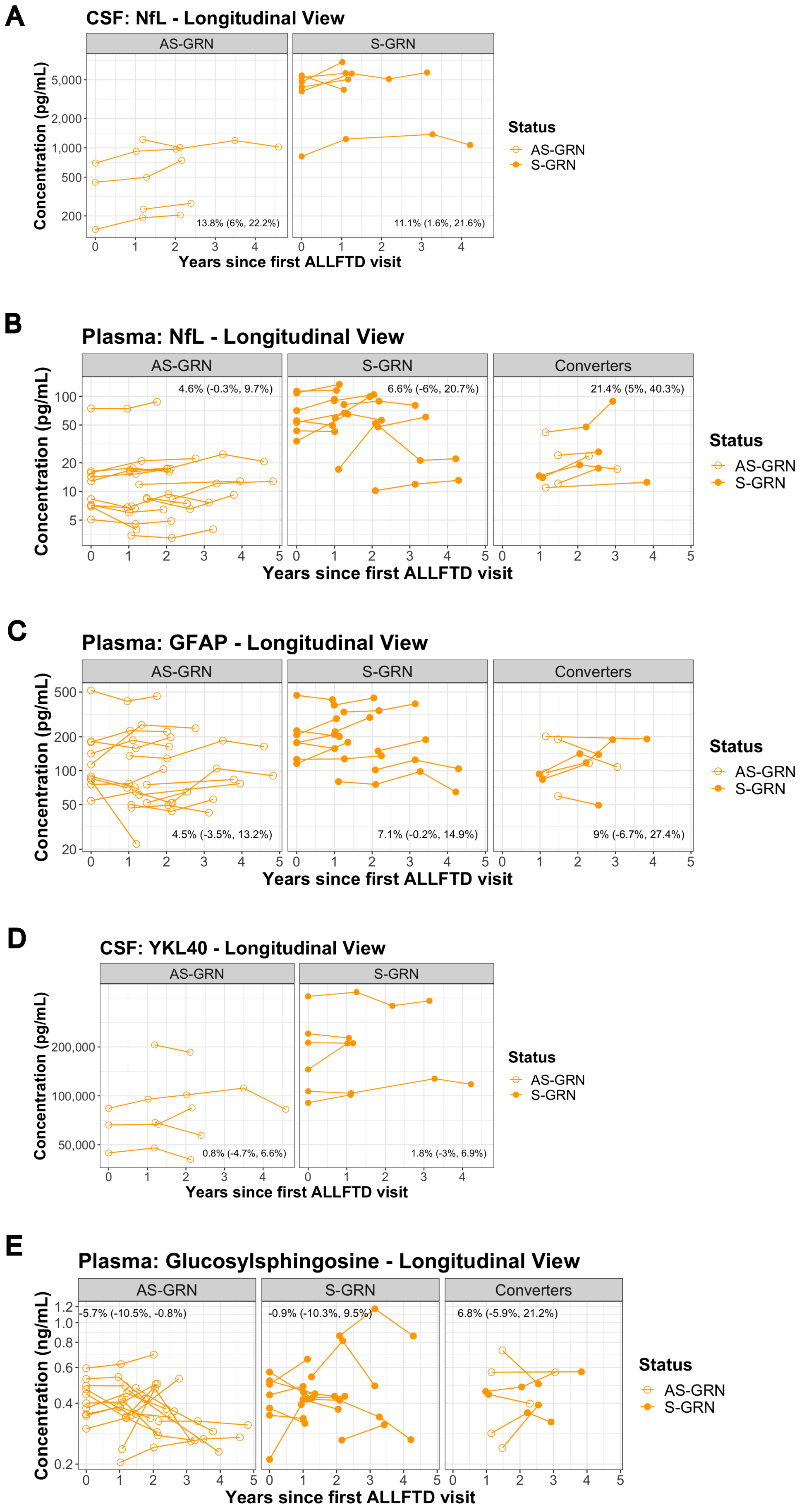

**Supplemental Table 1**

List of protein ID's and peptides in the targeted LC-MS/MS proteomic panel.

| Protein Gene | Peptide | Matrices Detected |
|---|---|---|
| APOB | ILGEELGFASLHDLQLLGK | Brain |
| APOE | LGPLVEQGR | Brain & CSF |
| | VQAAVGTSAAPVPSDNH | Brain & CSF |
| APP | LVFFAEDVGSNK | CSF |
| | THPHFVIPYR | Brain & CSF |
| C8G | SLPVSDSVLSGFEQR | CSF |
| | VQEAHLTEDQIFYFPK | CSF |
| CA1 | GGPFSDSYR | Brain |
| CHGA | EAVEEPSSK | Brain & CSF |
| | RPEDQELESLSAIEAELEK | Brain & CSF |
| | SGELEQEEER | Brain & CSF |
| CHI3L1 | EAGTLAYYEICDFLR | Brain & CSF |
| | LVMGIPTFGR | Brain & CSF |
| | TLLSVGGWNFGSQR | Brain & CSF |
| CLU | ASSIIDELFQDR | Brain & CSF |
| GRN | EVVSAQPATFLAR | CSF |
| HBB | LLGNVLVCVLAHHFGK | Brain |
| IGHA1 | TPLTATLSK | Brain & CSF |
| NEFL | ALYEQEIR | Brain |
| | VLEAELLVLR | Brain & CSF |
| NPTX1 | FQLTFPLR | Brain & CSF |
| | LENLEQYSR | Brain & CSF |
| NPTX2 | LESLEHQLR | CSF |
| | TESTLNALLQR | CSF |
| NPTXR | LVEAFGGATK | Brain & CSF |
| | VAELEHGSSAYSPPDAFK | Brain & CSF |
| PRNP | GENFTETDVK | Brain & CSF |
| | VVEQMCITQYER | Brain & CSF |
| PSAP | EIVDSYLPVILDIIK | Brain & CSF |
| | GCSFLPDPYQK | Brain & CSF |
| PTPRN2 | THTAQDRPPAEGDDR | CSF |

| Protein Gene | Peptide | Matrices Detected |
|---|---|---|
| SNCA | EQVTNVGGAVVTGVTAVAQK | Brain |
| TIMP1 | GFQALGDAADIR | CSF |
| TREM2 | NLQPHDAGLYQCQSLHGSEADTLR | CSF |
| | VLVEVLADPLDHR | Brain & CSF |
| | VVSTHNLWLLSFLR | CSF |
| UCHL1 | LGFEDGSVLK | Brain & CSF |
| | MPFPVNHGASSEDTLLK | Brain |
| VGF | EPVAGDAVPGPK | CSF |
| | THLGEALAPLSK | Brain & CSF |
| VSTM2B | ELLHELALSVPGAR | Brain & CSF |
| YWHAE | LICCDILDVLDK | Brain & CSF |
| YWHAG | DSTLIMQLLR | Brain & CSF |
| | NVTELNEPLSNEER | Brain & CSF |
| | TAFDDAIAELDTLNEDSYK | Brain |
| YWHAZ | SVTEQGAELSNEER | Brain & CSF |
| | TAFDEAIAELDTLSEESYK | Brain & CSF |

**Supplemental Table 2**

Correlation between biomarkers and CDR+NACC FTLD-SB. Spearman's rank correlation coefficients and p-values. Only patients with CDR+NACC FTLD-SB > 0 at the first available visit are included.

| Analyte | Matrix | Genotype | Subject, N | Spearman's Corr. Coef. (Est.) | p-value (unadjusted) |
|---|---|---|---|---|---|
| NFL | CSF | non-carrier | 40 | -0.087 | 0.593 |
| NFL | CSF | GRN | 18 | 0.373 | 0.127 |
| NFL | CSF | C9 | 18 | 0.536 | 0.022 |
| NFL | CSF | MAPT | 23 | 0.726 | 0.000 |
| NFL | Plasma | non-carrier | 47 | -0.070 | 0.640 |
| NFL | Plasma | GRN | 34 | 0.522 | 0.002 |
| NFL | Plasma | C9 | 18 | 0.722 | 0.001 |
| NFL | Plasma | MAPT | 25 | 0.666 | 0.000 |
| UCHL1 | CSF | non-carrier | 41 | 0.019 | 0.905 |
| UCHL1 | CSF | GRN | 18 | -0.030 | 0.906 |
| UCHL1 | CSF | C9 | 18 | 0.056 | 0.825 |
| UCHL1 | CSF | MAPT | 23 | 0.541 | 0.008 |
| GFAP | Plasma | non-carrier | 46 | 0.181 | 0.229 |
| GFAP | Plasma | GRN | 34 | 0.449 | 0.008 |
| GFAP | Plasma | C9 | 18 | 0.527 | 0.025 |
| GFAP | Plasma | MAPT | 24 | 0.626 | 0.001 |
| YKL40 | CSF | non-carrier | 41 | 0.097 | 0.547 |
| YKL40 | CSF | GRN | 18 | 0.201 | 0.424 |
| YKL40 | CSF | C9 | 18 | 0.329 | 0.183 |
| YKL40 | CSF | MAPT | 23 | 0.468 | 0.024 |
| GlcSph | Plasma | non-carrier | 47 | -0.303 | 0.038 |
| GlcSph | Plasma | GRN | 34 | 0.000 | 0.999 |
| GlcSph | Plasma | C9 | 18 | 0.673 | 0.002 |
| GlcSph | Plasma | MAPT | 25 | -0.040 | 0.851 |
| BMP(22:6/22:6) | CSF | non-carrier | 41 | -0.253 | 0.110 |
| BMP(22:6/22:6) | CSF | GRN | 18 | -0.567 | 0.014 |
| BMP(22:6/22:6) | CSF | C9 | 18 | -0.295 | 0.234 |

| Analyte | Matrix | Genotype | Subject, N | Spearman's Corr. Coef. (Est.) | p-value (unadjusted) |
|---|---|---|---|---|---|
| BMP(22:6/22:6) | CSF | MAPT | 23 | -0.046 | 0.835 |
| NPTXR: LVEAFGGATK | CSF | non-carrier | 39 | -0.123 | 0.457 |
| NPTXR: LVEAFGGATK | CSF | GRN | 18 | -0.532 | 0.023 |
| NPTXR: LVEAFGGATK | CSF | C9 | 15 | -0.040 | 0.889 |
| NPTXR: LVEAFGGATK | CSF | MAPT | 21 | -0.564 | 0.008 |
| NPTX2: LESLEHQLR | CSF | non-carrier | 34 | -0.227 | 0.196 |
| NPTX2: LESLEHQLR | CSF | GRN | 16 | -0.640 | 0.008 |
| NPTX2: LESLEHQLR | CSF | C9 | 12 | -0.540 | 0.070 |
| NPTX2: LESLEHQLR | CSF | MAPT | 15 | -0.746 | 0.001 |
| NPTX1: FQLTFPLR | CSF | non-carrier | 38 | -0.107 | 0.523 |
| NPTX1: FQLTFPLR | CSF | GRN | 17 | -0.603 | 0.010 |
| NPTX1: FQLTFPLR | CSF | C9 | 15 | -0.004 | 0.990 |
| NPTX1: FQLTFPLR | CSF | MAPT | 21 | -0.501 | 0.021 |
| VGF: THLGEALAPLSK | CSF | non-carrier | 38 | -0.165 | 0.323 |
| VGF: THLGEALAPLSK | CSF | GRN | 17 | -0.509 | 0.037 |
| VGF: THLGEALAPLSK | CSF | C9 | 15 | -0.014 | 0.959 |
| VGF: THLGEALAPLSK | CSF | MAPT | 21 | -0.651 | 0.001 |
| CHGA: RPEDQELESLSAIEAELEK | CSF | non-carrier | 38 | -0.164 | 0.325 |
| CHGA: RPEDQELESLSAIEAELEK | CSF | GRN | 17 | -0.467 | 0.059 |
| CHGA: RPEDQELESLSAIEAELEK | CSF | C9 | 15 | 0.023 | 0.934 |
| CHGA: RPEDQELESLSAIEAELEK | CSF | MAPT | 21 | -0.528 | 0.014 |

### Materials and Methods
#### Broad Targeted LMCS

**Supplemental Table 3: Metabolomics in negative mode parameters**

| Metabolite | Internal Std | Q1 (m/z) | Q3(m/z) | CE (V) |
|---|---|---|---|---|
| 13C3-Citric acid | N/A | 194.2 | 113.0 | -11 |
| 13C4-Fumaric acid | N/A | 119.2 | 74.3 | -13 |
| 13C5-Itaconic acid | N/A | 134.0 | 89.2 | -16 |
| 13C4-Malic acid | N/A | 137.2 | 119.2 | -15 |
| 13C4-Succinic acid | N/A | 121.3 | 76.0 | -12 |
| 13C5-Oxoglutaric acid | N/A | 150.2 | 105.3 | -14 |
| 15N5-ADP | N/A | 431.1 | 79 | -56 |
| 2-Hydroxyglutarate | Glutamic acid-d3 | 145 | 101 | -14 |
| 2-Phosphoglyceric acid | Glutamic acid-d3 | 185.5 | 97.0 | -35 |
| 3-Hydroxybutyric acid | Glutamic acid-d3 | 103.2 | 59.2 | -14 |
| 3-Phosphoglyceric acid | Glutamic acid-d3 | 185.5 | 97.1 | -35 |
| 6-Phosphogluconic acid | Glutamic acid-d3 | 275.0 | 97.0 | -21 |
| Acetoacetic acid | Glutamic acid-d3 | 101 | 101 | -5 |
| Adenosine monophosphate | U-13C10,U-15N5-Adenosine monophosphate | 346 | 79 | -42 |
| Adenosine triphosphate | 15N5-ADP | 506.1 | 158.9 | -84 |
| Adenosine triphosphate-d4 | N/A | 510 | 158.9 | -84 |
| ADP | 15N5-ADP | 426.1 | 79 | -92 |
| Alanine | Arginine-d4C13 | 88 | 88 | -19 |
| Alanine-d4 | N/A | 92 | 92 | -19 |
| Arginine | Arginine-d4C13 | 173 | 131 | -19 |
| Arginine-d4C13 | N/A | 177 | 136 | -19 |
| Asparagine | Glutamic acid-d3 | 131 | 114 | -18 |
| Aspartic acid-d3 | 13C4-Succinic acid | 132 | 88 | -18 |
| Aspartic acid-d3 | N/A | 135 | 91 | -18 |
| cis-Aconitic acid | Glutamic acid-d3 | 173.0 | 129.0 | -12 |
| Dihydroxyacetone phosphate | Glutamic acid-d3 | 169.1 | 79.0 | -38 |
| Erythose 4-phosphate | Glutamic acid-d3 | 199.0 | 97.0 | -21 |
| Fructose 1-phosphate | Glutamic acid-d3 | 259.1 | 79.0 | -74 |
| Fructose 1,6-bisphosphate | Glutamic acid-d3 | 339.0 | 97.0 | -74 |
| Fructose 6-phosphate | Glutamic acid-d3 | 259.0 | 97.1 | -74 |
| Fumaric acid | 13C4-Succinic acid | 115.2 | 71.3 | -74 |
| Glucose 1-phosphate | U-13C6-Glucose 6-phosphate | 259 | 241 | -21 |
| Glucose 6-phosphate | U-13C6-Glucose 6-phosphate | 259.0 | 97.0 | -21 |

| Metabolite | Internal Std | Q1 (m/z) | Q3(m/z) | CE (V) |
|---|---|---|---|---|
| 13C3-Citric acid | N/A | 194.2 | 113.0 | -11 |
| 13C4-Fumaric acid | N/A | 119.2 | 74.3 | -13 |
| 13C5-Itaconic acid | N/A | 134.0 | 89.2 | -16 |
| 13C4-Malic acid | N/A | 137.2 | 119.2 | -15 |
| 13C4-Succinic acid | N/A | 121.3 | 76.0 | -12 |
| 13C5-Oxoglutaric acid | N/A | 150.2 | 105.3 | -14 |
| 15N5-ADP | N/A | 431.1 | 79 | -56 |
| 2-Hydroxyglutarate | Glutamic acid-d3 | 145 | 101 | -14 |
| 2-Phosphoglyceric acid | Glutamic acid-d3 | 185.5 | 97.0 | -35 |
| 3-Hydroxybutyric acid | Glutamic acid-d3 | 103.2 | 59.2 | -14 |
| 3-Phosphoglyceric acid | Glutamic acid-d3 | 185.5 | 97.1 | -35 |
| 6-Phosphogluconic acid | Glutamic acid-d3 | 275.0 | 97.0 | -21 |
| Acetoacetic acid | Glutamic acid-d3 | 101 | 101 | -5 |
| Adenosine monophosphate | U-13C10,U-15N5-Adenosine monophosphate | 346 | 79 | -42 |
| Glutamic acid | Glutamic acid-d3 | 146 | 128 | -14 |
| Glutamic acid-d3 | Glutamic acid-d3 | 149 | 131 | -14 |
| Glutamine | Glutamic acid-d3 | 145 | 127 | -13 |
| Glyceraldehyde 3-phosphate | Glutamic acid-d3 | 169.1 | 97.0 | -15 |
| Itaconic acid | 13C5-Itaconic acid | 129.2 | 85.0 | -10 |
| Lactic acid | Glutamic acid-d3 | 89.2 | 43.2 | -16 |
| Malic acid | 13C4-Succinic acid | 133.2 | 115.2 | -15 |
| NAD | Glutamic acid-d3 | 662.1 | 540.3 | -17 |
| NADH | Glutamic acid-d3 | 664.1 | 408.1 | -15 |
| NADP | Glutamic acid-d3 | 742.1 | 620 | -44 |
| NADPH | Glutamic acid-d3 | 744.1 | 408 | -44 |
| Oxalacetic acid | U-13C6-Glucose 6-phosphate | 131.0 | 131.0 | -13 |
| Oxoglutaric acid | 13C4-Succinic acid | 145.2 | 101.3 | -14 |
| Phosphocreatine | Glutamic acid-d3 | 210 | 79 | -21 |
| Phosphoenolpyruvic acid | Glutamic acid-d3 | 167.0 | 79.0 | -25 |
| Pyruvic acid | U-13C3-Pyruvic acid | 87.2 | 43.2 | -12 |
| Ribose 5-phosphate | Glutamic acid-d3 | 229.0 | 97.0 | -19 |
| Ribulose 5-phosphate | Glutamic acid-d3 | 229.0 | 97.1 | -17 |
| Sedoheptulose 7-phosphate | Glutamic acid-d3 | 289.0 | 97.0 | -27 |
| Serine | Glutamic acid-d3 | 104 | 74 | -16 |
| Succinic acid | 13C4-Succinic acid | 117.2 | 73.2 | -14 |
| Sum of Citric acid and Isocitric acid | Glutamic acid-d3 | 191.2 | 111.0 | -19 |

| Metabolite | Internal Std | Q1 (m/z) | Q3(m/z) | CE (V) |
|---|---|---|---|---|
| 13C3-Citric acid | N/A | 194.2 | 113.0 | -11 |
| 13C4-Fumaric acid | N/A | 119.2 | 74.3 | -13 |
| 13C5-Itaconic acid | N/A | 134.0 | 89.2 | -16 |
| 13C4-Malic acid | N/A | 137.2 | 119.2 | -15 |
| 13C4-Succinic acid | N/A | 121.3 | 76.0 | -12 |
| 13C5-Oxoglutaric acid | N/A | 150.2 | 105.3 | -14 |
| 15N5-ADP | N/A | 431.1 | 79 | -56 |
| 2-Hydroxyglutarate | Glutamic acid-d3 | 145 | 101 | -14 |
| 2-Phosphoglyceric acid | Glutamic acid-d3 | 185.5 | 97.0 | -35 |
| 3-Hydroxybutyric acid | Glutamic acid-d3 | 103.2 | 59.2 | -14 |
| 3-Phosphoglyceric acid | Glutamic acid-d3 | 185.5 | 97.1 | -35 |
| 6-Phosphogluconic acid | Glutamic acid-d3 | 275.0 | 97.0 | -21 |
| Acetoacetic acid | Glutamic acid-d3 | 101 | 101 | -5 |
| Adenosine monophosphate | U-13C10,U-15N5-Adenosine monophosphate | 346 | 79 | -42 |
| U-13C10,U-15N5-Adenosine monophosphate | N/A | 361 | 79 | -42 |
| U-13C3-Pyruvic acid | N/A | 90.2 | 45.2 | -12 |
| U-13C6-Glucose 6-phosphate | N/A | 265.0 | 97.0 | -21 |
| Xylulose 5-phosphate | Glutamic acid-d3 | 229.1 | 97.0 | -21 |

**Supplemental Table 4: Lipidomics in positive mode parameters using TQ 6495C**

| Lipid | Internal Standard | Q1(m/z) | Q3(m/z) | CE (V) |
|---|---|---|---|---|
| 1-O-Palmitoyl-Cer(d18:1/18:0) | Cer(d18:1/16:0(d7)) | 786.8 | 502.5 | 35 |
| 24-Hydroxycholesterol | 24-Hydroxcholesterol-d7 | 385.3 | 367.3 | 30 |
| 24-Hydroxycholesterol(d7) | IS | 392.3 | 367.3 | 30 |
| 3-O-SulfoLacCer(d18:1/18:0) | LacCer(d18:1/17:0) | 970.8 | 548.5 | 61 |
| 4-beta-Hydroxycholesterol | Cholesterol(d7) | 420.3 | 385.3 | 15 |
| 7-keto-Cholesterol | Cholesterol(d7) | 401.3 | 383.3 | 15 |
| CE HETE | CE(18:1(d7)) | 706.6 | 369.2 | 25 |
| CE HODE | CE(18:1(d7)) | 682.6 | 369.2 | 25 |
| CE HpODE | CE(18:1(d7)) | 698.6 | 369.2 | 25 |
| CE oxoHETE | CE(18:1(d7)) | 704.6 | 369.2 | 25 |
| CE oxoODE | CE(18:1(d7)) | 680.6 | 369.2 | 25 |
| CE(16:1) | CE(18:1(d7)) | 640.6 | 369.3 | 26 |
| CE(18:1(d7)) | IS | 675.2 | 369.4 | 26 |
| CE(18:1) | CE(18:1(d7)) | 668.6 | 369.3 | 26 |
| CE(18:2) | CE(18:1(d7)) | 666.6 | 369.3 | 26 |
| CE(20:4) | CE(18:1(d7)) | 690.6 | 369.3 | 26 |
| CE(20:5) | CE(18:1(d7)) | 688.6 | 369.3 | 26 |
| CE(22:6) | CE(18:1(d7)) | 714.6 | 369.3 | 26 |
| Cer(d18:0/16:0) | Cer(d18:1/16:0(d7)) | 540.6 | 284.3 | 40 |
| Cer(d18:0/18:0) | Cer(d18:1/16:0(d7)) | 568.7 | 284.3 | 40 |
| Cer(d18:0/24:0) | Cer(d18:1/16:0(d7)) | 652.9 | 284.3 | 40 |
| Cer(d18:0/24:1) | Cer(d18:1/16:0(d7)) | 650.9 | 284.4 | 40 |
| Cer(d18:1/16:0(d7)) | IS | 545.5 | 271.4 | 40 |
| Cer(d18:1/16:0) | Cer(d18:1/16:0(d7)) | 538.5 | 264.3 | 40 |
| Cer(d18:1/18:0) | Cer(d18:1/16:0(d7)) | 566.6 | 264.3 | 40 |
| Cer(d18:1/24:0) | Cer(d18:1/16:0(d7)) | 650.6 | 264.3 | 40 |
| Cer(d18:1/24:1) | Cer(d18:1/16:0(d7)) | 648.6 | 264.3 | 40 |
| Cholesterol | Cholesterol(d7) | 369.3 | 369.3 | 10 |
| Cholesterol(d7) | IS | 376.2 | 376.2 | 10 |
| Cholesteryl hexoside | CE(18:1(d7)) | 566.6 | 369.3 | 17 |
| Coenzyme Q10 | TG(15:0/18:1(d7)/15:0) | 863.3 | 197.2 | 35 |
| DG(15:0/18:1(d7)) | IS | 605.6 | 346.5 | 30 |
| DG(16:0_18:1) | DG(15:0/18:1(d7)) | 612.4 | 313.3 | 30 |
| DG(16:0_20:4) | DG(15:0/18:1(d7)) | 634.5 | 313.3 | 30 |
| DG(18:0_18:1) | DG(15:0/18:1(d7)) | 640.4 | 341.3 | 30 |
| DG(18:0_20:4) | DG(15:0/18:1(d7)) | 662.5 | 341.3 | 30 |
| DG(18:0_22:6) | DG(15:0/18:1(d7)) | 686.6 | 341.3 | 30 |

| Lipid | Internal Standard | Q1(m/z) | Q3(m/z) | CE (V) |
|---|---|---|---|---|
| 1-O-Palmitoyl-Cer(d18:1/18:0) | Cer(d18:1/16:0(d7)) | 786.8 | 502.5 | 35 |
| 24-Hydroxycholesterol | 24-Hydroxcholesterol-d7 | 385.3 | 367.3 | 30 |
| 24-Hydroxycholesterol(d7) | IS | 392.3 | 367.3 | 30 |
| 3-O-SulfoLacCer(d18:1/18:0) | LacCer(d18:1/17:0) | 970.8 | 548.5 | 61 |
| DG(18:1_20:4) | DG(15:0/18:1(d7)) | 660.5 | 339.3 | 30 |
| DG(18:1/18:1) | DG(15:0/18:1(d7)) | 638.4 | 339.3 | 30 |
| GB3(d18:1/16:0) | GB3(d18:1/18:0(d3)) | 1025 | 520.5 | 40 |
| GB3(d18:1/18:0(d3)) | IS | 1056 | 551.6 | 40 |
| GB3(d18:1/18:0) | GB3(d18:1/18:0(d3)) | 1053 | 548.6 | 40 |
| GB3(d18:1/24:0) | GB3(d18:1/18:0(d3)) | 1137 | 632.6 | 40 |
| GB3(d18:1/24:1) | GB3(d18:1/18:0(d3)) | 1135 | 630.6 | 40 |
| GlcCer(d18:1(d5)/18:0) | IS | 733.6 | 269.3 | 45 |
| GlcCer(d18:1/16:0(d3)) | IS | 703.7 | 264.3 | 51 |
| Glucosylsphingosine(d5) | IS | 467.2 | 269.3 | 16 |
| HexCer(d18:1/12:0) | GlcCer(d18:1(d5)/18:0) | 644.5 | 264.3 | 40 |
| HexCer(d18:1/16:0) | GlcCer(d18:1(d5)/18:0) | 700.6 | 264.3 | 40 |
| HexCer(d18:1/18:0) | GlcCer(d18:1(d5)/18:0) | 728.6 | 264.3 | 40 |
| HexCer(d18:1/22:0) | GlcCer(d18:1(d5)/18:0) | 784.7 | 264.4 | 40 |
| HexCer(d18:1/24:0) | GlcCer(d18:1(d5)/18:0) | 812.7 | 264.3 | 40 |
| HexCer(d18:1/24:1) | GlcCer(d18:1(d5)/18:0) | 810.7 | 264.3 | 40 |
| Hexosylsphingosine | Glucosylsphingosine(d5) | 462.3 | 264.2 | 16 |
| LacCer(d18:1/16:0) | LacCer(d18:1/17:0) | 862.6 | 264.3 | 40 |
| LacCer(d18:1/17:0) | IS | 876.6 | 264.3 | 40 |
| LacCer(d18:1/18:0) | LacCer(d18:1/17:0) | 890.7 | 264.3 | 40 |
| LacCer(d18:1/24:0) | LacCer(d18:1/17:0) | 974.8 | 264.3 | 40 |
| LacCer(d18:1/24:1) | LacCer(d18:1/17:0) | 972.7 | 264.3 | 40 |
| Lactosylsphingosine | Glucosylsphingosine(d5) | 624.4 | 264.3 | 16 |
| LPC(16:0) | LPC(18:1(d7)) | 496.3 | 184.1 | 40 |
| LPC(16:1) | LPC(18:1(d7)) | 494.5 | 184.1 | 40 |
| LPC(18:0) | LPC(18:1(d7)) | 524.3 | 184.1 | 40 |
| LPC(18:1(d7)) | IS | 529.3 | 184.1 | 40 |
| LPC(18:1) | LPC(18:1(d7)) | 522.3 | 184.1 | 40 |
| LPC(20:4) | LPC(18:1(d7)) | 544.3 | 184.1 | 40 |
| LPC(22:6) | LPC(18:1(d7)) | 568.3 | 184.1 | 40 |
| LPC(24:0) | LPC(18:1(d7)) | 608.5 | 184.1 | 40 |
| LPC(24:1) | LPC(18:1(d7)) | 606.5 | 184.1 | 40 |
| LPC(26:0) | LPC(18:1(d7)) | 636.5 | 104.1 | 40 |
| LPC(26:1) | LPC(18:1(d7)) | 634.5 | 104.1 | 40 |
| lyso-GB3 | lyso-GB3-d7 | 786.6 | 264.3 | 46 |

| Lipid | Internal Standard | Q1(m/z) | Q3(m/z) | CE (V) |
|---|---|---|---|---|
| 1-O-Palmitoyl-Cer(d18:1/18:0) | Cer(d18:1/16:0(d7)) | 786.8 | 502.5 | 35 |
| 24-Hydroxycholesterol | 24-Hydroxcholesterol-d7 | 385.3 | 367.3 | 30 |
| 24-Hydroxycholesterol(d7) | IS | 392.3 | 367.3 | 30 |
| 3-O-SulfoLacCer(d18:1/18:0) | LacCer(d18:1/17:0) | 970.8 | 548.5 | 61 |
| lyso-GB3-d7 | IS | 793.5 | 271.3 | 46 |
| lyso-GB4 | GB3(d18:1/18:0(d3)) | 990.6 | 264.3 | 52 |
| MG(16:0) | MG(18:1(d7)) | 348.3 | 239.3 | 22 |
| MG(16:1) | MG(18:1(d7)) | 346.3 | 237.3 | 22 |
| MG(18:0) | MG(18:1(d7)) | 376.3 | 267.3 | 22 |
| MG(18:1(d7)) | IS | 381.3 | 272.5 | 22 |
| MG(18:1) | MG(18:1(d7)) | 374.3 | 265.3 | 22 |
| MG(20:4) | MG(18:1(d7)) | 396.3 | 287.3 | 22 |
| N-Oleoylethanolamine | LPC(18:1(d7)) | 326.3 | 62.1 | 23 |
| N-Palmitoyl-O-phosphocholineserine | LPC(18:1(d7)) | 509.5 | 184.1 | 40 |
| Palmitoylethanolamine | LPC(18:1(d7)) | 300.3 | 62.1 | 23 |
| PC(15:0/18:1(d7)) | IS | 754.6 | 184.1 | 40 |
| PC(16:0/5:0(CHO)) | PC(15:0/18:1(d7)) | 594.5 | 184.1 | 40 |
| PC(16:0/9:0(CHO)) | PC(15:0/18:1(d7)) | 650.4 | 184.1 | 40 |
| PC(16:0/9:0(COOH)) | PC(15:0/18:1(d7)) | 666.4 | 184.1 | 40 |
| PC(18:0/20:4(OH[S])) | PC(15:0/18:1(d7)) | 826.6 | 184.1 | 40 |
| PC(18:0/20:4(OOH[S])) | PC(15:0/18:1(d7)) | 842.6 | 184.1 | 40 |
| PC(34:1) | PC(15:0/18:1(d7)) | 760.6 | 184.1 | 40 |
| PC(36:1) | PC(15:0/18:1(d7)) | 788.6 | 184.1 | 40 |
| PC(36:2) | PC(15:0/18:1(d7)) | 786.6 | 184.1 | 40 |
| PC(36:4) | PC(15:0/18:1(d7)) | 782.6 | 184.1 | 40 |
| PC(38:4) | PC(15:0/18:1(d7)) | 810.6 | 184.1 | 40 |
| PC(38:6) | PC(15:0/18:1(d7)) | 806.6 | 184.1 | 40 |
| PC(40:5) | PC(15:0/18:1(d7)) | 836.6 | 184.1 | 40 |
| PC(40:6) | PC(15:0/18:1(d7)) | 834.6 | 184.1 | 40 |
| PC(O-16:0/0:0) | LPC(18:1(d7)) | 482.3 | 184.1 | 40 |
| PC(O-16:0/2:0) | LPC(18:1(d7)) | 524.3 | 184.2 | 40 |
| PC(O-18:0/2:0) | LPC(18:1(d7)) | 552.5 | 184.1 | 40 |
| PE(15:0/18:1(d7)) | IS | 711.6 | 570.5 | 40 |
| PE(18:0/20:4(OH[S])) | PE(15:0/18:1(d7)) | 784.5 | 643.4 | 40 |
| PE(18:0/20:4(OOH[S])) | PE(15:0/18:1(d7)) | 800.5 | 659.4 | 40 |
| PE(34:1) | PE(15:0/18:1(d7)) | 718.6 | 577.6 | 40 |
| PE(36:1) | PE(15:0/18:1(d7)) | 746.6 | 605.5 | 40 |
| PE(36:2) | PE(15:0/18:1(d7)) | 744.6 | 603.5 | 40 |

| Lipid | Internal Standard | Q1(m/z) | Q3(m/z) | CE (V) |
|---|---|---|---|---|
| 1-O-Palmitoyl-Cer(d18:1/18:0) | Cer(d18:1/16:0(d7)) | 786.8 | 502.5 | 35 |
| 24-Hydroxycholesterol | 24-Hydroxcholesterol-d7 | 385.3 | 367.3 | 30 |
| 24-Hydroxycholesterol(d7) | IS | 392.3 | 367.3 | 30 |
| 3-O-SulfoLacCer(d18:1/18:0) | LacCer(d18:1/17:0) | 970.8 | 548.5 | 61 |
| PE(36:4) | PE(15:0/18:1(d7)) | 740.6 | 599.5 | 40 |
| PE(38:4) | PE(15:0/18:1(d7)) | 768.6 | 627.5 | 40 |
| PE(38:5) | PE(15:0/18:1(d7)) | 766.6 | 625.5 | 40 |
| PE(38:6) | PE(15:0/18:1(d7)) | 764.6 | 623.5 | 40 |
| PE(40:4) | PE(15:0/18:1(d7)) | 796.6 | 655.5 | 40 |
| PE(40:5) | PE(15:0/18:1(d7)) | 794.6 | 635.5 | 40 |
| PE(40:6) | PE(15:0/18:1(d7)) | 792.6 | 651.5 | 40 |
| PE(40:7) | PE(15:0/18:1(d7)) | 790.6 | 649.5 | 40 |
| Sitosteryl hexoside | CE(18:1(d7)) | 594.6 | 397.4 | 17 |
| SM(d18:1(d9)/18:1) | IS | 738.7 | 184.1 | 40 |
| SM(d18:1/16:0) | SM(d18:1(d9)/18:1) | 703.6 | 184.1 | 40 |
| SM(d18:1/18:0) | SM(d18:1(d9)/18:1) | 731.6 | 184.1 | 40 |
| SM(d18:1/24:0) | SM(d18:1(d9)/18:1) | 815.7 | 184.1 | 40 |
| SM(d18:1/24:1) | SM(d18:1(d9)/18:1) | 813.7 | 184.1 | 40 |
| Sphinganine | Sphingosine(d17:1) | 302.2 | 284.3 | 20 |
| Sphinganine 1-phosphate | Sphingosine 1-phosphate-d7 | 382.3 | 284.3 | 18 |
| Sphingosine | Sphingosine(d17:1) | 300.2 | 264.3 | 20 |
| Sphingosine 1-phosphate | Sphingosine 1-phosphate-d7 | 380.3 | 264.3 | 25 |
| Sphingosine 1-phosphate-d7 | IS | 387.3 | 271.3 | 25 |
| Sphingosine 1-phosphocholine | LPC(18:1(d7)) | 465.5 | 184.1 | 40 |
| Sphingosine(d17:1) | IS | 286.2 | 250.3 | 20 |
| TG(15:0/18:1(d7)/15:0) | IS | 829.8 | 570.8 | 40 |
| TG(18:0_36:2) | TG(15:0/18:1(d7)/15:0) | 904.7 | 603.4 | 40 |
| TG(18:1_34:2) | TG(15:0/18:1(d7)/15:0) | 874.7 | 575.4 | 40 |
| TG(18:1_34:3) | TG(15:0/18:1(d7)/15:0) | 872.7 | 573.4 | 40 |
| TG(20:4_32:1) | TG(15:0/18:1(d7)/15:0) | 870.6 | 549.3 | 40 |
| TG(20:4_34:2) | TG(15:0/18:1(d7)/15:0) | 896.6 | 575.3 | 40 |
| TG(20:4_34:3) | TG(15:0/18:1(d7)/15:0) | 894.6 | 573.3 | 40 |
| TG(20:4_36:0) | TG(15:0/18:1(d7)/15:0) | 928.8 | 607.5 | 40 |
| TG(20:4_36:2) | TG(15:0/18:1(d7)/15:0) | 924.7 | 603.4 | 40 |
| TG(20:4_36:3) | TG(15:0/18:1(d7)/15:0) | 922.7 | 601.4 | 40 |

| **Lipid** | **Internal Standard** | **Q1(m/z)** | **Q3(m/z)** | **CE (V)** |
|---|---|---|---|---|
| 1-O-Palmitoyl-Cer(d18:1/18:0) | Cer(d18:1/16:0(d7)) | 786.8 | 502.5 | 35 |
| 24-Hydroxycholesterol | 24-Hydroxcholesterol-d7 | 385.3 | 367.3 | 30 |
| 24-Hydroxycholesterol(d7) | IS | 392.3 | 367.3 | 30 |
| 3-O-SulfoLacCer(d18:1/18:0) | LacCer(d18:1/17:0) | 970.8 | 548.5 | 61 |
| TG(22:6_36:2) | TG(15:0/18:1(d7)/15:0) | 948.7 | 603.4 | 40 |
| TG(22:6_38:1) | TG(15:0/18:1(d7)/15:0) | 978.7 | 633.4 | 40 |
| TG(22:6_38:2) | TG(15:0/18:1(d7)/15:0) | 976.7 | 631.4 | 40 |

#### Supplemental Table 5: Proteomics MRM MS Parameters

| Protein | Peptide | SIL | Precursor Ion | Product Ion | Collision Energy |
|---|---|---|---|---|---|
| sp\|P47972\|NPTX2_HUMAN | LESLEHQLR.light | FALSE | 375.540133 | 416.261593 | 11.7 |
| sp\|P47972\|NPTX2_HUMAN | LESLEHQLR.light | FALSE | 375.540133 | 553.320505 | 11.7 |
| sp\|P47972\|NPTX2_HUMAN | LESLEHQLR.light | FALSE | 375.540133 | 682.363098 | 11.7 |
| sp\|P47972\|NPTX2_HUMAN | LESLEHQLR.heavy | TRUE | 377.879188 | 423.278757 | 11.7 |
| sp\|P47972\|NPTX2_HUMAN | LESLEHQLR.heavy | TRUE | 377.879188 | 560.337669 | 11.7 |
| sp\|P47972\|NPTX2_HUMAN | LESLEHQLR.heavy | TRUE | 377.879188 | 689.380262 | 11.7 |
| sp\|O15240\|VGF_HUMAN | THLGEALAPLSK.light | FALSE | 412.903072 | 444.28166 | 4.1 |
| sp\|O15240\|VGF_HUMAN | THLGEALAPLSK.light | FALSE | 412.903072 | 515.318774 | 10.1 |
| sp\|O15240\|VGF_HUMAN | THLGEALAPLSK.light | FALSE | 412.903072 | 609.299101 | 7.1 |
| sp\|O15240\|VGF_HUMAN | THLGEALAPLSK.heavy | TRUE | 415.242126 | 451.298824 | 4.1 |
| sp\|O15240\|VGF_HUMAN | THLGEALAPLSK.heavy | TRUE | 415.242126 | 522.335938 | 10.1 |
| sp\|O15240\|VGF_HUMAN | THLGEALAPLSK.heavy | TRUE | 415.242126 | 609.299101 | 7.1 |
| sp\|P47972\|NPTX2_HUMAN | TESTLNALLQR.light | FALSE | 415.898098 | 529.345657 | 7.2 |
| sp\|P47972\|NPTX2_HUMAN | TESTLNALLQR.light | FALSE | 415.898098 | 600.382771 | 10.2 |
| sp\|P47972\|NPTX2_HUMAN | TESTLNALLQR.light | FALSE | 415.898098 | 717.34136 | 7.2 |
| sp\|Q92932\|PTPR2_HUMAN | THTAQDRPPAEGDDR.light | FALSE | 417.194845 | 462.194302 | 7.2 |
| sp\|Q92932\|PTPR2_HUMAN | THTAQDRPPAEGDDR.light | FALSE | 417.194845 | 591.236895 | 13.2 |
| sp\|Q92932\|PTPR2_HUMAN | THTAQDRPPAEGDDR.light | FALSE | 417.194845 | 662.274009 | 13.2 |
| sp\|P47972\|NPTX2_HUMAN | TESTLNALLQR.heavy | TRUE | 418.237152 | 536.362821 | 7.2 |
| sp\|P47972\|NPTX2_HUMAN | TESTLNALLQR.heavy | TRUE | 418.237152 | 607.399935 | 10.2 |
| sp\|P47972\|NPTX2_HUMAN | TESTLNALLQR.heavy | TRUE | 418.237152 | 717.34136 | 7.2 |
| sp\|Q92932\|PTPR2_HUMAN | THTAQDRPPAEGDDR.heavy | TRUE | 418.698297 | 462.194302 | 7.2 |
| sp\|Q92932\|PTPR2_HUMAN | THTAQDRPPAEGDDR.heavy | TRUE | 418.698297 | 591.236895 | 13.2 |
| sp\|Q92932\|PTPR2_HUMAN | THTAQDRPPAEGDDR.heavy | TRUE | 418.698297 | 662.274009 | 13.2 |
| sp\|P05067\|A4_HUMAN | THPHFVIPYR.light | FALSE | 422.896251 | 473.225542 | 13.4 |
| sp\|P05067\|A4_HUMAN | THPHFVIPYR.light | FALSE | 422.896251 | 548.319108 | 13.4 |
| sp\|P05067\|A4_HUMAN | THPHFVIPYR.light | FALSE | 422.896251 | 719.36237 | 7.4 |
| sp\|P05067\|A4_HUMAN | THPHFVIPYR.heavy | TRUE | 424.900854 | 473.225542 | 13.4 |
| sp\|P05067\|A4_HUMAN | THPHFVIPYR.heavy | TRUE | 424.900854 | 554.332917 | 13.4 |
| sp\|P05067\|A4_HUMAN | THPHFVIPYR.heavy | TRUE | 424.900854 | 719.36237 | 7.4 |
| sp\|P68871\|HBB_HUMAN | LLGNVLVC[+57.021464]VLAHHFGK.light | FALSE | 445.003996 | 488.261593 | 8.2 |
| sp\|P68871\|HBB_HUMAN | LLGNVLVC[+57.021464]VLAHHFGK.light | FALSE | 445.003996 | 497.308209 | 14.2 |
| sp\|P68871\|HBB_HUMAN | LLGNVLVC[+57.021464]VLAHHFGK.light | FALSE | 445.003996 | 610.392273 | 14.2 |
| sp\|P68871\|HBB_HUMAN | LLGNVLVC[+57.021464]VLAHHFGK.heavy | TRUE | 447.510803 | 497.308209 | 8.2 |
| sp\|P68871\|HBB_HUMAN | LLGNVLVC[+57.021464]VLAHHFGK.heavy | TRUE | 447.510803 | 498.288821 | 14.2 |
| sp\|P68871\|HBB_HUMAN | LLGNVLVC[+57.021464]VLAHHFGK.heavy | TRUE | 447.510803 | 610.392273 | 14.2 |

| Protein | Peptide | SIL | Precursor Ion | Product Ion | Collision Energy |
|---|---|---|---|---|---|
| sp\|P01876\|IGHA1_HUMAN | TPLTATLSK.light | FALSE | 466.276575 | 519.313689 | 24.5 |
| sp\|P01876\|IGHA1_HUMAN | TPLTATLSK.light | FALSE | 466.276575 | 620.361367 | 15.5 |
| sp\|P01876\|IGHA1_HUMAN | TPLTATLSK.light | FALSE | 466.276575 | 733.445431 | 12.5 |
| sp\|P01876\|IGHA1_HUMAN | TPLTATLSK.heavy | TRUE | 469.785157 | 526.330853 | 24.5 |
| sp\|P01876\|IGHA1_HUMAN | TPLTATLSK.heavy | TRUE | 469.785157 | 627.378531 | 15.5 |
| sp\|P01876\|IGHA1_HUMAN | TPLTATLSK.heavy | TRUE | 469.785157 | 740.462595 | 12.5 |
| sp\|P02649\|APOE_HUMAN | LGPLVEQGR.light | FALSE | 484.779816 | 588.31 | 19 |
| sp\|P02649\|APOE_HUMAN | LGPLVEQGR.light | FALSE | 484.779816 | 701.394064 | 25 |
| sp\|P02649\|APOE_HUMAN | LGPLVEQGR.light | FALSE | 484.779816 | 798.446828 | 19 |
| sp\|P02649\|APOE_HUMAN | LGPLVEQGR.heavy | TRUE | 487.78672 | 594.323809 | 19 |
| sp\|P02649\|APOE_HUMAN | LGPLVEQGR.heavy | TRUE | 487.78672 | 707.407873 | 25 |
| sp\|P02649\|APOE_HUMAN | LGPLVEQGR.heavy | TRUE | 487.78672 | 804.460637 | 19 |
| sp\|P10645\|CMGA_HUMAN | EAVEEPSSK.light | FALSE | 488.235104 | 418.229624 | 19.1 |
| sp\|P10645\|CMGA_HUMAN | EAVEEPSSK.light | FALSE | 488.235104 | 547.272218 | 13.1 |
| sp\|P10645\|CMGA_HUMAN | EAVEEPSSK.light | FALSE | 488.235104 | 676.314811 | 10.1 |
| sp\|P10645\|CMGA_HUMAN | EAVEEPSSK.heavy | TRUE | 491.242008 | 424.243433 | 19.1 |
| sp\|P10645\|CMGA_HUMAN | EAVEEPSSK.heavy | TRUE | 491.242008 | 553.286027 | 13.1 |
| sp\|P10645\|CMGA_HUMAN | EAVEEPSSK.heavy | TRUE | 491.242008 | 682.32862 | 10.1 |
| sp\|P28799\|GRN_HUMAN | VHC[+57.021464]C[+57.021464]PHGAFC[+57.021464]DLVHTR.light | FALSE | 492.221182 | 512.293956 | 12.9 |
| sp\|P28799\|GRN_HUMAN | VHC[+57.021464]C[+57.021464]PHGAFC[+57.021464]DLVHTR.light | FALSE | 492.221182 | 656.814396 | 21.9 |
| sp\|P28799\|GRN_HUMAN | VHC[+57.021464]C[+57.021464]PHGAFC[+57.021464]DLVHTR.light | FALSE | 492.221182 | 705.340777 | 15.9 |
| sp\|Q9NZC2\|TREM2_HUMAN | VLVEVLADPLDHR.light | FALSE | 492.612069 | 582.804019 | 6.9 |
| sp\|Q9NZC2\|TREM2_HUMAN | VLVEVLADPLDHR.light | FALSE | 492.612069 | 632.338226 | 6.9 |
| sp\|Q9NZC2\|TREM2_HUMAN | VLVEVLADPLDHR.light | FALSE | 492.612069 | 637.341635 | 21.9 |
| sp\|P00915\|CAH1_HUMAN | GGPFSDSYR.light | FALSE | 493.222331 | 627.27328 | 16.3 |
| sp\|P00915\|CAH1_HUMAN | GGPFSDSYR.light | FALSE | 493.222331 | 774.341694 | 16.3 |
| sp\|P00915\|CAH1_HUMAN | GGPFSDSYR.light | FALSE | 493.222331 | 871.394458 | 16.3 |
| sp\|P28799\|GRN_HUMAN | VHC[+57.021464]C[+57.021464]PHGAFC[+57.021464]DLVHTR.heavy | TRUE | 493.724635 | 518.307765 | 12.9 |
| sp\|P28799\|GRN_HUMAN | VHC[+57.021464]C[+57.021464]PHGAFC[+57.021464]DLVHTR.heavy | TRUE | 493.724635 | 659.8213 | 21.9 |
| sp\|P28799\|GRN_HUMAN | VHC[+57.021464]C[+57.021464]PHGAFC[+57.021464]DLVHTR.heavy | TRUE | 493.724635 | 708.347682 | 15.9 |
| sp\|Q9NZC2\|TREM2_HUMAN | VLVEVLADPLDHR.heavy | TRUE | 494.951123 | 586.312601 | 6.9 |
| sp\|Q9NZC2\|TREM2_HUMAN | VLVEVLADPLDHR.heavy | TRUE | 494.951123 | 635.846808 | 6.9 |
| sp\|Q9NZC2\|TREM2_HUMAN | VLVEVLADPLDHR.heavy | TRUE | 494.951123 | 644.358799 | 21.9 |
| sp\|O95502\|NPTXR_HUMAN | LVEAFGGATK.light | FALSE | 496.774199 | 580.308937 | 13.4 |
| sp\|O95502\|NPTXR_HUMAN | LVEAFGGATK.light | FALSE | 496.774199 | 651.346051 | 16.4 |
| sp\|O95502\|NPTXR_HUMAN | LVEAFGGATK.light | FALSE | 496.774199 | 780.388644 | 13.4 |
| sp\|P00915\|CAH1_HUMAN | GGPFSDSYR.heavy | TRUE | 498.235945 | 627.27328 | 16.3 |

| Protein | Peptide | SIL | Precursor Ion | Product Ion | Collision Energy |
|---|---|---|---|---|---|
| sp\|P00915\|CAH1_HUMAN | GGPFSDSYR.heavy | TRUE | 498.235945 | 784.368922 | 16.3 |
| sp\|P00915\|CAH1_HUMAN | GGPFSDSYR.heavy | TRUE | 498.235945 | 881.421686 | 16.3 |
| sp\|O95502\|NPTXR_HUMAN | LVEAFGGATK.heavy | TRUE | 501.787813 | 590.336165 | 13.4 |
| sp\|O95502\|NPTXR_HUMAN | LVEAFGGATK.heavy | TRUE | 501.787813 | 661.373279 | 16.4 |
| sp\|O95502\|NPTXR_HUMAN | LVEAFGGATK.heavy | TRUE | 501.787813 | 790.415872 | 13.4 |
| sp\|A6NLU5\|VTM2B_HUMAN | ELLHELALSVPGAR.light | FALSE | 502.287585 | 400.230293 | 7.3 |
| sp\|A6NLU5\|VTM2B_HUMAN | ELLHELALSVPGAR.light | FALSE | 502.287585 | 586.330736 | 13.3 |
| sp\|A6NLU5\|VTM2B_HUMAN | ELLHELALSVPGAR.light | FALSE | 502.287585 | 806.44068 | 13.3 |
| tr\|Q8TCZ8\|Q8TCZ8_HUMAN | LGADMEDVR.light | FALSE | 503.237124 | 518.256902 | 10.6 |
| tr\|Q8TCZ8\|Q8TCZ8_HUMAN | LGADMEDVR.light | FALSE | 503.237124 | 649.297387 | 16.6 |
| tr\|Q8TCZ8\|Q8TCZ8_HUMAN | LGADMEDVR.light | FALSE | 503.237124 | 764.32433 | 16.6 |
| sp\|A6NLU5\|VTM2B_HUMAN | ELLHELALSVPGAR.heavy | TRUE | 504.292188 | 400.230293 | 7.3 |
| sp\|A6NLU5\|VTM2B_HUMAN | ELLHELALSVPGAR.heavy | TRUE | 504.292188 | 592.344545 | 13.3 |
| sp\|A6NLU5\|VTM2B_HUMAN | ELLHELALSVPGAR.heavy | TRUE | 504.292188 | 806.44068 | 13.3 |
| tr\|Q8TCZ8\|Q8TCZ8_HUMAN | LGADMEDVR.heavy | TRUE | 506.244028 | 524.270711 | 10.6 |
| tr\|Q8TCZ8\|Q8TCZ8_HUMAN | LGADMEDVR.heavy | TRUE | 506.244028 | 655.311196 | 16.6 |
| tr\|Q8TCZ8\|Q8TCZ8_HUMAN | LGADMEDVR.heavy | TRUE | 506.244028 | 770.338139 | 16.6 |
| sp\|P07196\|NFL_HUMAN | ALYEQEIR.light | FALSE | 511.269281 | 545.304186 | 16.8 |
| sp\|P07196\|NFL_HUMAN | ALYEQEIR.light | FALSE | 511.269281 | 674.34678 | 13.8 |
| sp\|P07196\|NFL_HUMAN | ALYEQEIR.light | FALSE | 511.269281 | 837.410108 | 10.8 |
| sp\|Q15818\|NPTX1_HUMAN | FQLTFPLR.light | FALSE | 511.295102 | 532.324194 | 25.9 |
| sp\|Q15818\|NPTX1_HUMAN | FQLTFPLR.light | FALSE | 511.295102 | 633.371872 | 16.9 |
| sp\|Q15818\|NPTX1_HUMAN | FQLTFPLR.light | FALSE | 511.295102 | 746.455936 | 10.9 |
| sp\|A6NLU5\|VTM2B_HUMAN | VSDYSDDDTQEHK.light | FALSE | 513.546229 | 541.272886 | 16.7 |
| sp\|A6NLU5\|VTM2B_HUMAN | VSDYSDDDTQEHK.light | FALSE | 513.546229 | 642.320565 | 16.7 |
| sp\|A6NLU5\|VTM2B_HUMAN | VSDYSDDDTQEHK.light | FALSE | 513.546229 | 757.347508 | 13.7 |
| sp\|P07196\|NFL_HUMAN | ALYEQEIR.heavy | TRUE | 514.777863 | 545.304186 | 16.8 |
| sp\|P07196\|NFL_HUMAN | ALYEQEIR.heavy | TRUE | 514.777863 | 674.34678 | 13.8 |
| sp\|P07196\|NFL_HUMAN | ALYEQEIR.heavy | TRUE | 514.777863 | 837.410108 | 10.8 |
| sp\|Q15818\|NPTX1_HUMAN | FQLTFPLR.heavy | TRUE | 514.803684 | 539.341358 | 25.9 |
| sp\|Q15818\|NPTX1_HUMAN | FQLTFPLR.heavy | TRUE | 514.803684 | 640.389036 | 16.9 |
| sp\|Q15818\|NPTX1_HUMAN | FQLTFPLR.heavy | TRUE | 514.803684 | 753.4731 | 10.9 |
| sp\|A6NLU5\|VTM2B_HUMAN | VSDYSDDDTQEHK.heavy | TRUE | 516.888639 | 541.272886 | 16.7 |
| sp\|A6NLU5\|VTM2B_HUMAN | VSDYSDDDTQEHK.heavy | TRUE | 516.888639 | 642.320565 | 16.7 |
| sp\|A6NLU5\|VTM2B_HUMAN | VSDYSDDDTQEHK.heavy | TRUE | 516.888639 | 757.347508 | 13.7 |
| sp\|P63104\|1433Z_HUMAN | SVTEQGAELSNEER.light | FALSE | 516.909391 | 547.247065 | 10.8 |

| Protein | Peptide | SIL | Precursor Ion | Product Ion | Collision Energy |
|---|---|---|---|---|---|
| sp\|P63104\|1433Z_HUMAN | SVTEQGAELSNEER.light | FALSE | 516.909391 | 634.279094 | 10.8 |
| sp\|P63104\|1433Z_HUMAN | SVTEQGAELSNEER.light | FALSE | 516.909391 | 747.363158 | 16.8 |
| sp\|P04156\|PRIO_HUMAN | VVEQMC[+57.021464]ITQYER.light | FALSE | 519.246292 | 587.285759 | 10.9 |
| sp\|P04156\|PRIO_HUMAN | VVEQMC[+57.021464]ITQYER.light | FALSE | 519.246292 | 595.283451 | 10.9 |
| sp\|P04156\|PRIO_HUMAN | VVEQMC[+57.021464]ITQYER.light | FALSE | 519.246292 | 696.33113 | 22.9 |
| sp\|P63104\|1433Z_HUMAN | SVTEQGAELSNEER.heavy | TRUE | 519.248445 | 547.247065 | 10.8 |
| sp\|P63104\|1433Z_HUMAN | SVTEQGAELSNEER.heavy | TRUE | 519.248445 | 634.279094 | 10.8 |
| sp\|P63104\|1433Z_HUMAN | SVTEQGAELSNEER.heavy | TRUE | 519.248445 | 754.380322 | 16.8 |
| sp\|P04156\|PRIO_HUMAN | VVEQMC[+57.021464]ITQYER.heavy | TRUE | 521.250895 | 593.299568 | 10.9 |
| sp\|P04156\|PRIO_HUMAN | VVEQMC[+57.021464]ITQYER.heavy | TRUE | 521.250895 | 595.283451 | 10.9 |
| sp\|P04156\|PRIO_HUMAN | VVEQMC[+57.021464]ITQYER.heavy | TRUE | 521.250895 | 696.33113 | 22.9 |
| sp\|P09936\|UCHL1_HUMAN | LGFEDGSVLK.light | FALSE | 532.784764 | 618.345717 | 14.5 |
| sp\|P09936\|UCHL1_HUMAN | LGFEDGSVLK.light | FALSE | 532.784764 | 747.38831 | 20.5 |
| sp\|P09936\|UCHL1_HUMAN | LGFEDGSVLK.light | FALSE | 532.784764 | 894.456724 | 14.5 |
| sp\|P09936\|UCHL1_HUMAN | LGFEDGSVLK.heavy | TRUE | 535.791668 | 624.359526 | 14.5 |
| sp\|P09936\|UCHL1_HUMAN | LGFEDGSVLK.heavy | TRUE | 535.791668 | 753.402119 | 20.5 |
| sp\|P09936\|UCHL1_HUMAN | LGFEDGSVLK.heavy | TRUE | 535.791668 | 900.470533 | 14.5 |
| sp\|P02649\|APOE_HUMAN | VQAAVGTSAAPVPSDNH.light | FALSE | 540.937392 | 569.231415 | 14.7 |
| sp\|P02649\|APOE_HUMAN | VQAAVGTSAAPVPSDNH.light | FALSE | 540.937392 | 765.352593 | 8.7 |
| sp\|P02649\|APOE_HUMAN | VQAAVGTSAAPVPSDNH.light | FALSE | 540.937392 | 785.415194 | 11.7 |
| sp\|P07360\|CO8G_HUMAN | SLPVSDSVLSGFEQR.light | FALSE | 540.945776 | 579.288536 | 5.7 |
| sp\|P07360\|CO8G_HUMAN | SLPVSDSVLSGFEQR.light | FALSE | 540.945776 | 636.31 | 14.7 |
| sp\|P07360\|CO8G_HUMAN | SLPVSDSVLSGFEQR.light | FALSE | 540.945776 | 723.342029 | 23.7 |
| sp\|P02649\|APOE_HUMAN | VQAAVGTSAAPVPSDNH.heavy | TRUE | 542.941995 | 569.231415 | 14.7 |
| sp\|P02649\|APOE_HUMAN | VQAAVGTSAAPVPSDNH.heavy | TRUE | 542.941995 | 771.366402 | 8.7 |
| sp\|P02649\|APOE_HUMAN | VQAAVGTSAAPVPSDNH.heavy | TRUE | 542.941995 | 785.415194 | 11.7 |
| sp\|P07360\|CO8G_HUMAN | SLPVSDSVLSGFEQR.heavy | TRUE | 544.288186 | 589.315764 | 5.7 |
| sp\|P07360\|CO8G_HUMAN | SLPVSDSVLSGFEQR.heavy | TRUE | 544.288186 | 646.337228 | 14.7 |
| sp\|P07360\|CO8G_HUMAN | SLPVSDSVLSGFEQR.heavy | TRUE | 544.288186 | 733.369257 | 23.7 |
| sp\|P36222\|CH3L1_HUMAN | LVMGIPTFGR.light | FALSE | 545.807519 | 514.305766 | 8.9 |
| sp\|P36222\|CH3L1_HUMAN | LVMGIPTFGR.light | FALSE | 545.807519 | 577.309272 | 17.9 |
| sp\|P36222\|CH3L1_HUMAN | LVMGIPTFGR.light | FALSE | 545.807519 | 747.4148 | 14.9 |
| sp\|P61981\|1433G_HUMAN | NVTELNEPLSNEER.light | FALSE | 548.600557 | 634.279094 | 14.9 |
| sp\|P61981\|1433G_HUMAN | NVTELNEPLSNEER.light | FALSE | 548.600557 | 800.378474 | 11.9 |
| sp\|P61981\|1433G_HUMAN | NVTELNEPLSNEER.light | FALSE | 548.600557 | 844.415922 | 14.9 |
| sp\|P01034\|CYTC_HUMAN | LVGGPMDASVEEEGVR.light | FALSE | 548.934771 | 589.294016 | 18 |
| sp\|P01034\|CYTC_HUMAN | LVGGPMDASVEEEGVR.light | FALSE | 548.934771 | 670.322873 | 9 |

| Protein | Peptide | SIL | Precursor Ion | Product Ion | Collision Energy |
|---|---|---|---|---|---|
| sp\|P01034\|CYTC_HUMAN | LVGGPMDASVEEEGVR.light | FALSE | 548.934771 | 718.336609 | 6 |
| sp\|P36222\|CH3L1_HUMAN | LVMGIPTFGR.heavy | TRUE | 550.821133 | 514.305766 | 8.9 |
| sp\|P36222\|CH3L1_HUMAN | LVMGIPTFGR.heavy | TRUE | 550.821133 | 587.3365 | 17.9 |
| sp\|P36222\|CH3L1_HUMAN | LVMGIPTFGR.heavy | TRUE | 550.821133 | 757.442028 | 14.9 |
| sp\|P01034\|CYTC_HUMAN | LVGGPMDASVEEEGVR.heavy | TRUE | 550.939374 | 589.294016 | 18 |
| sp\|P01034\|CYTC_HUMAN | LVGGPMDASVEEEGVR.heavy | TRUE | 550.939374 | 670.322873 | 9 |
| sp\|P01034\|CYTC_HUMAN | LVGGPMDASVEEEGVR.heavy | TRUE | 550.939374 | 718.336609 | 6 |
| sp\|P61981\|1433G_HUMAN | NVTELNEPLSNEER.heavy | TRUE | 550.939612 | 634.279094 | 14.9 |
| sp\|P61981\|1433G_HUMAN | NVTELNEPLSNEER.heavy | TRUE | 550.939612 | 800.378474 | 11.9 |
| sp\|P61981\|1433G_HUMAN | NVTELNEPLSNEER.heavy | TRUE | 550.939612 | 851.433086 | 14.9 |
| sp\|P00918\|CAH2_HUMAN | AVQQPDGLAVLGIFLK.light | FALSE | 556.99429 | 577.370809 | 15.3 |
| sp\|P00918\|CAH2_HUMAN | AVQQPDGLAVLGIFLK.light | FALSE | 556.99429 | 690.454873 | 15.3 |
| sp\|P00918\|CAH2_HUMAN | AVQQPDGLAVLGIFLK.light | FALSE | 556.99429 | 789.523287 | 15.3 |
| sp\|P00918\|CAH2_HUMAN | AVQQPDGLAVLGIFLK.heavy | TRUE | 559.333345 | 584.387973 | 15.3 |
| sp\|P00918\|CAH2_HUMAN | AVQQPDGLAVLGIFLK.heavy | TRUE | 559.333345 | 697.472037 | 15.3 |
| sp\|P00918\|CAH2_HUMAN | AVQQPDGLAVLGIFLK.heavy | TRUE | 559.333345 | 796.540451 | 15.3 |
| sp\|Q9NZC2\|TREM2_HUMAN | VVSTHNLWLLSFLR.light | FALSE | 562.32263 | 635.387522 | 9.4 |
| sp\|Q9NZC2\|TREM2_HUMAN | VVSTHNLWLLSFLR.light | FALSE | 562.32263 | 748.471586 | 15.4 |
| sp\|Q9NZC2\|TREM2_HUMAN | VVSTHNLWLLSFLR.light | FALSE | 562.32263 | 751.409714 | 18.4 |
| sp\|Q9NZC2\|TREM2_HUMAN | VVSTHNLWLLSFLR.heavy | TRUE | 564.661684 | 642.404686 | 9.4 |
| sp\|Q9NZC2\|TREM2_HUMAN | VVSTHNLWLLSFLR.heavy | TRUE | 564.661684 | 751.409714 | 18.4 |
| sp\|Q9NZC2\|TREM2_HUMAN | VVSTHNLWLLSFLR.heavy | TRUE | 564.661684 | 755.48875 | 15.4 |
| sp\|O15240\|VGF_HUMAN | EPVAGDAVPGPK.light | FALSE | 568.800945 | 398.239795 | 21.6 |
| sp\|O15240\|VGF_HUMAN | EPVAGDAVPGPK.light | FALSE | 568.800945 | 740.39373 | 21.6 |
| sp\|O15240\|VGF_HUMAN | EPVAGDAVPGPK.light | FALSE | 568.800945 | 811.430844 | 18.6 |
| sp\|P04156\|PRIO_HUMAN | GENFTETDVK.light | FALSE | 570.264393 | 591.298432 | 15.7 |
| sp\|P04156\|PRIO_HUMAN | GENFTETDVK.light | FALSE | 570.264393 | 692.346111 | 18.7 |
| sp\|P04156\|PRIO_HUMAN | GENFTETDVK.light | FALSE | 570.264393 | 839.414525 | 15.7 |
| sp\|O15240\|VGF_HUMAN | EPVAGDAVPGPK.heavy | TRUE | 571.80785 | 398.239795 | 21.6 |
| sp\|O15240\|VGF_HUMAN | EPVAGDAVPGPK.heavy | TRUE | 571.80785 | 746.407539 | 21.6 |
| sp\|O15240\|VGF_HUMAN | EPVAGDAVPGPK.heavy | TRUE | 571.80785 | 817.444653 | 18.6 |
| sp\|P04156\|PRIO_HUMAN | GENFTETDVK.heavy | TRUE | 573.271297 | 597.312241 | 15.7 |
| sp\|P04156\|PRIO_HUMAN | GENFTETDVK.heavy | TRUE | 573.271297 | 698.35992 | 18.7 |
| sp\|P04156\|PRIO_HUMAN | GENFTETDVK.heavy | TRUE | 573.271297 | 845.428334 | 15.7 |
| sp\|Q15818\|NPTX1_HUMAN | LENLEQYSR.light | FALSE | 576.288202 | 682.315479 | 12.9 |
| sp\|Q15818\|NPTX1_HUMAN | LENLEQYSR.light | FALSE | 576.288202 | 795.399543 | 18.9 |
| sp\|Q15818\|NPTX1_HUMAN | LENLEQYSR.light | FALSE | 576.288202 | 909.442471 | 21.9 |

| Protein | Peptide | SIL | Precursor Ion | Product Ion | Collision Energy |
|---|---|---|---|---|---|
| sp\|P07602\|SAP_HUMAN | EIVDSYLPVILDIIK.light | FALSE | 577.337723 | 707.324647 | 13 |
| sp\|P07602\|SAP_HUMAN | EIVDSYLPVILDIIK.light | FALSE | 577.337723 | 714.476003 | 19 |
| sp\|P07602\|SAP_HUMAN | EIVDSYLPVILDIIK.light | FALSE | 577.337723 | 910.597181 | 13 |
| sp\|P07196\|NFL_HUMAN | VLEAELLVLR.light | FALSE | 577.860806 | 613.439558 | 24.9 |
| sp\|P07196\|NFL_HUMAN | VLEAELLVLR.light | FALSE | 577.860806 | 742.482151 | 15.9 |
| sp\|P07196\|NFL_HUMAN | VLEAELLVLR.light | FALSE | 577.860806 | 813.519265 | 27.9 |
| sp\|P07602\|SAP_HUMAN | EIVDSYLPVILDIIK.heavy | TRUE | 579.676777 | 707.324647 | 13 |
| sp\|P07602\|SAP_HUMAN | EIVDSYLPVILDIIK.heavy | TRUE | 579.676777 | 721.493167 | 19 |
| sp\|P07602\|SAP_HUMAN | EIVDSYLPVILDIIK.heavy | TRUE | 579.676777 | 917.614345 | 13 |
| sp\|Q15818\|NPTX1_HUMAN | LENLEQYSR.heavy | TRUE | 579.796784 | 682.315479 | 12.9 |
| sp\|Q15818\|NPTX1_HUMAN | LENLEQYSR.heavy | TRUE | 579.796784 | 802.416707 | 18.9 |
| sp\|Q15818\|NPTX1_HUMAN | LENLEQYSR.heavy | TRUE | 579.796784 | 916.459635 | 21.9 |
| sp\|P07196\|NFL_HUMAN | VLEAELLVLR.heavy | TRUE | 580.86771 | 619.453367 | 24.9 |
| sp\|P07196\|NFL_HUMAN | VLEAELLVLR.heavy | TRUE | 580.86771 | 748.49596 | 15.9 |
| sp\|P07196\|NFL_HUMAN | VLEAELLVLR.heavy | TRUE | 580.86771 | 819.533074 | 27.9 |
| sp\|P02745\|C1QA_HUMAN | GEQGEPGAPGIR.light | FALSE | 584.291276 | 442.277243 | 28.1 |
| sp\|P02745\|C1QA_HUMAN | GEQGEPGAPGIR.light | FALSE | 584.291276 | 501.193967 | 13.1 |
| sp\|P02745\|C1QA_HUMAN | GEQGEPGAPGIR.light | FALSE | 584.291276 | 667.388585 | 16.1 |
| sp\|P02745\|C1QA_HUMAN | GEQGEPGAPGIR.heavy | TRUE | 587.298181 | 448.291052 | 28.1 |
| sp\|P02745\|C1QA_HUMAN | GEQGEPGAPGIR.heavy | TRUE | 587.298181 | 501.193967 | 13.1 |
| sp\|P02745\|C1QA_HUMAN | GEQGEPGAPGIR.heavy | TRUE | 587.298181 | 673.402394 | 16.1 |
| sp\|P61981\|1433G_HUMAN | DSTLIMQLLR.light | FALSE | 595.334098 | 660.386142 | 16.5 |
| sp\|P61981\|1433G_HUMAN | DSTLIMQLLR.light | FALSE | 595.334098 | 773.470206 | 28.5 |
| sp\|P61981\|1433G_HUMAN | DSTLIMQLLR.light | FALSE | 595.334098 | 886.55427 | 16.5 |
| sp\|P61981\|1433G_HUMAN | DSTLIMQLLR.heavy | TRUE | 598.84268 | 667.403306 | 16.5 |
| sp\|P61981\|1433G_HUMAN | DSTLIMQLLR.heavy | TRUE | 598.84268 | 780.48737 | 28.5 |
| sp\|P61981\|1433G_HUMAN | DSTLIMQLLR.heavy | TRUE | 598.84268 | 893.571434 | 16.5 |
| sp\|P10645\|CMGA_HUMAN | SGELEQEEER.light | FALSE | 603.267664 | 690.305309 | 16.7 |
| sp\|P10645\|CMGA_HUMAN | SGELEQEEER.light | FALSE | 603.267664 | 819.347902 | 28.7 |
| sp\|P10645\|CMGA_HUMAN | SGELEQEEER.light | FALSE | 603.267664 | 932.431966 | 22.7 |
| sp\|P10645\|CMGA_HUMAN | SGELEQEEER.heavy | TRUE | 606.776246 | 690.305309 | 16.7 |
| sp\|P10645\|CMGA_HUMAN | SGELEQEEER.heavy | TRUE | 606.776246 | 819.347902 | 28.7 |
| sp\|P10645\|CMGA_HUMAN | SGELEQEEER.heavy | TRUE | 606.776246 | 939.44913 | 22.7 |
| sp\|P36222\|CH3L1_HUMAN | EAGTLAYYEIC[+57.021464]DFLR.light | FALSE | 607.622305 | 710.329021 | 11.1 |
| sp\|P36222\|CH3L1_HUMAN | EAGTLAYYEIC[+57.021464]DFLR.light | FALSE | 607.622305 | 823.413085 | 26.1 |
| sp\|P36222\|CH3L1_HUMAN | EAGTLAYYEIC[+57.021464]DFLR.light | FALSE | 607.622305 | 952.455678 | 26.1 |
| sp\|P36222\|CH3L1_HUMAN | EAGTLAYYEIC[+57.021464]DFLR.heavy | TRUE | 609.96136 | 717.346185 | 11.1 |

| Protein | Peptide | SIL | Precursor Ion | Product Ion | Collision Energy |
|---|---|---|---|---|---|
| sp\|P36222\|CH3L1_HUMAN | EAGTLAYYEIC[+57.021464]DFLR.heavy | TRUE | 609.96136 | 830.430249 | 26.1 |
| sp\|P36222\|CH3L1_HUMAN | EAGTLAYYEIC[+57.021464]DFLR.heavy | TRUE | 609.96136 | 959.472842 | 26.1 |
| sp\|P02745\|C1QA_HUMAN | VGYPGPSGPLGAR.light | FALSE | 614.327662 | 320.160482 | 14 |
| sp\|P02745\|C1QA_HUMAN | VGYPGPSGPLGAR.light | FALSE | 614.327662 | 811.442077 | 29 |
| sp\|P02745\|C1QA_HUMAN | VGYPGPSGPLGAR.light | FALSE | 614.327662 | 908.494841 | 23 |
| sp\|P09936\|UCHL1_HUMAN | MPFPVNHGASSEDTLLK.light | FALSE | 614.973338 | 376.168938 | 26.3 |
| sp\|P09936\|UCHL1_HUMAN | MPFPVNHGASSEDTLLK.light | FALSE | 614.973338 | 805.430175 | 17.3 |
| sp\|P09936\|UCHL1_HUMAN | MPFPVNHGASSEDTLLK.light | FALSE | 614.973338 | 892.462203 | 17.3 |
| sp\|P09936\|UCHL1_HUMAN | MPFPVNHGASSEDTLLK.heavy | TRUE | 617.312392 | 376.168938 | 26.3 |
| sp\|P09936\|UCHL1_HUMAN | MPFPVNHGASSEDTLLK.heavy | TRUE | 617.312392 | 812.447339 | 17.3 |
| sp\|P09936\|UCHL1_HUMAN | MPFPVNHGASSEDTLLK.heavy | TRUE | 617.312392 | 899.479367 | 17.3 |
| sp\|P01033\|TIMP1_HUMAN | GFQALGDAADIR.light | FALSE | 617.314751 | 660.33113 | 20.1 |
| sp\|P01033\|TIMP1_HUMAN | GFQALGDAADIR.light | FALSE | 617.314751 | 717.352593 | 17.1 |
| sp\|P01033\|TIMP1_HUMAN | GFQALGDAADIR.light | FALSE | 617.314751 | 830.436657 | 20.1 |
| sp\|P02745\|C1QA_HUMAN | VGYPGPSGPLGAR.heavy | TRUE | 617.836244 | 320.160482 | 14 |
| sp\|P02745\|C1QA_HUMAN | VGYPGPSGPLGAR.heavy | TRUE | 617.836244 | 818.459241 | 29 |
| sp\|P02745\|C1QA_HUMAN | VGYPGPSGPLGAR.heavy | TRUE | 617.836244 | 915.512005 | 23 |
| sp\|P01033\|TIMP1_HUMAN | GFQALGDAADIR.heavy | TRUE | 620.823333 | 660.33113 | 20.1 |
| sp\|P01033\|TIMP1_HUMAN | GFQALGDAADIR.heavy | TRUE | 620.823333 | 717.352593 | 17.1 |
| sp\|P01033\|TIMP1_HUMAN | GFQALGDAADIR.heavy | TRUE | 620.823333 | 837.453821 | 20.1 |
| sp\|P62258\|1433E_HUMAN | YLAEFATGNDR.light | FALSE | 628.798934 | 780.363492 | 20.5 |
| sp\|P62258\|1433E_HUMAN | YLAEFATGNDR.light | FALSE | 628.798934 | 909.406085 | 20.5 |
| sp\|P62258\|1433E_HUMAN | YLAEFATGNDR.light | FALSE | 628.798934 | 980.443199 | 20.5 |
| sp\|P62258\|1433E_HUMAN | YLAEFATGNDR.heavy | TRUE | 633.812548 | 790.39072 | 20.5 |
| sp\|P62258\|1433E_HUMAN | YLAEFATGNDR.heavy | TRUE | 633.812548 | 919.433313 | 20.5 |
| sp\|P62258\|1433E_HUMAN | YLAEFATGNDR.heavy | TRUE | 633.812548 | 990.470427 | 20.5 |
| sp\|O95502\|NPTXR_HUMAN | VAELEHGSSAYSPPDAFK.light | FALSE | 635.639012 | 577.298038 | 21.1 |
| sp\|O95502\|NPTXR_HUMAN | VAELEHGSSAYSPPDAFK.light | FALSE | 635.639012 | 674.350802 | 15.1 |
| sp\|O95502\|NPTXR_HUMAN | VAELEHGSSAYSPPDAFK.light | FALSE | 635.639012 | 761.382831 | 18.1 |
| sp\|O95502\|NPTXR_HUMAN | VAELEHGSSAYSPPDAFK.heavy | TRUE | 638.981421 | 587.325266 | 21.1 |
| sp\|O95502\|NPTXR_HUMAN | VAELEHGSSAYSPPDAFK.heavy | TRUE | 638.981421 | 684.37803 | 15.1 |
| sp\|O95502\|NPTXR_HUMAN | VAELEHGSSAYSPPDAFK.heavy | TRUE | 638.981421 | 771.410059 | 18.1 |
| sp\|P37840\|SYUA_HUMAN | EQVTNVGGAVVTGVTAVAQK.light | FALSE | 643.353094 | 716.430115 | 9.4 |
| sp\|P37840\|SYUA_HUMAN | EQVTNVGGAVVTGVTAVAQK.light | FALSE | 643.353094 | 773.451579 | 18.4 |
| sp\|P37840\|SYUA_HUMAN | EQVTNVGGAVVTGVTAVAQK.light | FALSE | 643.353094 | 874.499258 | 18.4 |
| sp\|P37840\|SYUA_HUMAN | EQVTNVGGAVVTGVTAVAQK.heavy | TRUE | 645.357697 | 722.443924 | 9.4 |
| sp\|P37840\|SYUA_HUMAN | EQVTNVGGAVVTGVTAVAQK.heavy | TRUE | 645.357697 | 779.465388 | 18.4 |

| Protein | Peptide | SIL | Precursor Ion | Product Ion | Collision Energy |
|---|---|---|---|---|---|
| sp\|P37840\|SYUA_HUMAN | EQVTNVGGAVVTGVTAVAQK.heavy | TRUE | 645.357697 | 880.513067 | 18.4 |
| sp\|P07360\|CO8G_HUMAN | VQEAHLTEDQIFYFPK.light | FALSE | 655.663269 | 678.35695 | 27.8 |
| sp\|P07360\|CO8G_HUMAN | VQEAHLTEDQIFYFPK.light | FALSE | 655.663269 | 701.365724 | 18.8 |
| sp\|P07360\|CO8G_HUMAN | VQEAHLTEDQIFYFPK.light | FALSE | 655.663269 | 814.449788 | 12.8 |
| sp\|P07602\|SAP_HUMAN | GC[+57.021464]SFLPDPYQK.light | FALSE | 656.305538 | 747.367181 | 15.3 |
| sp\|P07602\|SAP_HUMAN | GC[+57.021464]SFLPDPYQK.light | FALSE | 656.305538 | 860.451245 | 18.3 |
| sp\|P07602\|SAP_HUMAN | GC[+57.021464]SFLPDPYQK.light | FALSE | 656.305538 | 1094.55169 | 18.3 |
| sp\|P07360\|CO8G_HUMAN | VQEAHLTEDQIFYFPK.heavy | TRUE | 659.005678 | 678.35695 | 27.8 |
| sp\|P07360\|CO8G_HUMAN | VQEAHLTEDQIFYFPK.heavy | TRUE | 659.005678 | 711.392952 | 18.8 |
| sp\|P07360\|CO8G_HUMAN | VQEAHLTEDQIFYFPK.heavy | TRUE | 659.005678 | 824.477016 | 12.8 |
| sp\|P07602\|SAP_HUMAN | GC[+57.021464]SFLPDPYQK.heavy | TRUE | 659.312442 | 753.38099 | 15.3 |
| sp\|P07602\|SAP_HUMAN | GC[+57.021464]SFLPDPYQK.heavy | TRUE | 659.312442 | 866.465054 | 18.3 |
| sp\|P07602\|SAP_HUMAN | GC[+57.021464]SFLPDPYQK.heavy | TRUE | 659.312442 | 1100.5655 | 18.3 |
| sp\|P05067\|A4_HUMAN | LVFFAEDVGSNK.light | FALSE | 663.340435 | 748.347174 | 15.6 |
| sp\|P05067\|A4_HUMAN | LVFFAEDVGSNK.light | FALSE | 663.340435 | 819.384287 | 21.6 |
| sp\|P05067\|A4_HUMAN | LVFFAEDVGSNK.light | FALSE | 663.340435 | 966.452701 | 18.6 |
| sp\|Q92932\|PTPR2_HUMAN | AALGESGEQADGPK.light | FALSE | 665.317688 | 744.352259 | 21.6 |
| sp\|Q92932\|PTPR2_HUMAN | AALGESGEQADGPK.light | FALSE | 665.317688 | 801.373723 | 21.6 |
| sp\|Q92932\|PTPR2_HUMAN | AALGESGEQADGPK.light | FALSE | 665.317688 | 888.405751 | 21.6 |
| sp\|P05067\|A4_HUMAN | LVFFAEDVGSNK.heavy | TRUE | 666.347339 | 754.360983 | 15.6 |
| sp\|P05067\|A4_HUMAN | LVFFAEDVGSNK.heavy | TRUE | 666.347339 | 825.398096 | 21.6 |
| sp\|P05067\|A4_HUMAN | LVFFAEDVGSNK.heavy | TRUE | 666.347339 | 972.46651 | 18.6 |
| sp\|Q92932\|PTPR2_HUMAN | AALGESGEQADGPK.heavy | TRUE | 668.324592 | 750.366068 | 21.6 |
| sp\|Q92932\|PTPR2_HUMAN | AALGESGEQADGPK.heavy | TRUE | 668.324592 | 807.387532 | 21.6 |
| sp\|Q92932\|PTPR2_HUMAN | AALGESGEQADGPK.heavy | TRUE | 668.324592 | 894.41956 | 21.6 |
| sp\|Q9NZC2\|TREM2_HUMAN | NLQPHDAGLYQC[+57.021464]QSLHGSEADTLR.light | FALSE | 678.321408 | 356.192845 | 28.6 |
| sp\|Q9NZC2\|TREM2_HUMAN | NLQPHDAGLYQC[+57.021464]QSLHGSEADTLR.light | FALSE | 678.321408 | 704.357344 | 28.6 |
| sp\|Q9NZC2\|TREM2_HUMAN | NLQPHDAGLYQC[+57.021464]QSLHGSEADTLR.light | FALSE | 678.321408 | 848.410836 | 25.6 |
| sp\|Q9NZC2\|TREM2_HUMAN | NLQPHDAGLYQC[+57.021464]QSLHGSEADTLR.heavy | TRUE | 680.075699 | 356.192845 | 28.6 |
| sp\|Q9NZC2\|TREM2_HUMAN | NLQPHDAGLYQC[+57.021464]QSLHGSEADTLR.heavy | TRUE | 680.075699 | 711.374508 | 28.6 |
| sp\|Q9NZC2\|TREM2_HUMAN | NLQPHDAGLYQC[+57.021464]QSLHGSEADTLR.heavy | TRUE | 680.075699 | 855.428 | 25.6 |
| sp\|P04114\|APOB_HUMAN | ILGEELGFASLHDLQLLGK.light | FALSE | 685.049123 | 786.47198 | 19.9 |
| sp\|P04114\|APOB_HUMAN | ILGEELGFASLHDLQLLGK.light | FALSE | 685.049123 | 923.530892 | 19.9 |
| sp\|P04114\|APOB_HUMAN | ILGEELGFASLHDLQLLGK.light | FALSE | 685.049123 | 1036.61496 | 19.9 |
| sp\|P04114\|APOB_HUMAN | ILGEELGFASLHDLQLLGK.heavy | TRUE | 688.391533 | 786.47198 | 19.9 |
| sp\|P04114\|APOB_HUMAN | ILGEELGFASLHDLQLLGK.heavy | TRUE | 688.391533 | 923.530892 | 19.9 |
| sp\|P04114\|APOB_HUMAN | ILGEELGFASLHDLQLLGK.heavy | TRUE | 688.391533 | 1036.61496 | 19.9 |

| Protein | Peptide | SIL | Precursor Ion | Product Ion | Collision Energy |
|---|---|---|---|---|---|
| sp\|P28799\|GRN_HUMAN | EVVSAQPATFLAR.light | FALSE | 694.880258 | 775.4461 | 16.5 |
| sp\|P28799\|GRN_HUMAN | EVVSAQPATFLAR.light | FALSE | 694.880258 | 903.504677 | 16.5 |
| sp\|P28799\|GRN_HUMAN | EVVSAQPATFLAR.light | FALSE | 694.880258 | 974.541791 | 25.5 |
| sp\|P10909\|CLUS_HUMAN | ASSIIDELFQDR.light | FALSE | 697.351531 | 807.399543 | 25.6 |
| sp\|P10909\|CLUS_HUMAN | ASSIIDELFQDR.light | FALSE | 697.351531 | 922.426487 | 31.6 |
| sp\|P10909\|CLUS_HUMAN | ASSIIDELFQDR.light | FALSE | 697.351531 | 1035.51055 | 22.6 |
| sp\|P28799\|GRN_HUMAN | EVVSAQPATFLAR.heavy | TRUE | 699.893872 | 785.473328 | 16.5 |
| sp\|P28799\|GRN_HUMAN | EVVSAQPATFLAR.heavy | TRUE | 699.893872 | 913.531905 | 16.5 |
| sp\|P28799\|GRN_HUMAN | EVVSAQPATFLAR.heavy | TRUE | 699.893872 | 984.569019 | 25.5 |
| sp\|P10909\|CLUS_HUMAN | ASSIIDELFQDR.heavy | TRUE | 700.860113 | 814.416707 | 25.6 |
| sp\|P10909\|CLUS_HUMAN | ASSIIDELFQDR.heavy | TRUE | 700.860113 | 929.443651 | 31.6 |
| sp\|P10909\|CLUS_HUMAN | ASSIIDELFQDR.heavy | TRUE | 700.860113 | 1042.52772 | 22.6 |
| sp\|P61981\|1433G_HUMAN | TAFDDAIAELDTLNEDSYK.light | FALSE | 710.995301 | 734.335546 | 29.8 |
| sp\|P61981\|1433G_HUMAN | TAFDDAIAELDTLNEDSYK.light | FALSE | 710.995301 | 755.320624 | 23.8 |
| sp\|P61981\|1433G_HUMAN | TAFDDAIAELDTLNEDSYK.light | FALSE | 710.995301 | 969.452367 | 29.8 |
| sp\|P63104\|1433Z_HUMAN | TAFDEAIAELDTLSEESYK.light | FALSE | 711.335435 | 742.325375 | 17.8 |
| sp\|P63104\|1433Z_HUMAN | TAFDEAIAELDTLSEESYK.light | FALSE | 711.335435 | 748.351196 | 23.8 |
| sp\|P63104\|1433Z_HUMAN | TAFDEAIAELDTLSEESYK.light | FALSE | 711.335435 | 855.409439 | 29.8 |
| sp\|P63104\|1433Z_HUMAN | TAFDEAIAELDTLSEESYK.heavy | TRUE | 713.674489 | 742.325375 | 17.8 |
| sp\|P63104\|1433Z_HUMAN | TAFDEAIAELDTLSEESYK.heavy | TRUE | 713.674489 | 748.351196 | 23.8 |
| sp\|P63104\|1433Z_HUMAN | TAFDEAIAELDTLSEESYK.heavy | TRUE | 713.674489 | 862.426603 | 29.8 |
| sp\|P61981\|1433G_HUMAN | TAFDDAIAELDTLNEDSYK.heavy | TRUE | 714.33771 | 734.335546 | 29.8 |
| sp\|P61981\|1433G_HUMAN | TAFDDAIAELDTLNEDSYK.heavy | TRUE | 714.33771 | 765.347852 | 23.8 |
| sp\|P61981\|1433G_HUMAN | TAFDDAIAELDTLNEDSYK.heavy | TRUE | 714.33771 | 979.479595 | 29.8 |
| sp\|P10645\|CMGA_HUMAN | RPEDQELESLSAIEAELEK.light | FALSE | 729.365617 | 831.445825 | 12.5 |
| sp\|P10645\|CMGA_HUMAN | RPEDQELESLSAIEAELEK.light | FALSE | 729.365617 | 989.514967 | 30.5 |
| sp\|P10645\|CMGA_HUMAN | RPEDQELESLSAIEAELEK.light | FALSE | 729.365617 | 997.458515 | 30.5 |
| sp\|P10645\|CMGA_HUMAN | RPEDQELESLSAIEAELEK.heavy | TRUE | 731.704672 | 838.462989 | 12.5 |
| sp\|P10645\|CMGA_HUMAN | RPEDQELESLSAIEAELEK.heavy | TRUE | 731.704672 | 996.532131 | 30.5 |
| sp\|P10645\|CMGA_HUMAN | RPEDQELESLSAIEAELEK.heavy | TRUE | 731.704672 | 997.458515 | 30.5 |
| sp\|P62258\|1433E_HUMAN | LIC[+57.021464]C[+57.021464]DILDVLDK.light | FALSE | 738.87547 | 815.487296 | 26.9 |
| sp\|P62258\|1433E_HUMAN | LIC[+57.021464]C[+57.021464]DILDVLDK.light | FALSE | 738.87547 | 930.514239 | 26.9 |
| sp\|P62258\|1433E_HUMAN | LIC[+57.021464]C[+57.021464]DILDVLDK.light | FALSE | 738.87547 | 1090.54489 | 26.9 |
| sp\|P62258\|1433E_HUMAN | LIC[+57.021464]C[+57.021464]DILDVLDK.heavy | TRUE | 741.882374 | 821.501105 | 26.9 |
| sp\|P62258\|1433E_HUMAN | LIC[+57.021464]C[+57.021464]DILDVLDK.heavy | TRUE | 741.882374 | 936.528048 | 26.9 |
| sp\|P62258\|1433E_HUMAN | LIC[+57.021464]C[+57.021464]DILDVLDK.heavy | TRUE | 741.882374 | 1096.5587 | 26.9 |
| sp\|P36222\|CH3L1_HUMAN | TLLSVGGWNFGSQR.light | FALSE | 761.394064 | 894.421676 | 18.6 |

| Protein | Peptide | SIL | Precursor Ion | Product Ion | Collision Energy |
|---|---|---|---|---|---|
| sp\|P36222\|CH3L1_HUMAN | TLLSVGGWNFGSQR.light | FALSE | 761.394064 | 951.44314 | 21.6 |
| sp\|P36222\|CH3L1_HUMAN | TLLSVGGWNFGSQR.light | FALSE | 761.394064 | 1008.4646 | 27.6 |
| sp\|P36222\|CH3L1_HUMAN | TLLSVGGWNFGSQR.heavy | TRUE | 766.407678 | 904.448904 | 18.6 |
| sp\|P36222\|CH3L1_HUMAN | TLLSVGGWNFGSQR.heavy | TRUE | 766.407678 | 961.470368 | 21.6 |
| sp\|P36222\|CH3L1_HUMAN | TLLSVGGWNFGSQR.heavy | TRUE | 766.407678 | 1018.49183 | 27.6 |
